# Convergent evolution of a novel blood-red nectar pigment in vertebrate-pollinated flowers

**DOI:** 10.1101/2021.03.24.436888

**Authors:** Rahul Roy, Nickolas Moreno, Stephen A. Brockman, Adam Kostanecki, Amod Zambre, Catherine Holl, Erik M. Solhaug, Anzu Minami, Emilie Snell-Rood, Marshall Hampton, Mark A. Bee, Ylenia Chiari, Adrian D. Hegeman, Clay J. Carter

## Abstract

*Nesocodon mauritianus* (Campanulaceae) produces a blood-red nectar that has been proposed to serve as a visual attractant for pollinator visitation. Here we show that the nectar’s red color is derived from a novel alkaloid termed nesocodin. The first nectar produced is acidic and pale yellow in color, but slowly becomes alkaline before taking on its characteristic red color. Three enzymes secreted into the nectar are either necessary or sufficient for pigment production, including (1) a carbonic anhydrase that creates an alkaline environment, (2) an aryl alcohol oxidase that generates sinapaldehyde, a pigment precursor, and (3) a ferritin-like catalase that protects nesocodin from degradation by hydrogen peroxide. Our findings demonstrate how these three enzymatic activities allow for the condensation of sinapaldehyde and proline to form a novel pigment with a stable imine bond, which in turn is attractive to *Phelsuma* geckos, the presumed pollinators of *Nesocodon*. We also identify nesocodin in the red nectar of the distantly related *Jaltomata herrerae* and provide evidence for convergent evolution of this trait. While the overall enzymatic activities required for red pigment formation in both *Nesocodon* and *J. herrerae* nectars are identical, the associated genes encoding the enzymes are not orthologous and, in the case of the aryl alcohol oxidase, even belong to different protein families. This work cumulatively identifies a novel, convergently evolved trait in two vertebrate-pollinated species, suggesting the red pigment is selectively favored and that only a limited number of compounds are likely to underlie this adaptation.

**Significance:** Nearly 90% of flowering plants produce nectar to attract pollinators. Beyond sugars, many types of nectar solutes play important ecological roles; however, the molecular basis for the diversity of nectar composition across species is less explored. One rare trait among flowering plants is the production of colored nectar, which may function to attract and guide prospective pollinators. Our findings indicate convergent evolution of a red-colored nectar across two distantly related plant species. Behavioral data show that the red pigment attracts diurnal geckos, a presumed pollinator of one of these plants. These findings join a growing list of examples of distinct biochemical and molecular mechanisms underlying evolutionary convergence, and provide a fascinating system for testing how interactions across species drive the evolution of novel pigments in an understudied context.

## MAIN TEXT

Pigments mediate essential physiological and ecological functions throughout all domains of life, from roles in UV protection in both eukaryotes and prokaryotes (1), to light-regulated development and establishment of circadian rhythms (2), to visual signaling and mate choice in animals (3). Among plants, flower color is often crucial in pollinator attraction (4). Plant-pollinator interactions have driven at least a portion of the massive species radiation observed in the angiosperms, leading not only to extremely diverse floral size and morphology (5), but also the accompanying chemistries behind attractants (color and scent) and rewards (nectar and pollen) (e.g. (6–9)). Although rare, one floral trait that has evolved multiple times is colored nectar (10), which has been suggested to serve as an ‘honest’ visual cue of a reward to prospective pollinators (10, 11). Approximately 70 plant species are known to produce colored nectars (10), but the identities and biosyntheses of these pigments, as well as their associated functions, are largely unknown. One charismatic example is the blood-red nectar of *Nesocodon mauritianus* (12), which appears to play a role in the attraction of day geckos (*Phelsuma* spp.) (13). This study reports an investigation into the biochemical nature and biological function of this red-pigmented nectar within a phylogenetic and evolutionary framework to address our gap in knowledge of colored nectars.

### Nectar color with respect to floral phenology

*Nesocodon* nectar is well known to have a deep-red color, but a phenological examination of nectar production has not been reported. Toward this end, we found that newly opened *Nesocodon* flowers (0 h) actually have a yellow nectar (Fig. 1A-B, fig. S1), which gradually turns orange (Fig. 1E) and then red (Fig. 1C-E). The progression from yellow-to-orange-to-red positively corresponds to nectar alkalinization (Fig. 1E). Artificial alkalinization of yellow nectar (0 h) shifts the UV/visible absorbance maximum (λ_MAX_) from 343 nm to 430 nm, but does not yield a red color (fig. S2A). Conversely, step-wise acidification of red nectar (+24 h) yields a yellow color by proportionally decreasing the major absorbance peak at 505 nm, with a concomitant increase in a peak present in the yellow (0 h) nectar at ~343 nm and a separate peak at 403 nm (fig. S2B). The red color rapidly returns with re-alkalination of the artificially acidified +24 h nectar with a λ_MAX_ of 505 nm. These results cumulatively indicate that the red pigment λ_MAX_ of 505 nm under alkaline conditions shifts to 403 nm upon acidification.

**Fig. 1.**
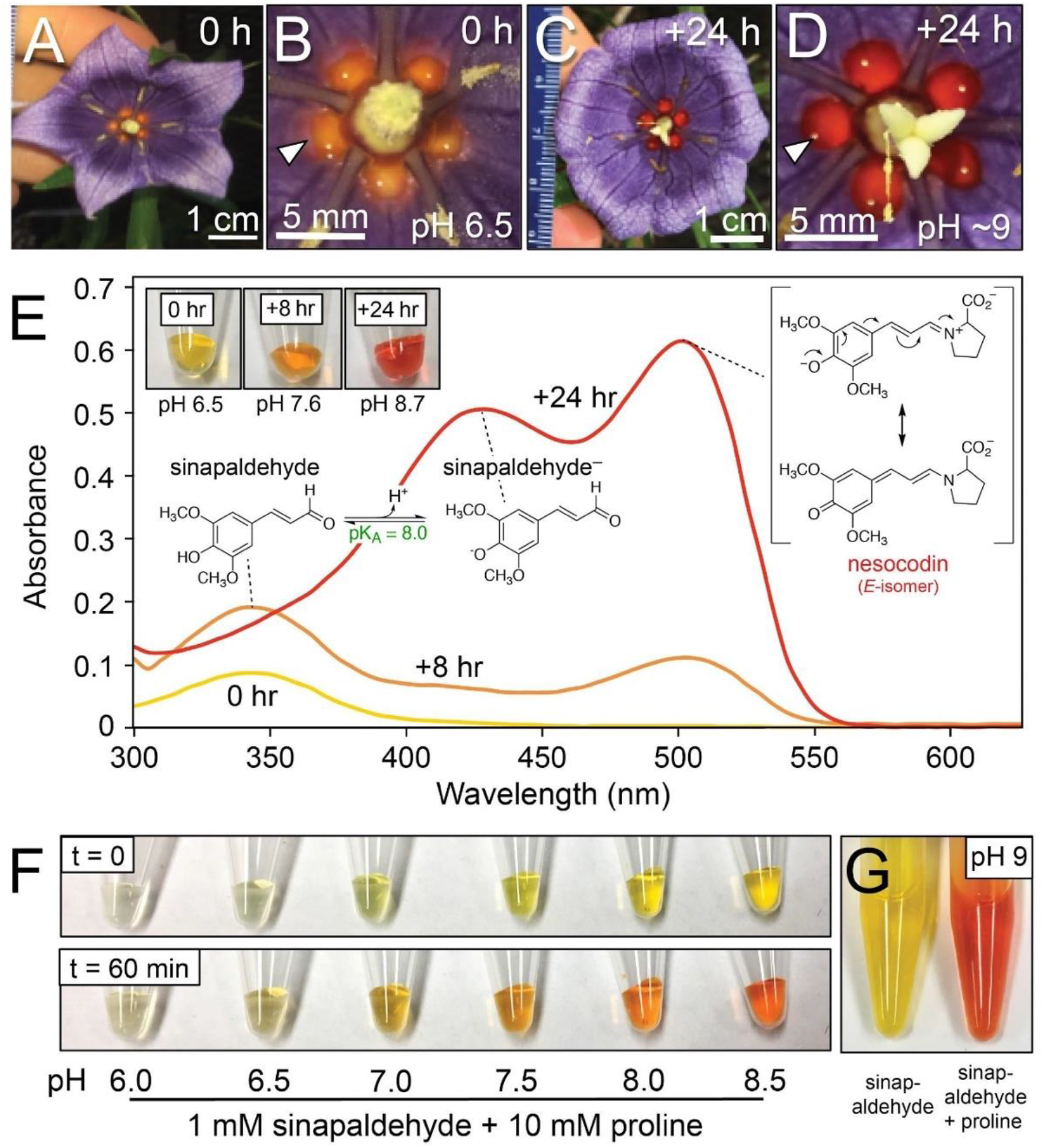
The color and pH of *N. mauritianus* nectar change over time. **(A and B)** Newly opened flower (0 h); **B** is a close-up of **A** showing the color of five distinct nectar droplets (arrowhead). **(C and D)** Flower ~24 hours after opening; **D** is a close-up of **C**. **(E)** UV/visible absorbance spectra of nectars collected at 0 h (yellow), +8 h (orange) and +24 h (red) after dilution at 1:50 in deionized H_2_O. Images of each undiluted nectar and pH are noted in the inset. Structures of the pigments contributing to the absorbance peaks were initially identified by LC-MS (fig. S3, S4, S9) and confirmed by multiple approaches (fig. S7, S8, S10-14). **(F)** pH-dependent formation of colored products derived from 1 mM sinapaldehyde and 10 mM proline in 25 mM buffers of varying pH, including 6.0 (MES), 6.5 (MES), 7.0 (HEPES), 7.5 (HEPES), 8.0 (HEPES), and 8.5 (Tricine). **(G)** Color of 1 mM sinapaldehyde in 50 mM Na(H)CO_3_, pH 9.0, with and without 10 mM L-proline after a 30 min incubation at 21°C.

### Pigment identification

Using these pH-dependent absorbance maxima as key indicators, LC-MS & MS/MS analyses of both yellow (0 h) and red (+24 h) nectars under acidic mobile conditions identified an elution peak with an *m/z* of 209.0807(±0.0004) and λ_MAX_ of ~343 nm (fig. S3). This compound was identified as sinapaldehyde [(E)-3,5-dimethoxy-4-hydroxycinnamaldehyde]) by fragmentation analysis and comparison of chromatographic and UV/visible spectroscopic properties to a commercially obtained standard (fig. S3, S4).

LC-MS/MS analysis of red nectar (+24 h) identified another major peak with an *m/z* of 306.1333(±0.0006) and λ_MAX_ of 403 nm matching that of acidified red nectar (fig. S2B, S3). Similarities in MS/MS fragmentation suggested that the red pigment results from conjugation of sinapaldehyde with another nectar constituent. The exact mass difference between the conjugate and sinapaldehyde, plus the mass of water of hydrolysis is 115.0632, which matches the elemental composition of the amino acid proline (C_5_H_9_NO_2_). Like the nectars of several distantly related species, including those of tobacco and soybean (14), we found that free proline accumulates to high levels in *Nesocodon* nectar (up to ~60 mM; e.g., fig. S6). We postulated that the red pigment is an imine conjugate of proline and sinapaldehyde, the formation of which depends on the deprotonation of proline such that the equilibrium shifts toward the formation of the red conjugate with increasing pH (fig. S5A). Indeed, aqueous mixtures of sinapaldehyde and proline produced a pH-dependent, red-colored compound *in vitro*, hereafter referred to as ‘nesocodin’ (Fig. 1F-G, fig. S5B). Reduction of nesocodin using sodium borohydride (NaBH_4_)or sodium borodeuteride (NaBD_4_) resulted in both a loss of color and accumulation of the expected products of imine reduction (fig. S7). Spectroscopic, spectrometric and chromatographic properties of synthetic nesocodin were identical to those of the natural pigment (fig. S8, S9). Anhydrous synthetic reaction conditions were then devised and provided quantitative conversion of L-proline and sinapaldehyde to nesocodin, which was structurally characterized by NMR (fig. S10-14) and found to form as a mixture of *E*- and *Z*-isomers (~2:1). Similar separation of these isomers by UHPLC analysis in both synthetic and nectar-derived nesocodin (fig. S15) suggests that imine conjugation *in vivo* is non-enzymatic.

The pH-dependence of nesocodin production led to several conclusions: (1) nectar alkalinization is essential for proline deprotonation to shift the equilibrium towards nesocodin formation (fig. S5); (2) the absorbance spectra of both nesocodin and sinapaldehyde display bathochromic shifts as these compounds are deprotonated (pK_A_s of 6.5 and 8.0 respectively, Fig. S5, S16), which is sufficient for the blood-red coloration of mature nectar; and (3) nesocodin is likely stabilized by resonance delocalization (Fig. 1E) upon deprotonation, a phenomenon expected to positively affect it’s accumulation in aqueous systems (15).

### Pigment biosynthesis

Since nesocodin formation and stability requires alkaline conditions, we next sought factors that increase nectar pH. Three major proteins were identified in *Nesocodon* nectar: a putative α-type carbonic anhydrase (NmNec1, 28 kDa), a desiccation-related protein (NmNec2, 33 kDa) and a mandelonitrile lyase-like protein (NmNec3, 65 kDa) (Fig. 2A; fig. S17, S18). *NmNec1* and *NmNec3* display enriched expression in nectaries, whereas *NmNec2* appears to be ubiquitously expressed in flowers (fig. S17D).

**Fig. 2.**
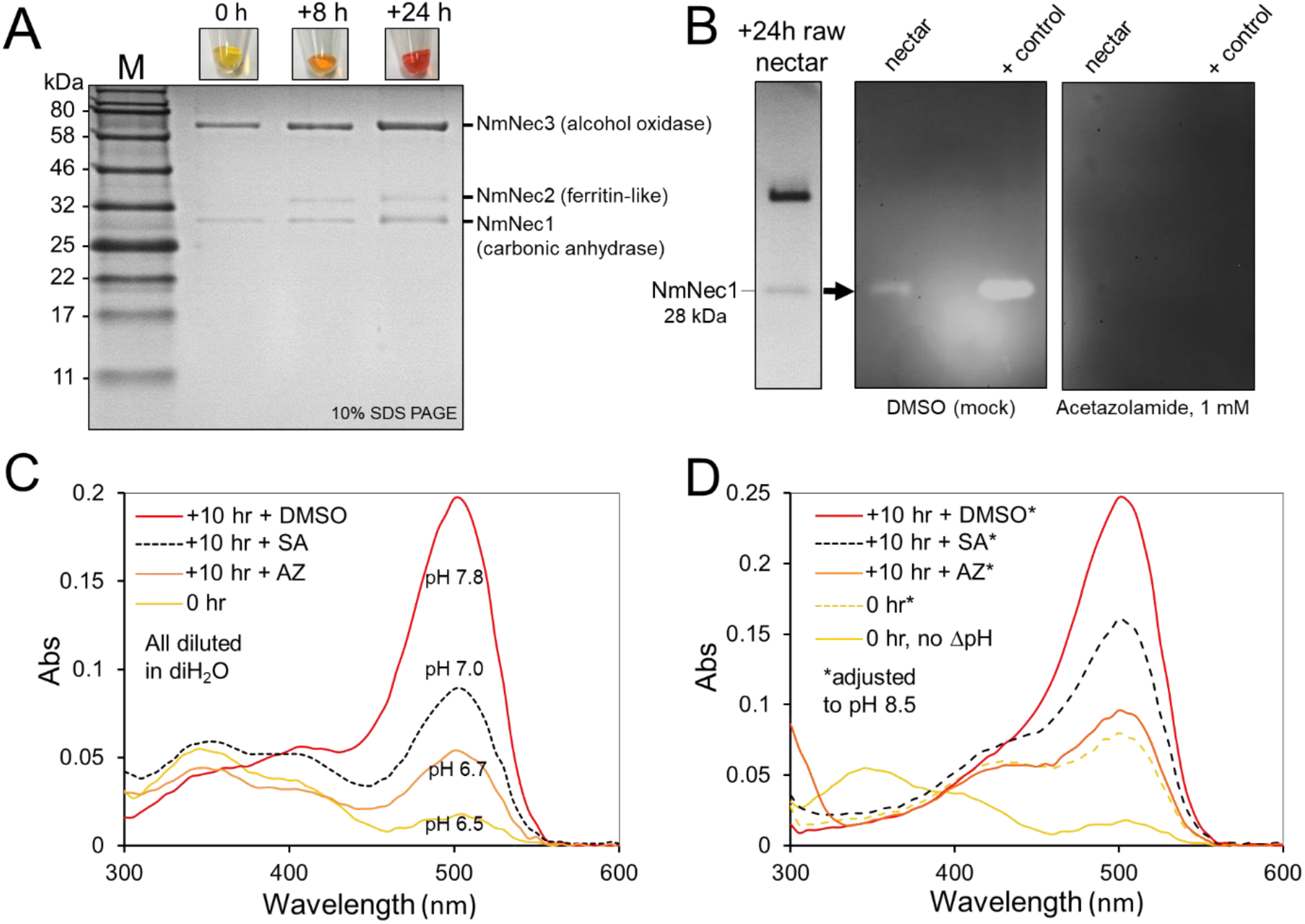
A carbonic anhydrase is responsible for nectar alkalinization. **(A)** SDS PAGE analysis of 0 h, +8 h and +24 h nectars (18 μL each; see fig. S17 for protein identification). **(B)** In-gel carbonic anhydrase assay of 18 μL of +24 h nectar and a positive control without (left) and with (right) 1 mM acetazolamide added. **(C)** UV/visible absorbance spectra and pH of nectar droplets treated *in situ* with carbonic anhydrase inhibitors. The carbonic anhydrase inhibitors sulfanilamide (SA) or acetazolamide (AZ) were added to separate 0 h nectar droplets in the same flower to a final concentration of 1 mM; a mock treatment containing an equal volume of 10% DMSO was included; one untreated nectar droplet was collected at 0 h and stored at 4°C in the dark; the treated nectar droplets were subsequently collected at +10 h, measured for pH, diluted 1:50 in H_2_O and evaluated by UV-Vis spectrophotometry. **(D)** Absorbance spectra of the same nectars from panel **C** after diluting 1:50 with 50 mM Tricine, pH 8.5. Additional representative results from individual treated flowers are shown in fig. S20.

Carbonic anhydrases help regulate both extracellular and cytosolic pH in animals, including cases of extreme alkalinization (16), thus we predicted NmNec1 plays a role in increasing nectar pH. NmNec1 has carbonic anhydrase activity that is inhibitable with either sulfanilamide or acetazolamide (e.g. Fig. 2B; fig. S19A). Adding bicarbonate to 0 h nectar increased nectar pH, whereas buffers of equivalent molarity and pH did not (fig. S19B). Similarly, removal of yellow nectar (0 h) from flowers prevented further alkalinization (fig. S19C), suggesting the nectary supplies nectar with bicarbonate. The nectar droplets on *Nesocodon* flowers remain physically separated until a few days after opening (Fig. 1A). We took advantage of this characteristic by adding carbonic anhydrase inhibitors directly to individual nectar droplets *in situ*. Addition of either sulfanilamide, acetazolamide or 6-ethoxy-2-benzothiazolesulfonamide decreased both nectar alkalinization and absorbance at 505 nm (Fig. 2C, fig. S20). The same treated nectar droplets showed less red pigment formation relative to control droplets when adjusted to alkaline pH (Fig. 2D), confirming that little nesocodin is produced under acidic-to-neutral conditions (fig. S5B).

Given the acidic pH of most intracellular compartments, it is unlikely that nesocodin could be made intracellularly and then secreted, so we next questioned whether other nectar proteins mediate pigment formation. NmNec3 belongs to the GMC flavoenzyme oxidoreductase family (fig. S21), as determined by the NCBI Conserved Domain Search tool. Purified NmNec3 (Fig. 3A) was verified as a flavoprotein (fig. S22A) with alcohol oxidase activity (fig. S22B), including toward sinapyl alcohol to produce sinapaldehyde (Fig. 3A). Moreover, addition of proline to the reaction drove nesocodin formation (Fig. 3B-D). Sinapyl alcohol, the precursor to sinapaldehyde, was also detected in nectar at very low levels (fig. S23), suggesting it is exported from nectary cells prior to oxidation. Unsurprisingly, transcripts encoding enzymes for the phenylpropanoid pathway from phenylalanine to sinapyl alcohol were highly expressed in nectaries (fig. S24).

**Fig. 3.**
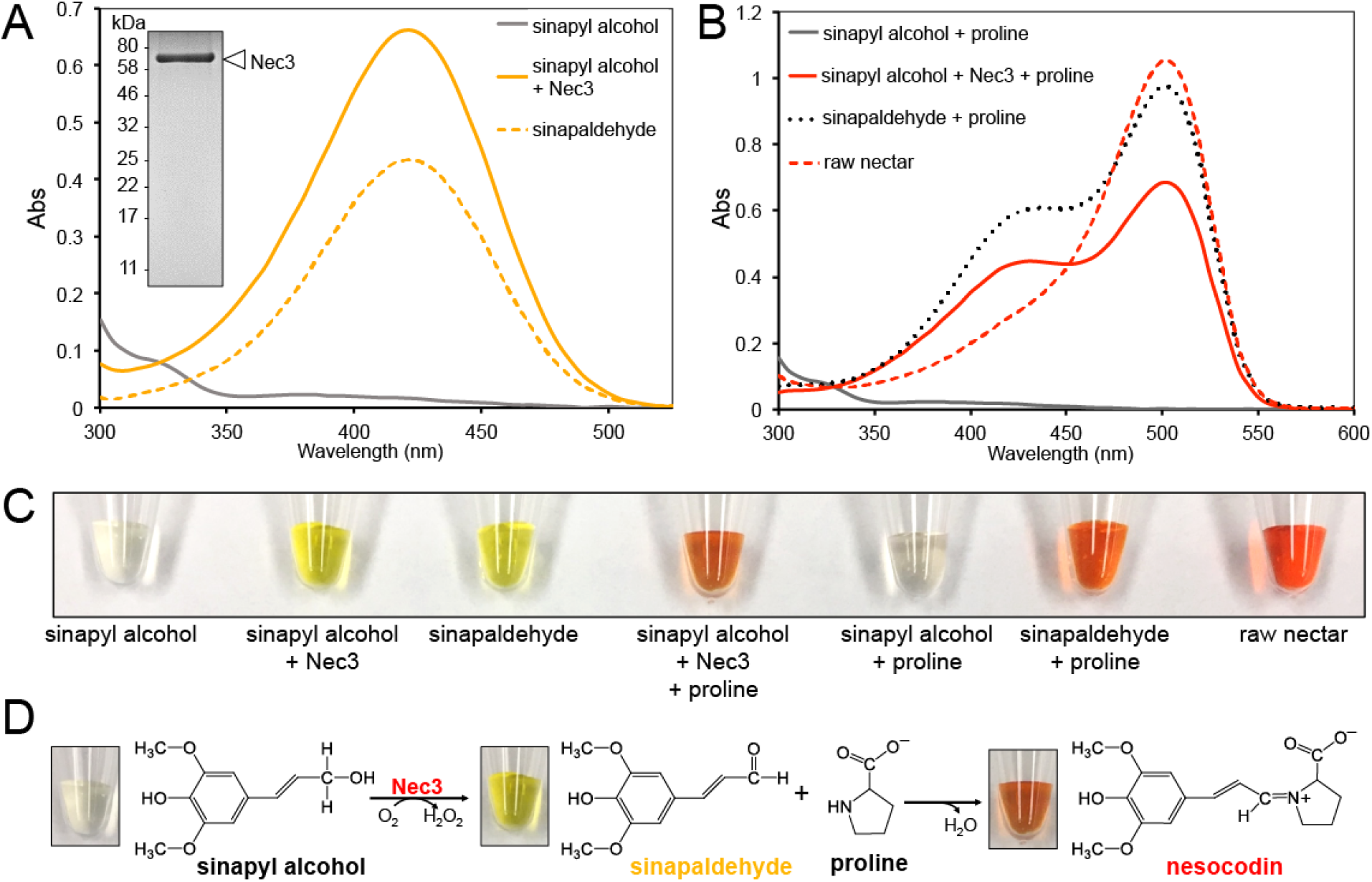
NmNec3 can oxidize sinapyl alcohol into sinapaldehyde to serve as a precursor to nesocodin. **(A)** Purified NmNec3 (inset) has sinapyl alcohol oxidase activity and can produce sinapaldehyde. The absorbance spectrum shown by solid yellow line is the result of a reaction mix containing 1 mM sinapyl alcohol and 0.1 μg/mL NmNec3 in 50 mM Na(H)CO_3_, pH 9.0. Negative controls with sinapyl alcohol but no NmNec3 did not yield any sinapaldehyde (grey line); a 0.5 mM sinapaldehyde standard in 50 mM Na(H)CO_3_, pH 9.0 was used as a reference. **(B)** The same reactions as in panel **A** (with NmNec3), but also containing 10 mM proline produced a red colored product (solid red line) consistent with both synthetic nesocodin (black dotted line; 1 mM sinapaldehyde plus 10 mM proline) and raw nectar (red dashed line; diluted 1:20). **(C)** Representative images showing the colors of the products of panels **A** and **B**. **(D)** Reaction sequence between sinapyl alcohol, NmNec3 and proline to yield nesocodin.

NmNec2 has high similarity to plant ‘desiccation-related proteins’ (fig. S25). Though with unknown function, these proteins contain a ferritin-like domain, with a subgroup of related bacterial proteins being Mn-dependent catalases (17). Catalase dismutates hydrogen peroxide (H_2_O_2_) into water and O_2_(g). We first determined that fresh, raw *Nesocodon* nectar has no detectable H_2_O_2_, suggesting the possibility of active removal, especially since one of the products from NmNec3’s alcohol oxidase activity is H_2_O_2_ (fig. S3D). Subsequent in-gel (Fig. 4A) and proteomic analysis (fig. S26A) demonstrated that non-boiled NmNec2 is a multimeric catalase (~90 kDa), which dissociates into monomers upon boiling (~30 kDa) (Fig. 4A), consistent with the detergent-resistant, multimeric ferritin-like Mn catalases found in some bacteria (17). Similarly, purified total nectar proteins have catalase activity that is destroyed by boiling (Fig. 4B, fig. S26B).

**Fig. 4.**
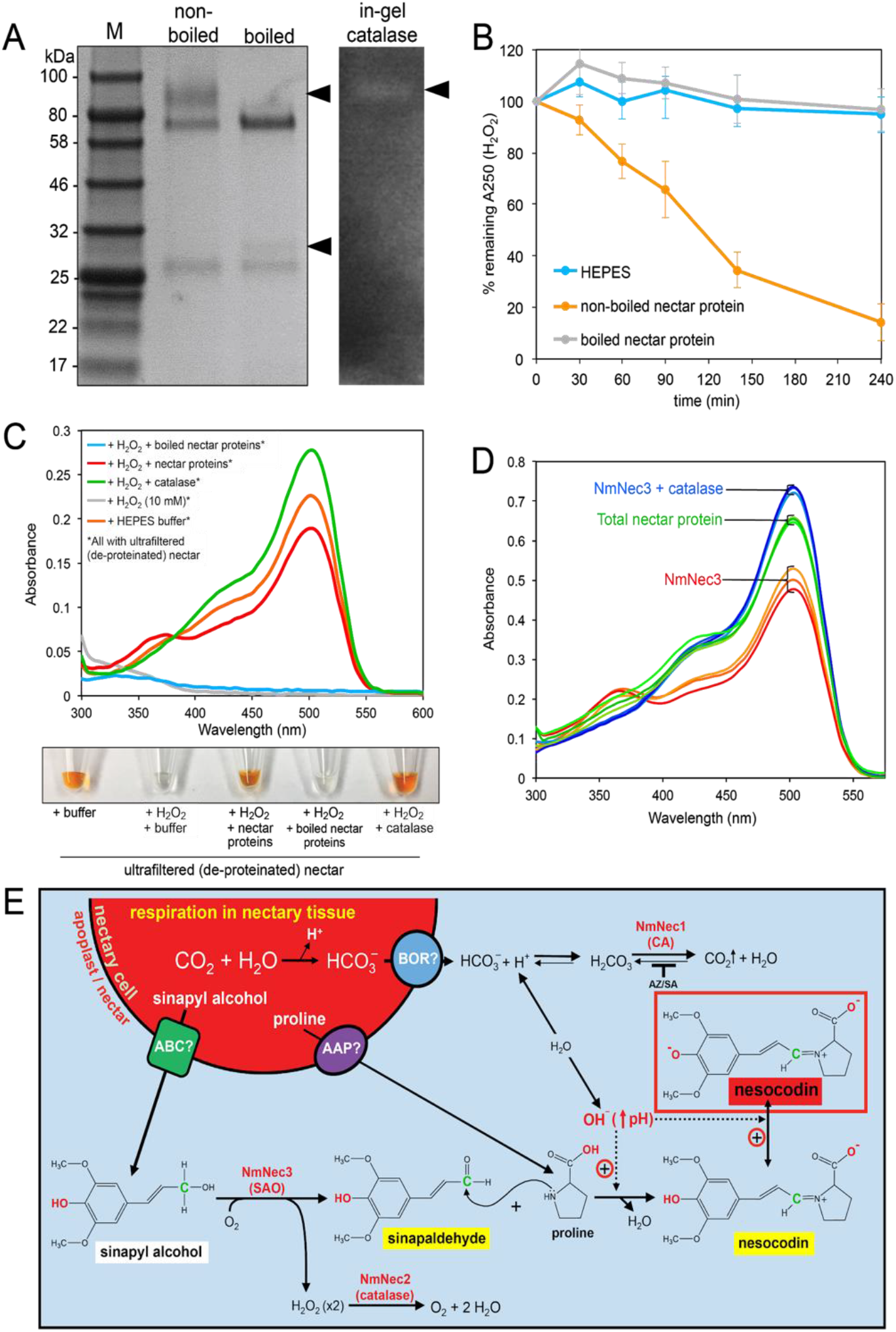
NmNec2 has catalase activity that can protect nesocodin from H_2_O_2_-mediated degradation. **(A)** Left, SDS PAGE (4-20%) analysis of 18 μL of raw non-boiled and boiled nectar stained with Coomassie Blue; right, in-gel catalase assay after SDS PAGE without sample boiling or addition of reducing agent. Arrowheads indicate location of NmNec3 and associated activity band. **(B)** Catalase activity of total non-boiled (orange line) or boiled (grey) nectar proteins. Assays contained 25 mM H_2_O_2_ in 50 mM HEPES pH 8.0 and either 0.17 μg/μL of total nectar proteins (either boiled or non-boiled) or an equivalent volume of HEPES buffer as a negative control. **(C)** Catalase-mediated protection of natural nesocodin. Filtered (de-proteinated) nectar was incubated with 10 mM H_2_O_2_ for 18 h with either total non-boiled (yellow line) or boiled (blue) nectar proteins (0.1 μg/mL final). Controls included the addition of heme-based catalase (green), no protein (grey), and no H_2_O_2_ added [orange, ‘+HEPES’ (25 mM pH 8.0)]. **(D)** Catalase-mediated protection of quasi-synthetic nesocodin. All reactions contained 1 mM sinapyl alcohol, 10 mM proline and 0.1 μg/μL NmNec3 in 50 mM Na(H)CO_3_, pH 9.0. A subset of reactions contained either 0.2 μg/μL total nectar protein or 0.1 μg/μL of heme-based catalase. Traces of absorbance spectra from three individual replicates are presented as labeled. **(E)** Model for *in vivo* nesocodin synthesis.

To determine a role for catalase activity in nectar, exogenous H_2_O_2_ was applied to either ultrafiltered (de-proteinated) +24 h nectar or synthetic nesocodin, which degraded the pigment in both cases (Fig. 4C, fig. S26). Conversely, addition of purified nectar proteins (non-boiled) or commercial heme-based catalase protected the pigment (Fig. 4C). While the sinapyl alcohol oxidase activity of NmNec3 may be essential for red pigment formation, H_2_O_2_ produced by this reaction may limit final nectar color intensity. Indeed, addition of purified nectar proteins or heme-based catalase to reactions containing NmNec3, sinapyl alcohol, and proline increased total pigment yield (Fig. 4D).

We propose a mechanistic model for nectar pigment synthesis and stability in *Nesocodon* (Fig. 4E). Nectar from newly opened flowers is weakly acidic and pale yellow, but its pH progressively increases through the action of a carbonic anhydrase (NmNec1), likely supplied with bicarbonate through nectary respiration. Simultaneously, sinapyl alcohol is exported from nectaries and oxidized to sinapaldehyde by the alcohol oxidase NmNec3. Nectar alkalinization supports (1) condensation of sinapaldehyde with proline to form nesocodin, and (2) the deprotonation of both sinapaldehyde and nesocodin, which is required for color formation and stability. Lastly, NmNec2 has catalase activity that protects nesocodin from degradation by the H_2_O_2_ generated by NmNec3.

### A functional evaluation of *Nesocodon’s* red-colored nectar

We next considered a functional role of *Nesocodon’s* red-colored nectar. Several Mauritian plants with colored nectars, including *Nesocodon*, are known or suspected to be pollinated by day geckos (13). For instance, a previous field study reported that two Mauritian plants with colored nectar are visited and pollinated by day geckos (*Phelsuma* spp.) and that artificial nectars containing commercially-available red food coloring are attractive to day geckos (13). However, these suggestive observations lack conclusive data linking the specific nectar pigments to the sensory system and behavior of their putative pollinators. To investigate the function of nesocodin as an attractant to day geckos, we used a physiological model of gecko vision and a behavioral experiment to assess the visibility and attractiveness of nesocodin to gold dust day geckos, *Phelsuma laticauda*. We measured nectar reflectance within the context of the surrounding petals (Fig. 5A) in order to model visibility and conspicuity with respect to known retinal response patterns (Fig. 5B & 5C). *Phelsuma* have four types of photoreceptors (ultraviolet, blue, green, red) and hence their visual space can be represented in the form of a tetrahedron with each vertex corresponding to a specific photoreceptor (18, 19). The reflectance spectra of *Nesocodon* nectar (Fig. 5A) mapped onto this visual space indicated that the red color of the nectar should be visible to *Phelsuma* (Fig. 5B). Moreover, the contrast between the nectar and the surrounding petals is highly conspicuous (Fig. 5C).

**Figure 5.**
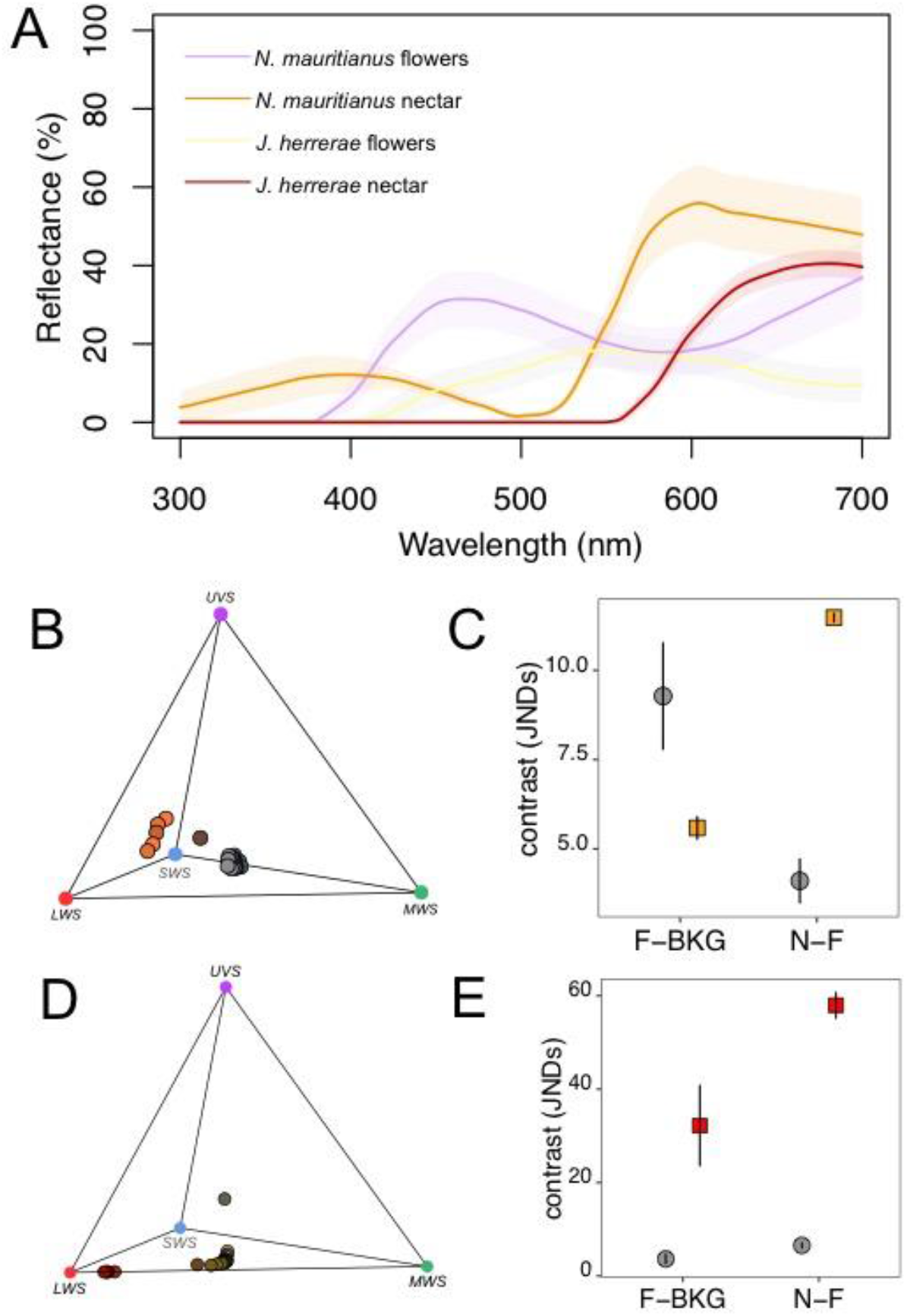
Nectars containing nesocodin are visible and conspicuous to diurnal geckos and birds. **(A)** Reflectance of *N. mauritianus* and *J. herrerae* nectar (note that the curves overlie one another) and the surrounding petals. Shaded areas around the curves represent one standard deviation **(B)** Tetraplot showing *N. mauritianus* reflectance data from panel A projected onto the visual space of diurnal gecko *Phelsuma ornata*. The vertices of the tetrahedron correspond to four different photoreceptors. **(C)** Achromatic contrast (gray points) and chromatic contrast (orange points) between nectar and the adjacent petals (N-F) and petals and the background (F-BKG) for *N. mauritianus*. **(D)** Tetraplot showing *J. herrerae* reflectance data from panel A projected onto the visual space of Green-backed fire crown hummingbird *Sephanoides sephaniodes*. **(E)** Achromatic contrast (gray points) and chromatic contrast (red points) between nectar and the adjacent petals (N-F) and petals and the background (F-BKG) for *J. herrerae*. Contrasts are expressed in units of JND (just noticeable differences), and the higher the value the more conspicuous the color should appear to the pollinators. Error bars are 95% CI.

In a behavioral preference test (Fig. 6A, fig. S27), *Phelsuma* were allowed to visit and consume two artificial nectars that differed solely in the absence or presence of synthetic nesocodin, and hence color (clear versus red), but were otherwise chemically identical. The geckos not only investigated sources of red nectar containing nesocodin more often than sources of clear nectar without nesocodin, but they also consumed significantly more of the red-colored nectar (Fig. 6B & 6C, table S1). These cumulative results strongly suggest that nesocodin’s red color is both visible and attractive to *Phelsuma* day geckos, thus supporting a role in pollinator attraction as a conspicuous ‘honest reward’, as previously suggested (10, 11, 20, 21).

**Figure 6.**
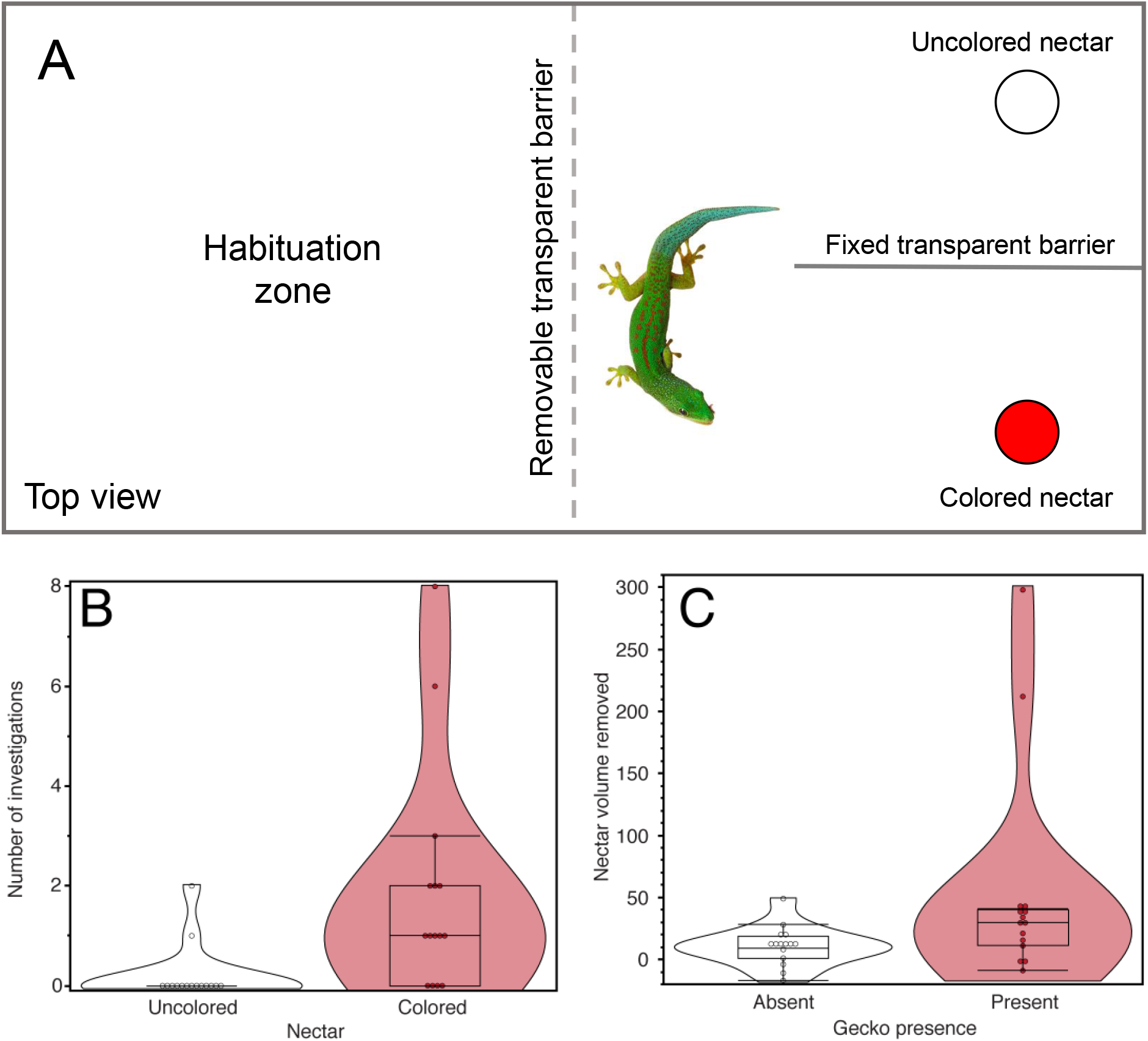
Nectars containing nesocodin are attractive to diurnal geckos (*Phelsuma*). **(A)** Overhead schematic of the experimental setup for gecko behavioral tests in terraria (also see fig. S27). **(B)** Number of investigations made by geckos (N = 15) to tubes containing synthetic nectars with or without 3 mM nesocodin. Sources of nectar with nesocodin were visited significantly more often than sources without nesocodin (Wilcoxon signed ranks test: W = 60, z = 2.65, two-tailed P = 0.008). **(C)** The net volume of nectar removed by geckos (N = 15) relative to evaporative control nectars in adjacent terraria without geckos. Significantly more of the nectar with nesocodin was consumed by the geckos (Wilcoxon signed ranks test: W = 77, z = 2.4, two-tailed P = 0.0164).

### Co-evolution of an identical trait in a hummingbird-pollinated plant

Given the finding of a novel pigment in a Mauritian nectar, we investigated the possibility of distantly related species with red-colored nectars also containing nesocodin. Toward this end, we evaluated the red nectar of *Jaltomata herrerae* and the non-colored nectar of one of its close relatives, *Jaltomata procumbens*. *Jaltomata* spp. with red nectars are visited by hummingbirds in the wild, whereas ones with non-colored nectars tend to be visited by insects (22). Furthermore, *J. herrerae* is endemic to the high elevations (ca. 3000-4000 m) of the mountains of southern Peru and Bolivia, and *J. procumbens* is native to a range from Arizona (U.S.A.) down to northern South America (personal communication, Thomas Mione, Central Connecticut State University, and Segundo Leiva G., Universidad Privada Antenor Orrego). Estimates of divergence times indicate that *Nesocodon* and *Jaltomata* last shared a common ancestor approximately 104 million years ago (23). Like *Nesocodon, J. herrereae* nectar is highly alkaline (pH ~9) and contains nesocodin (Fig. 7A, fig. s29), whereas *J. procumbens* nectar is acidic (pH ~6) and colorless. Similarly, *J. herrerae* nectar contains both an α-carbonic anhydrase (α-CA) and an alcohol oxidase (AO in Fig. 7, fig. S30) analogous to the ones found in *Nescodon* (see below); however, while the acidic nectar of *J. procumbens* also contains an alcohol oxidase, it lacks the carbonic anhydrase and associated activity found in both *J. herrerae* (Fig. 6A & 6B) and *Nesocodon* nectars (Fig. 2). Finally, modeling of *J. herrereae* nectar and petal reflectance (Fig. 5A) onto the visual space of hummingbirds suggests that the nectar should also be highly visible and conspicuous (Fig. 5D & 5E), as predicted for the *Nesocodon-Phelsuma* system (Fig. 5B & 5C).

**Figure 6.**
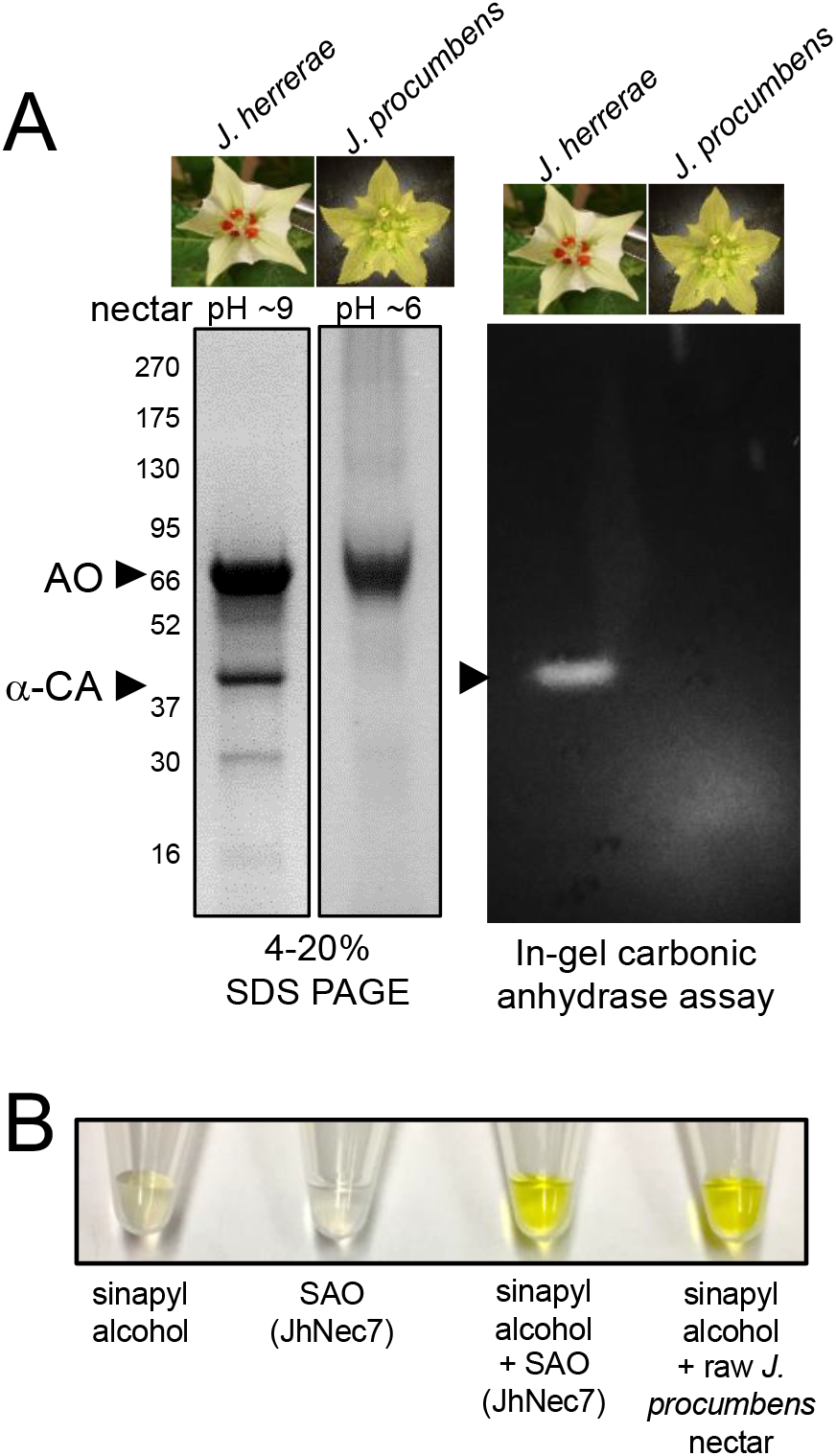
*Jaltomata herrerae* nectar contains nesocodin and analogous enzymes for its production. **(A)** SDS PAGE analysis of nectar proteins (left panels) from *J. herrerae* and *J. procumbens* and in-gel carbonic anhydrase activity assays (right). The arrowheads indicate the locations of an α-carbonic anhydrase (α-CA) and alcohol oxidase (AO). **(B)** Sinapyl alcohol oxidase activity in *Jaltomata* nectars. Images of sinapyl alcohol and purified alcohol oxidase (SAO) from *J. herrerae* are shown in the left two tubes. The right two tubes show the production of yellow sinapaldehyde after mixing sinapyl alcohol with either purified enzyme from *J. herrerae* or raw nectar from *J. procumbens* (note: *J. procumbens* produces too little nectar to allow for purification of the enzyme prior to the assay).

While interesting, the finding of nesocodin in both *Nesocodon* and *Jaltomata* nectars could be a conserved or independently evolved trait. Two key pieces of information strongly suggest that a case of convergent evolution has occurred in the production of this novel nectar pigment in these two members of the long-diverged Campanulaceae (*Nesocodon*) and Solanaceae (*Jaltomata*).Firstly, the carbonic anhydrases found in *Nesocodon* and *J. hererrae* nectars only share ~42% identity (figure S30) and have closer homologs in each other’s genomes. Secondly, the alcohol oxidases found in *Nesocodon* and *Jaltomata* nectars are clearly not orthologous (~21% identity, figure S31), as they belong to two different enzyme families (see alignment in fig S30). Specifically, *Nesocodon* Nec3 (alcohol oxidase) belongs to the GMC flavoenzyme oxidoreductase family, whereas the *Jaltomata* alcohol oxidase is a member of the berberine-bridge family of enzymes within the FAD/FMN-containing dehydrogenase superfamily. These findings strongly suggest that the genes encoding these enzymes have independently evolved to have the same function in pigment formation.

## Conclusion

This work cumulatively reports the identification of a new, convergently-evolved plant pigment and its associated synthesis. These findings add to a growing list of convergent evolution in complex biosynthetic pathways (24–27). More specifically, it illustrates how modulation of the nectar chemical environment, largely through action of a carbonic anhydrase, impacts the production of red-colored nectars and how distantly related plants have independently converged on the same biochemical solution of how to produce a red nectar. These red pigments likely function, at least in part, to attract and direct the vertebrate pollinators these plants rely on in habitats with few potential insect pollinators, like island cliff sides (*Nesocodon* of Mauritius) and mountains (*Jaltomata* in the Andes of South America). Indeed, it has been speculated that geographically isolated areas with numerous vertebrates, but relatively few insects, may have given rise to plant species with colored nectars (21). We anticipate our findings will serve as a starting point for more in-depth comparative biochemical and behavioral studies on pigment-driven plant-pollinator interactions within an evolutionary context.

## Supporting information

Table S1 - Gecko behavioral data

Video S1 - Gecko drinking nectar

## ACKNOWLEDGEMENTS

We thank the University of Minnesota Center for Mass Spectrometry and Proteomics and the Minnesota NMR Center (with support from the National Institute of General Medical Sciences [1S10OD021536 (G.Veglia)]. We also thank Lisa Aston-Philander, Angie Koebler, and Alex Eilts of the CBS Conservatory of the University of Minnesota for helpful discussions and access to plant materials.

## Author contributions

RR, SAB, AK, CH, EMS, ADH and CJC identified sinapaldehyde and nesocodin in the nectar. ADH performed structural characterization of nesocodin by NMR. RR, AK and CJC studied nesocodin biosynthesis in vivo via the nectar enzymes. AM and MH investigated gene expression in *Nesocodon* nectaries. AZ and ESR performed the pollinator visual modeling studies. NM, YC and MAB conducted and analyzed gecko behavior towards nectars containing nesocodin. CC and AH wrote a majority of the manuscript, with all co-authors providing extensive addit ions and editing.

## Data and materials availability

Accession numbers are pending for RNA-seq data and nectar protein accessions. All data is available in the main text or the supplementary materials. Samples of *Nesocodon* and *Jaltomata* nectar are available upon request.

## SUPPLEMENTARY MATERIALS

Materials and Methods

Supplementary text

Fig S1 – S31

Table S1

Video S1

## REFERENCES CITED

1. M. C. P. P. Reis-Mansur et al., Carotenoids from UV-resistant Antarctic *Microbacterium* sp. LEMMJ01. Sci Rep-Uk 9 (2019).

2. J. Chory et al., From seed germination to flowering, light controls plant development via the pigment phytochrome. P Natl Acad Sci USA 93, 12066–12071 (1996).

3. I. C. Cuthill et al., The biology of color. Science 357 (2017).

4. H. M. Schaefer, V. Schaefer, D. J. Levey, How plant-animal interactions signal new insights in communication. Trends Ecol Evol 19, 577–584 (2004).

5. E. Moyroud, B. J. Glover, The evolution of diverse floral morphologies. Current Biology 27, R941–R951 (2017).

6. M. T. Clegg, M. L. Durbin, Flower color variation: A model for the experimental study of evolution. P Natl Acad Sci USA 97, 7016–7023 (2000).

7. R. A. Raguso, R. A. Levin, S. E. Foose, M. W. Holmberg, L. A. McDade, Fragrance chemistry, nocturnal rhythms and pollination “syndromes” in *Nicotiana*. Phytochemistry 63, 265–284 (2003).

8. M. Vanderplanck et al., The importance of pollen chemistry in evolutionary host shifts of bees. Sci Rep-Uk 7 (2017).

9. A. G. Dyer et al., Parallel evolution of angiosperm colour signals: common evolutionary pressures linked to hymenopteran vision. P Roy Soc B-Biol Sci 279, 3606–3615 (2012).

10. D. M. Hansen, J. M. Olesen, T. Mione, S. D. Johnson, C. B. Muller, Coloured nectar: distribution, ecology, and evolution of an enigmatic floral trait. Biological Reviews 82, 83–111 (2007).

11. F. P. Zhang, Z. Larson-Rabin, D. Z. Li, H. Wang, Colored nectar as an honest signal in plant-animal interactions. Plant Signal Behav 7, 811–812 (2012).

12. J. M. Olesen et al., Mauritian red nectar remains a mystery. Nature 393, 529 (1998).

13. D. M. Hansen, K. Beer, C. B. Muller, Mauritian coloured nectar no longer a mystery: a visual signal for lizard pollinators. Biol Letters 2, 165–168 (2006).

14. C. Carter, S. Shafir, L. Yehonatan, R. G. Palmer, R. Thornburg, A novel role for proline in plant floral nectars. Naturwissenschaften 93, 72–79 (2006).

15. Y. R. Mo, S. D. Peyerimhoff, Theoretical analysis of electronic delocalization. J Chem Phys 109, 1687–1697 (1998).

16. P. J. Linser, K. E. Smith, T. J. Seron, M. N. Oviedo, Carbonic anhydrases and anion transport in mosquito midgut pH regulation. J Exp Biol 212, 1662–1671 (2009).

17. J. W. Whittaker, Non-heme manganese catalase--the ‘other’ catalase. Arch Biochem Biophys 525, 111–120 (2012).

18. G. B. Arden, K. Tansley, The electroretinogram of a diurnal gecko. J Gen Physiol 45, 1145–1161 (1962).

19. Y. Taniguchi, O. Hisatomi, M. Yoshida, F. Tokunaga, Pinopsin expressed in the retinal photoreceptors of a diurnal gecko. FEBS Lett 496, 69–74 (2001).

20. K. Ito, M. F. Suzuki, K. Mochizuki, Evolution of honest reward signal in flowers. P Roy Soc B-Biol Sci 288 (2021).

21. I. A. Minnaar, A. Kohler, C. Purchase, S. W. Nicolson, Coloured and toxic nectar: Feeding choices of the Madagascar Giant Day Gecko, *Phelsuma grandis*. Ethology 119, 417–426 (2013).

22. K. C. Plourd, T. Mione, Pollination does not affect floral nectar production, and is required for fruit-set by a hummingbird-visited Andean plant species. Phytologia 98, 313–317 (2016).

23. S. Kumar, G. Stecher, M. Suleski, S. B. Hedges, TimeTree: A resource for timelines, timetrees, and divergence times. Molecular Biology and Evolution 34, 1812–1819 (2017).

24. R. Q. Huang, A. J. O’Donnell, J. J. Barboline, T. J. Barkman, Convergent evolution of caffeine in plants by co-option of exapted ancestral enzymes. P Natl Acad Sci USA 113, 10613–10618 (2016).

25. K. Heyduk, J. J. Moreno-Villena, I. S. Gilman, P. A. Christin, E. J. Edwards, The genetics of convergent evolution: insights from plant photosynthesis. Nat Rev Genet 20, 485–493 (2019).

26. R. Greenway et al., Convergent evolution of conserved mitochondrial pathways underlies repeated adaptation to extreme environments. P Natl Acad Sci USA 117, 16424–16430 (2020).

27. N. B. Jensen et al., Convergent evolution in biosynthesis of cyanogenic defence compounds in plants and insects. Nat Commun 2 (2011).

