## Supplementary material for "Convergent evolution of a novel blood-red nectar pigment in vertebrate-pollinated flowers": Materials & Methods and Supp Figures

**This PDF file includes:**

Materials and Methods
Supplemental text
Figs. S1 to S31
Supplemental References 1-21

### Materials and Methods

Plant materials
*Nesocodon mauritianus* was curated and maintained by the College of Biological Sciences Conservatory at the University of Minnesota, St. Paul, Minnesota, U.S.A. Plants were grown in the Cloud Forest + Maritime Climate Room with the temperature maintained between 18 and 24 ˚C and 90-100% relative humidity.

pH measurements
Nectar pH was determined by mixing raw nectar 1:1 with 100 μg/mL bromothymol blue dissolved in di$H_2O$ and measuring the absorbance at 613 nm relative to a standard curve consisting of 50 mM buffers at pH 6.5 (MES), 7.0 (HEPES), 7.5 (HEPES), 8.0 (HEPES), 8.5 (Tricine) and TAPS (9.0). The relative accuracy of pH calculations were either confirmed with colorpHast® 5-10 pH strips (EM-Reagents Cat. 9588) or by direct measurement with a Spectrum Technologies Scout 400 pH meter with a blunt tip ISFET probe by placing the nectar in the rubber storage cap.

UV/visible absorbance spectroscopy
A BioTek PowerWave HT 96-well plate reader or an Implen NanoPhotometer™ Pearl were used for all spectrophotometric analyses.

Nectar proteomics
Eighteen microliters of *Nesocodon mauritianus* nectar was electrophoresed on a 4-20% Tricine gel under denaturing conditions and stained with PAGE-Blue (Thermo-Scientific Cat#24620). Protein bands were excised and submitted for identification at the University of Minnesota's Center for Mass Spectrometry and Proteomics Facility. In-gel trypsin digestion and STAGE Tip peptide cleanup was performed as previously described (1) except that iodoacetamide was used as the alkylating reagent during the digest protocol. The peptides were analyzed by capillary LC-MS/MS on an Orbitrap Velos system with data dependent acquisition (Orbi-Orbi mode with HCD activation) as described previously (2) . The PEAKS Studio 8.5 (3) (Bioinformatics Solutions, Waterloo, ON, CA) was used for database searching. We formatted a custom protein sequence database from the theoretically translated Nesocodon nectary RNASeq data merged with NCBI Reference Sequence *Arabidopsis thaliana* (tax id 3702) downloaded on 5/14/2018 and common lab contaminant proteins (from https://www.thegpm.org/crap/). Search parameters were: de novo precursor tolerance 20 ppm and fragment ion tolerance 0.1 Da; database search precursor mass tolerance 50.0 ppm; fragment mass error tolerance 0.1 Da; precursor mass search type monoisotopic; trypsin enzyme specificity; fixed modification cysteine carbamidomethylation; variable modifications methionine oxidation, pyroglutamic acid, deamidation of asparagine and glutamine, protein N-terminal acetylation; maximum variable modifications per peptide 3; false discovery rate calculation On; spectra merge off; charge state correction; spectral filter quality >0.65. We employed the SPIDER module for consideration of amino acid substitutions in the target protein sequence database.

NmNec3 purification
For a standard preparation, 250 μL nectar was diluted 1:1 with 50 mM HEPES, pH 8.0, heated for two minutes at 50˚C and then centrifuged through a 50000 MWCO Amicon® Ultra-

0.5 Centrifugal Filter (MilliporeSigma) for 5 minutes at 14,000 xg, with the remaining ~50 μL of retentate diluted back up to 500 μL in 50 mM HEPES, pH 8.0 and centrifugation repeated four more times. Purity of the final product was evaluated by 10% SDS PAGE under denaturing conditions and gel staining with PAGE-Blue (Thermo-Scientific Cat#24620). Final protein concentration was determined by UV absorbance at 280 nm and a predicted extinction coefficient of 56,520 $M^{-1}$ $cm^{-1}$ (based on the sequence of the mature protein without the predicted signal peptide).

<u>Preparation of de-proteinated (ultrafiltered) nectar and total nectar protein</u>

Total protein was removed from 400 μL of raw nectar via centrifugation through a 3000 MWCO Amicon® Ultra-0.5 Centrifugal Filter (MilliporeSigma) for 10 minutes at 14,000 xg. The flow-through was collected as 'de-proteinated' nectar and validated for protein removal by 4-20% SDS PAGE. Meanwhile, the retentate was brought back up to 400 μL with 50 mM HEPES, pH 8.0 and re-centrifuged and washed a total of five times to remove nectar sugars and other small molecule components.

<u>Enzymatic assays</u>

<u>In-gel carbonic anhydrase assay</u>

Fresh *Nesocodon* nectar was collected and 18 μL was immediately subjected to standard SDS PAGE (4-20%), except the loading buffer contained no β-mercaptoethanol and the samples were not boiled prior to loading the gel. Following electrophoresis, the gel was sliced vertically with one lane being stained in PAGE Blue and the remaining lanes being processed for an in-gel carbonic anhydrase activity assay as previously described (4), except that dry ice was used to bubble $CO_2$ into the distilled water instead of a compressed $CO_2$ tank. The positive control carbonic anhydrase was purchased from Millipore Sigma (C9207-1VL).

<u>Colorimetric carbonic anhydrase assay</u>

Assays with raw 0 h nectar contained either carbonic anhydrase inhibitors (acetazolamide, sulfanilamide or 6-ethoxy-2-benzothiazolesulfonamide) or an equivalent amount of DMSO (10% DMSO (v/v) for acetazolamide and sulfanilamide, 50% DMSO (v/v) for 6-ethoxy-2-benzothiazolesulfonamide). Reactions were initiated by adding $NaHCO_3$, pH 7.45 at a final concentration of 4 mM, with the absorbance at 420 nm (positively correlating to an increase in pH) being monitored for three minutes. Assays recapitulating that of raw nectar contained 125 μM sinapaldehyde in 5 mM $NaHCO_3$ (5 mM final) either with or without commercial carbonic anhydrase (Sigma C9207-1VL) added.

<u>Sinapyl alcohol oxidase assay</u>

Reactions consisted of 1 mM sinapyl alcohol and 0.1 mg/mL NmNec3 in 50 mM $Na(H)CO_3$, pH 9.0 incubated at 21˚C for 30 minutes followed by conducting wavelength scans of 300-600 nm. Negative controls with sinapyl alcohol but no NmNec3 did not yield any sinapaldehyde (grey line); a 0.5 mM sinapaldehyde standard in 50 mM $Na(H)CO_3$, pH 9.0 was used as a reference.

<u>In-gel alcohol oxidase assay</u>

Twelve μL of raw nectar was directly loaded (no loading buffer) into separate lanes of a 4-20% TRICINE PAGE gel and electrophoresed under non-denaturing conditions at 120 V for ~1 h. The gel was then incubated at 37°C in a staining solution containing 25 mM TRIS, pH 8.2 and either 10% methanol (MeOH), 10% ethanol (EtOH), 10% 1-propanol, 10% 2-propanol or 5 mM sinapyl alcohol, along with 5 U/mL horseradish peroxidase (HRP), 0.5mg/mL 3,3' diaminobenzidine, 20% EtOH and 0.1 mg/mL flavin adenine dinucleotide for ~3 hours.

In-gel catalase assay

18 μL of fresh, raw nectar was electrophoresed on a 4-20% Tricine gel in the presence of SDS, but without boiling or β-mercaptoethanol. Catalase activity was detected in the gel as previously described (5).

Spectrophotometric catalase assay

Assays were performed by monitoring the breakdown of $H_2O_2$ by measuring the absorbance at 250 nm (6) . Briefly, individual assays contained 25 mM $H_2O_2$ in 50 mM HEPES pH 8.0 and either 0.17 mg/mL of total nectar proteins (either boiled or nonnboiled) or an equivalent volume of HEPES buffer as a negative control and the absorbance was monitored at 250 nm over time.

In-situ pharmacological treatments

Individual nectar droplets from newly opened flowers (0 h) were treated by the addition of carbonic anhydrase inhibitors (acetazolamide, sulfanilamide or 6-ethoxy-2-benzothiazolesulfonamide) or equivalent amounts of DMSO [10% DMSO (v/v) for acetazolamide and sulfanilamide, 50% DMSO (v/v) for 6-ethoxy-2-benzothiazolesulfonamide] and thorough mixing by repeated pipetting. After 8-20 hours, nectar was removed and analyzed for pH and subjected to absorbance wavelength scans.

Fluorescence spectroscopy

Raw nectar and synthetic nesocodin (sinapaldehyde + proline) were diluted 1:99 in 25 mM HEPES, pH 8.0 and the excitation and emission spectra were measured with a BioTek Synergy™ MX Microplate Reader.

Thin-layer chromatography of synthetic nesocodin

Two microliters each of synthetic nesocodin (1 mM sinapaldehyde + 10 mM proline in 50 mM HEPES pH 8.0), raw nectar or 1 mM sinapaldehyde alone were spotted onto silica gel-on-glass TLC plates (Sigma Z12,268-8) with a mobile phase of 100 mM sodium acetate, pH 4.8 being used under ambient air. Plates were air dried and imaged under UV light (340 nm).

LC-HRMS analysis

Chromatographic separation and high-resolution mass spectral analyses of *Nesocodon* nectar and synthetic nesocodin were performed using a UHPLC coupled to a hybrid quadrupole-Orbitrap mass spectrometer (Ultimate® 3000 HPLC, Q Exactive™, Thermo Scientific). The UHPLC is equipped with a flow-through Photo Diode Array (PDA) detector allowing concurrent UV/visible spectrophotometric and MS analysis of separated analytes post-column. Nectar samples were first purified by $C_{18}$ solid phase extraction using a Ziptip (Millipore) conditioned in ~20 µL of acetonitrile, washed with 20 µL of 0.1% formic acid in water, loaded with ~20 µL of nectar and washed with ~100 µL of 0.1% formic acid in water prior to elution with ~10 µL of acetonitrile. The eluate was then transferred into LC-MS autosampler vials and 1 µL was injected (via autosampler) onto a reversed-phase $C_{18}$ HSS T3 1.8 µm particle size, 2.1 x 100 mm column (Waters). Column temperature was 40°C and solvent flow rate 0.45 mL/min. A 20-minute linear gradient using mobile phases A: 0.1% formic acid in water and B: 0.1% formic acid in acetonitrile was run according to the following gradient 25 minute elution profile: initial, 2% B; 20 min, 98% B; 21 min, 98% B; 22 min, 2% B (hold to 25 min). The following MS conditions were used: full scan mass scan range: 100-1000 *m/z,* resolution: 70,000, data type: profile, desolvation temperature 350°C, capillary voltage: 3800 V (+), 4000 V (-). Xcalibur™ software version 2.1 (Thermo Scientific) was used to record and visualize the chromatograms and spectra. Tandem MS (MS/MS) spectra of nectar-derived sinapaldehyde and nesocodin were sequentially fragmented in the HCD collision cell with normalized collision energy of 10%, 20%, 25%, 30%, 35% and 40%. MS/MS scans were acquired with 17.5 k resolution and the target value was $2.0 \times 10^5$ with 100 ms of maximum injection time. An isolation width of 2.0 *m/z* was used for precursor ion selection in MS/MS mode.

Proline quantification

Proline in *Nesocodon* nectar was quantified using isotope dilution and a GC-MS-based amino acid analysis method (7). Briefly, three 100 µL aliquots of red (+48 h) *Nesocodon* nectar were prepared and 99 atom% $^{13}C$ L-[$^{13}C_5$]proline (604801, Sigma) was added to a concentration of 10 mM. The samples were acidified by adding 1 mL of 0.01 M HCl and amino acids were loaded by addition to a strong cation exchange SPE column (AT209800, Alltech) preconditioned with 1 mL of 0.01 M HCl and 3 additions of 1 mL of di$H_2O$. Once loaded, the SPE columns were washed twice with 1 mL of 80% (v/v) methanol in water before aminoacids were eluted with 250 µL of a 1 to 1 (v/v) mixture of 8 M $NH_4OH$ and methanol. Aliquots (50 µL) of each eluate in dry glass vials were derivatized by adding 5 µL of pyridine (Sigma) and 5 µL of methyl chloroformate (Sigma). The reactions were allowed to proceed for 1 min. before 90 µL of chloroform and 90µL of 50 mM sodium bicarbonate in water were added to stop the reaction. The chloroform layers were removed and dried over anhydrous sodium sulfate prior to GC-MS analysis. GC-MS was performed on an Agilent 7890A/5975 MSD instrument using 2 µL sample injection volume in splitless mode, a DB-50 capillary GC column, inlet temperature of 240 ˚C and interface temperature of 290 ˚C. The initial oven temperature was 70 ˚C and was held for 3 min. following injection and was increased at a rate of 25 ˚C/min. to 280 ˚C where it was held for 5 min. before returning to initial settings. Data were analyzed using *ChemStation* software.

In parallel experiments, proline in 0 h and +24 h nectar was evaluated by thin layer chromatography. In brief, 2 µl total (in 1 µl increments) of 0 h and +24 h were spotted onto 10 x 10 cm HPTLC cellulose-on-glass plates (Merck 1.05787.0001). After airing drying the spotted plates, separation was performed in an air-tight glass container with a mobile phase of 60:25:15 of 1-butanol:$H_2O$:acetic acid. Plates were then air dried and developed by spraying with 0.3% ninhydrin in 1-butanol/3% acetic acid. Proline reaction with ninhydrin uniquely develops as a yellow product, whereas most other amino acids generate a purple or red products.

Nesocodin derivatization by reduction with $NaBH_4$ or $NaBD_4$.

Nesocodin was synthesized in a 1 mL reaction containing 10 mM proline and 1 mM sinapaldehyde in 50 mM Na(H)$CO_3$ buffer pH 9.0. The reaction started to turn visibly red immediately and was allowed to incubate at 21˚C for 30 min prior to being divided into 3 equal aliquots (330 µL each). Additions of 23 mg of $NaBH_4$ and 22 mg of $NaBD_4$ were made to the first and second aliquots respectively. Reductant was in excess of sinapaldehyde concentration by over three orders of magnitude with ~1.8 M $NaBH_4$ and ~1.6 M $NaBD_4$. Both reduced aliquots bubbled immediately upon addition of the solid reducing agent but stopped after a few seconds. Reaction tubes were capped and mixed by inversion and the uncapped to release residual pressure. Within the mixing time both reduced aliquots became colorless. The third, red aliquot was reserved as a negative control. All three aliquots were acidified by addition of 330 µL of 1% (v/v) formic acid in water. The first two aliquots bubbled upon addition of the acid and were allowed to react for several minutes uncapped before proceeding. All three aliquots were first purified by $C_{18}$ solid phase extraction using an OMIX 100 µL tip (Agillent) conditioned in ~200 µL of 90% (v/v) methanol in water with 0.1% (v/v) formic acid, washed with 200 µL of 0.085% formic acid in water, loaded with ~600 µL of each aliquot, and washed 3 times with 100 µL of 0.085% formic acid in water prior to elution with ~20 µL of 90% (v/v) methanol in water with 0.1% (v/v) formic acid. Samples were subsequently analyzed by LC-HRMS as described above.

Nesocodin synthesis for structural validation

For NMR and other structural analyses, nesocodin was prepared by dissolving 223.9 mg (1.086 mmoles) of sinapaldehyde in 10 mL methanol (to 0.1086 M) and then adding 125.0 mg of L- proline (1.086 mmoles) followed by a sub-stoichiometric addition (100 µL, ~0.42 mmoles) of tributylamine (as a sterically hindered base catalyst with low nucleophilicity). As the mixture is allowed to react, a yellow to red color change is observable immediately and the reaction runs to completion within minutes at room temperature. Once the reaction is complete, the contents of the reaction flask are diluted ten-fold into room temperature ethyl acetate. A red solid precipitate consisting of a 64% E- to 36% Z- mixture of nesocodin isomers was collected by filtration, washed with a small volume of ice-cold ethyl acetate and dried *in vacuo* overnight.

Synthetic nesocodin NMR

One milligram of the dried red solid nesocodin was dissolved in one mL of methanol-$d_4$ containing 0.1% TMS as an internal standard and transferred into a 5 mm thin walled quartz NMR tube (Wilmad 535-PP-7). NMR data including 1D $^1$H-NMR, 2D $^1$H-DQF-COSY and 1D $^{13}$C-NMR were obtained on a Bruker 600 MHz NMR at the University of Minnesota Nuclear Magnetic Resonance Center on a fee for service basis and analyzed using TopSpin v4.0.6.

Semi-quantitative RT-PCR analysis

Total RNA from frozen samples was extracted using TRIzol RNA Reagent (Sigma-Aldrich). The RNA was treated with TURBO DNA-free kit (Invitrogen) according to the manufacturer's protocol. First-strand cDNA was synthesized from 1 µg of total RNA using iScript cDNA synthesis kit (Bio-Rad). The semi-quantitative reverse transcription PCR (RT-PCR) was performed with GoTaq DNA polymerase (Promega). The *NmGAPDH* gene was used as an internal control. For PCR reaction, the primers used for RT-PCR are listed in Fig. S17.

Nectary transcriptomic analysis

*Nesocodon* nectary tissue was manually dissected and total RNA was extracted using TRIzol RNA Reagent (Sigma-Aldrich). The RNA was treated with the TURBO DNA-free kit (Invitrogen) according to the manufacturer's protocol and submitted to the University of Minnesota Genomics Center for mRNA isolation, barcoded library creation and Illumina HiSeq

2500 sequencing via paired-end 125 bp runs using Rapid chemistry. A total of 7.3 million paired-end 125 bp reads were generated. Trinity version 2.4.0 (8) was used to assemble 125,688 contigs from the reads *de novo*.

Nectar protein maximum likelihood tree generation

Protein sequences were aligned with MUSCLE 3.8.31 (9) (*20*) and refined with GBLOCKS 0.91b (10). Phylogenetic trees were computed with PhyML 3.1 using the WAG matrix with four gamma-distributed rate categories (11), and edited using the ETE toolkit (12).

Behavioral testing

All capture, handling, and experimental protocols were approved by Institutional Animal Care and Use Committee at George Mason University (IACUC protocol #1478025) and were carried out in accordance with all relevant guidelines and regulations.

Fifteen individuals (8 adult females, 6 adult males, 1 juvenile) of the Malagasy gold dust day gecko (*Phelsuma laticauda*) were used as subjects in a behavioral experiment based on a two-alternative choice test to investigate nectar color preference (Table S1). As in other species belonging to the same genus, gold dust day geckos eat pollen and nectar from flowers in their natural environment in Madagascar (Sauroy-Toucouère pers. comm. and (13). All tested animals were collected from wild populations, and they were housed in the same room at George Mason University in the laboratory of one of the authors (YC). All geckos were housed separately in glass terraria (Exoterra, 30 cm x 30 cm x 45 cm, W × D × H), exposed to 12:12 hour light:dark cycles, and given access to additional light from UV lamps (ReptiSun 10 UVB bulbs). Opaque barriers were placed between terraria to prevent the geckos from seeing individuals in adjacent terraria. The room temperature was maintained between 24.5-26.5°C. Each terrarium contained one or more fake plants and a heat pad for thermoregulation. The humidity level and temperature in the room, as well as the health of each gecko, were monitored daily. Over the course of this study, no gecko showed signs of stress or health problems (e.g., pale coloration or emaciation), and all geckos continued to exhibit normal, species-typical behaviors in their home terraria. No geckos were tested while they were shedding. Water was accessible at all times in a shallow bowl in each terrarium. Geckos were fed three times each week with a combination of crickets dusted with calcium and vitamins, mealworms, and a fruit supplement. Feeding for an experimental subject was temporarily suspended for 3 or 4 days prior to the day it was tested, and the normal feeding regime was resumed immediately after testing of that subject was completed.

Experiments were carried out in a dedicated testing room during the 12-hour light portion of the light:dark cycle and always started at the same approximate time (1100 h). Temperature in the experimental room was recorded at the beginning and end of each experiment and was stable at approximately 25°C over the entire length of the study. Animals were always handled by the same person (author NM), and no one else had access to the tested animals and the testing room over the duration of the study. Animals were tested individually and in a single choice test, with only one animal tested on a single day. No food or water – beside the nectars being tested – were provided during a choice test, which lasted 3 h 45 min.

The experimental set-up consisted of two identical, custom-made arenas (61.5 cm × 30.5 cm × 21 cm, L × W × H) placed next to each other on the floor of the testing room. Each arena was constructed from Plexiglas and consisted of a floor, four walls, a removable lid, a removable transparent barrier (30.5 cm × 21 cm, L × H) bisecting the arena along its long axis into a habituation zone and a choice zone, and a fixed transparent barrier (18 cm × 21 cm, L × H) dividing one end of the arena into two stimulus chambers (Fig. S27). The floor and outside walls of each arena were covered with white paper to avoid distractions during testing. On opposite walls of the two stimulus chambers of each arena, we affixed 1.5-mL Eppendorf tubes that were used for stimulus delivery (Fig. S27). The stimuli consisted of synthetic red colored versus non-colored nectars. Synthetic nectars, with and without nesocodin (colored and non-colored, respectively), contained: 20% sucrose (w/v), 10 mM proline, 1 mM Tricine pH. 8.5, and with or without 3 mM nesocodin (pigment), with the final pH being adjusted to 9.0 in order to match the absorbance spectrum of a freshly collected nectar sample (Fig. S28). Nectar was preserved frozen in aliquots until the day before it was used in a choice test, when it was placed in a refrigerator. One of the two arenas was designated as the “test arena” in which behavioral choice tests were conducted. Two video-cameras were placed at the two ends of the test arena to record a gecko’s behavior over the entire duration of a choice trial from different camera views (Fig. S27; Video S1). The other arena was designated as a “control arena;” its sole purpose was to control for the possibility that colored and non-colored nectars evaporated at different rates during the choice test (i.e., over 3 h 45 min). Over the duration of a choice test, the overhead lights in the room were the only source of light (luminosity in the testing room was measured at 282 lux).

Prior to beginning a choice test, four new Eppendorf tubes were mounted in the test and control arenas, and new aliquots of nectar were removed from the refrigerator and used to fill each Eppendorf tube using a pipettor. Thus, both the test and control arenas contained one Eppendorf tube filled to the top (1.6 ml) with colored nectar and one Eppendorf tube filled to the top (1.6 ml) with non-colored nectar (Fig. S27). The side of the arena on which the colored nectar was placed was randomly determined for each tested gecko and was the same in both arenas. To begin a choice test, the subject was released into the habituation zone of the test arena from a fixed release point (Fig. S27) to begin a 45-min habituation phase during which the gecko was free to explore and acclimate to the new environment of the test arena. The removable transparent barrier bisecting the arena was in place during the habituation phase, allowing the gecko to see the stimuli without being able to access them. At the end of the habituation phase, the removable transparent barrier was removed to commence a 3-h choice phase, during which the gecko was able to move around the entire test arena. No one was in the testing room during the habituation and choice phases except to remove the transparent barrier at the end of the habituation phase. At the conclusion of the choice test, we used a pipettor to measure the volume of nectar that remained in each Eppendorf tube in both the test and control arenas. Following these measurements, the Eppendorf tubes were discarded, and the test arena was washed with soap and hot water to remove any scent or biological marking left by the tested gecko.

We scored three measures of nectar preference from video recordings of behavior and measurements of nectar volume. First, we scored which nectar the subject approached first by noting which stimulus chamber the animal entered first while exploring the choice zone, where entry was defined as occurring when the entire body of the gecko had crossed over an imaginary boundary into a stimulus chamber (Fig. S27). We used a two-tailed binomial test to test the null hypothesis that 50% of the geckos first entered each stimulus chamber. The data were consistent with this null hypothesis: 9 of 15 geckos (60%, two-tailed binomial $P = 0.6072$) visited the side of the chamber with the nectar containing nesocodin first. Second, we counted the number of times a subject investigated each nectar stimulus over the 3-h choice phase, where an investigation was operationally defined as occurring when a gecko touched an Eppendorf tube with its snout or licked the tube or its contents (Video S1). Multiple touches or licks during a single visit to a tube were considered a single investigation for analysis purposes; different investigations were recorded only if the gecko had moved away from the tube and later returned, and they were always minutes apart. We used a two-tailed Wilcoxon signed-ranks test to test the null hypothesis that colored and non-colored nectars were investigated an equal number of times. Finally, we compared the differences between the final volumes of colored and non-colored nectar remaining in the test arena at the end of each choice test to the same differences measured in the control arena after the test. If differential rates of evaporation were the sole cause of any volume differences between the colored and non-colored nectars within an arena, then we expected to find no differences between the two arenas in this response variable. Any difference between the test and control arenas would, thus, represent a measure of the differential volume of colored versus non-colored nectar consumed by the subject during the test. Differences were computed such that positive values indicate greater removal of colored nectar (i.e., non-colored minus colored). We used a two-tailed Wilcoxon signed-ranks test to test the null hypothesis that volume differences in the test and control arenas did not differ.

Visual modeling of day geckos and hummingbirds

To examine what role petal and nectar colors play in pollinator attraction, we generated visual models of day gecko and hummingbird vision. Visual models allowed us to answer two crucial questions about pollinator attraction- 1. Can pollinators perceive petal and nectar colors and 2. How conspicuous do these colors appear to pollinator given their specific visual capacities (14). To construct visual models, we used published cone sensitivities of Ornate day gecko *Phelsuma ornata* (UVS $\lambda_{max}$ = 380, SWS $\lambda_{max}$= 437, MWS $\lambda_{max}$= 470, LWS $\lambda_{max}$= 560; ratio of UVS: SWS: MWS: LWS = 1:1:3.5:6) and Green-backed fire crown hummingbird *Sephanoides sephaniodes* (UVS $\lambda_{max}$ = 371, SWS $\lambda_{max}$= 444, MWS $\lambda_{max}$= 504, LWS $\lambda_{max}$= 560; ratio of UVS: SWS: MWS: LWS = 1:2:2:4) (15-17). For both *P. ornata* and *S. sephaniodes,* we were unable to find data on cone abundance. Hence we used cone abundance data from diurnal agamid (*Ctenophorus ornatus*), and chicken (*Gallus gallus domesticus*) as substitutes for *P.ornata* and *S. sephaniodes* respectively (16, 18). For visual modelling, we used RNL models (Receptor-Noise Limited models), set the irradiance conditions to bright daylight (‘D65’) and applied von Kries transformation to account for light adaptation (19, 20). Subsequently, we measured reflectance of *N. mauritianus* and *J. herrerae* petals and nectar (N = 5 flowers each) using a UV-VIS spectrophotometer (OceanOptic JAZ-A2474 with PX lamp). Reflectance was measured by setting the triggering rate of the PX lamp to 10ms, and the boxcar to 5. The spectral probe was maintained at an angle of 45° against the petals, which were placed on black velvet paper. This records reflectance of approximately 3mm x 3mm region of the petals. For nectar, 200ul freshly collected nectar was transferred onto a tissue paper and allowed to dry. Subsequently, reflectance data of the dried nectar was recorded using the same protocol used for measuring reflectance of petals. All measurements were corrected against white and black reflectance standards. These spectral data were then imported in R, smoothened ($\alpha$ = 0.20 or 0.35) and negative values were fixed to ‘0’ using the ‘fixneg’ function in ‘pavo2.0’ R package (14). These spectra were then projected onto the pollinator specific visual space generated using photoreceptor sensitivities (see above). Day geckos and hummingbirds both have four distinct photoreceptor types, and hence their visual spaces can be represented as tetrahedrons whose vertices correspond to each of the four photoreceptors (fig. 5 B, D). The location of petal and nectar spectra (represented as individual points) in this visual space depend on the estimated stimulation of each cone type in response to these spectra. Spectra that fall within the tetrahedral space lie within the perceptual range of the pollinator and their corresponding colors can be perceived by pollinators. We found that spectra of petals and nectar of both *N. mauritianus* and *J. herrerae* lie within visual spaces of day gecko and hummingbird respectively (fig. 5 B, D). This indicates that day geckos and hummingbirds can indeed perceive the red nectar and colored petals of *N. mauritianus* and *J. herrerae*.

Additionally, to examine how conspicuous petal and nectar colors appear to the pollinators, we used the same visual models to calculate chromatic contrast (differences in hue and chroma) and achromatic contrast (difference in luminance) of petals and nectars against natural backgrounds as likely viewed by the pollinators. Higher contrast values indicate greater conspicuousness which is typically associated with greater pollinator attraction. *Nesocodon mauritianus* grows on open brown rocky cliffs so we used spectra of brown rocks as background to calculate contrast for petals. *J. herrerae* on the other hand have dense leaf growth around their flowers and hence we used spectra from conspecific leaves as background to calculate contrast for their petals. To calculate contrasts *N. mauritianus* and *J. herrerae* nectars, their respective petals were used as backgrounds. Here it should be noted that in birds and lizards both, only the *LWS* cone is thought to mediate the perception of luminance, hence sensitivity of only the *LWS* cone was used to calculate achromatic contrasts (18, 21).

We found that for both *N. mauritianus* and *J. herrerae*, chromatic contrast of the nectar was significantly higher than the achromatic contrast of the nectar and the chromatic and achromatic contrasts of the petals (Pairwise contrasts: *N. mauritianus:* all $t > 2.6$, $p < 0.04$; *J. herrerae:* all $t > 9$, $p < 0.03$; Fig. 5 C, E). This suggests that red colored nectar is the most conspicuous signal produced by both *N. mauritianus* and *J. herrerae* from their pollinators perspective and may therefore play an important role in pollinator attraction.

**Supplemental text**

**On the molecular model of nesocodin synthesis shown in Fig. 4E.** In our model, sinapyl alcohol and proline are respectively secreted from nectary parenchymal cells into the apoplast/nectar by a putative, and yet unknown, ABC transporter (ABC?) and amino acid permease (AAP?), respectively. Concurrently, the nectar may be supplied with bicarbonate (carbonic acid) through nectary respiration and export through a bicarbonate (BOR?) transporter.

Fig. S1

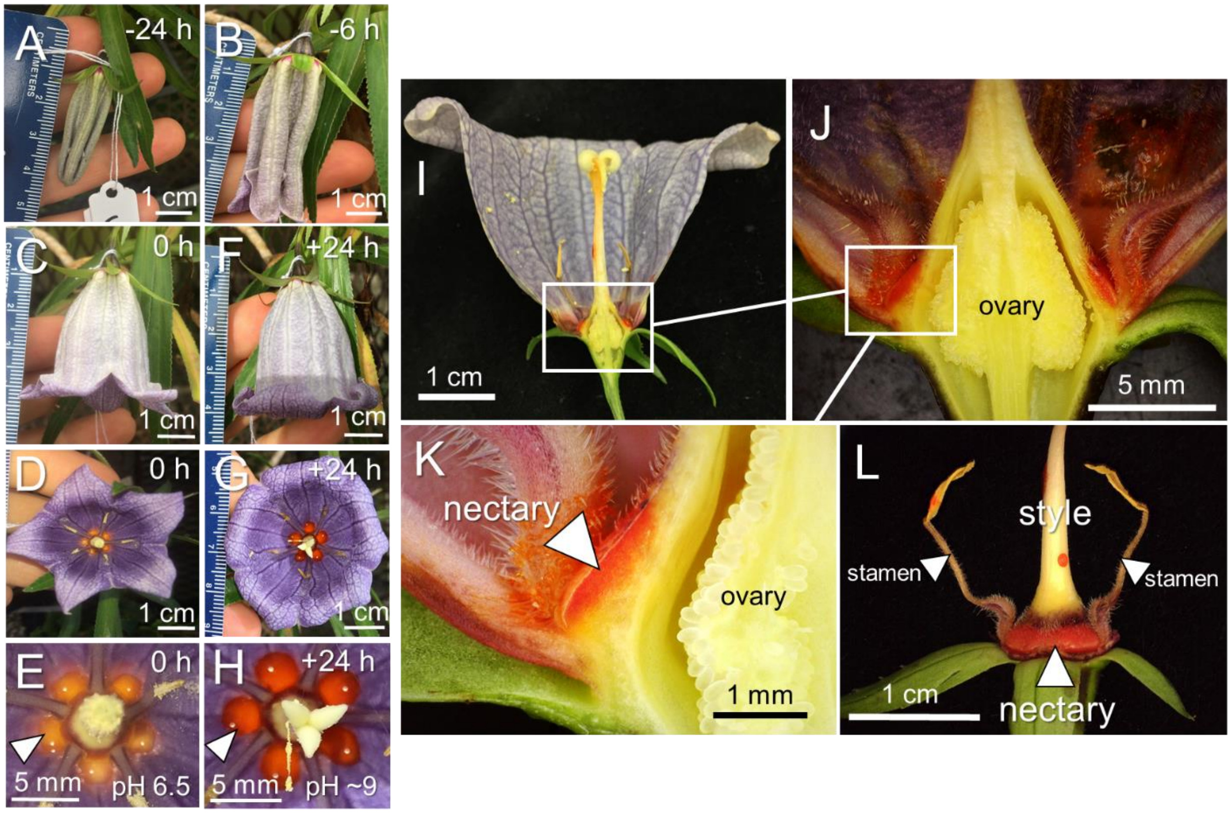


**Fig. S1. Example of developmental progression of *N. mauritianus* flowers used in this study**

**(A and B)** Small flower bud ~24 and ~6 h prior to fully opening.

**(C through E)** New fully open flower (0 hr) as viewed from the side (C) and top (D and E); panel E is a close-up of pane D showing color of five distinct nectar droplets (arrowhead points to one yellow droplet).

**(F through H)** Open flower ~24 hours after opening as viewed from the side (F) and top (G and H); panel H is a close-up of pane G showing color of five distinct nectar droplets (arrowhead points to one droplet). The pH of the nectar droplets is noted in panels E (0 h) and H (+24 h).

**(I through K)** Longitudinal section of fully mature flower displaying successive close-ups of the red nectary.

**(L)** Flower with petals removed to display the red nectary located at the base of the style and stamens.

Fig. S2

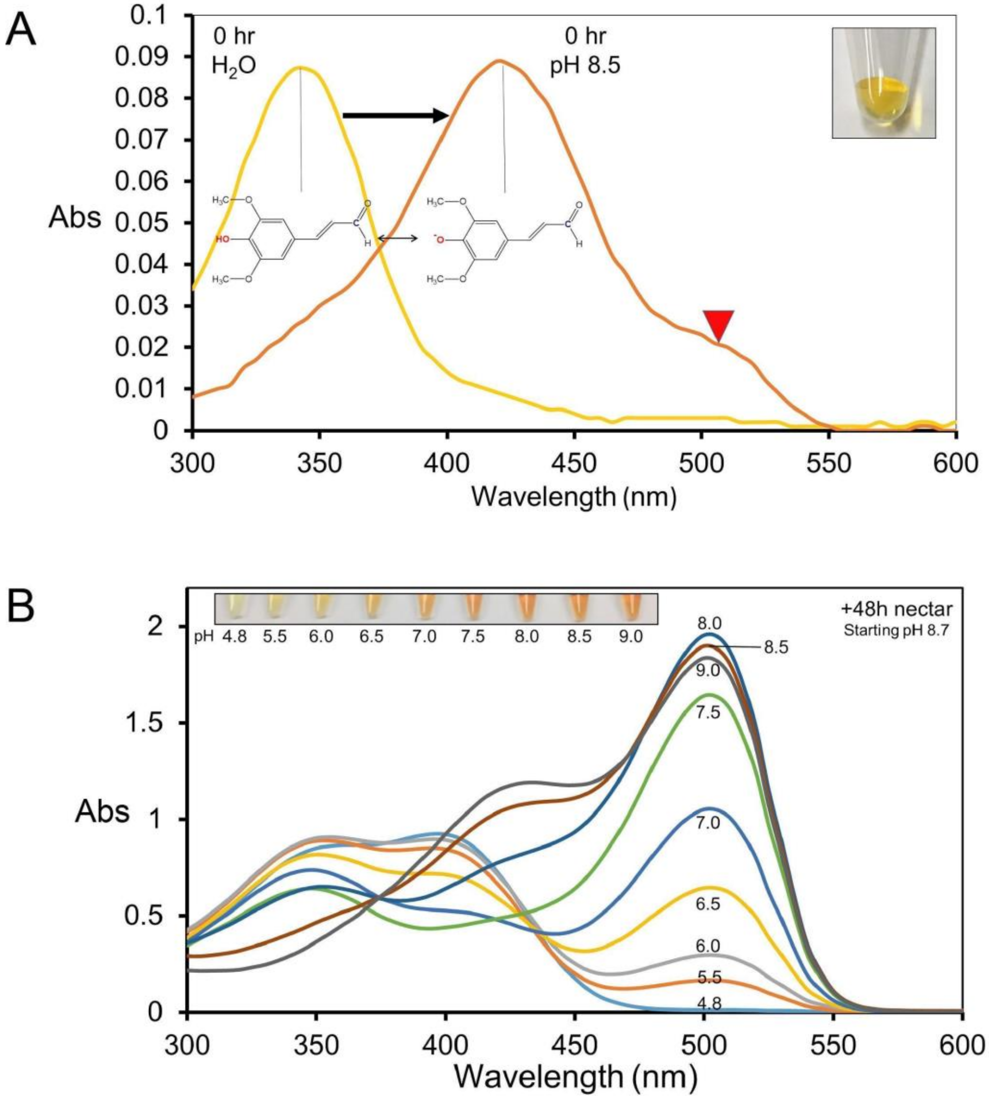


**Fig. S2. Absorbance spectra of yellow (0 h) and red (+48 h) nectar as a function of pH.**

**(A)** Absorbance spectra of 0 h (yellow) nectar after a 1:50 dilution in either water or 50 mM Tricine, pH 8.5. An image of the non-diluted nectar is shown in the inset.

**(B)** Absorbance spectra of +48 h48 h48 h8 h (red) nectar as a function of pH. The absorbance spectra of raw red nectar (+48 h48 h48 h8 h) diluted 1:20 in 50 mM buffers at of varying pH, including: 4.8 (sodium acetate), 5.5 (MES), 6.0 (MES), 6.5 (MES), 7.0 (HEPES), 7.5 (HEPES), 8.0 (HEPES), 8.5 (Tricine), and 9.0 (TAPS).

**Fig. S3**

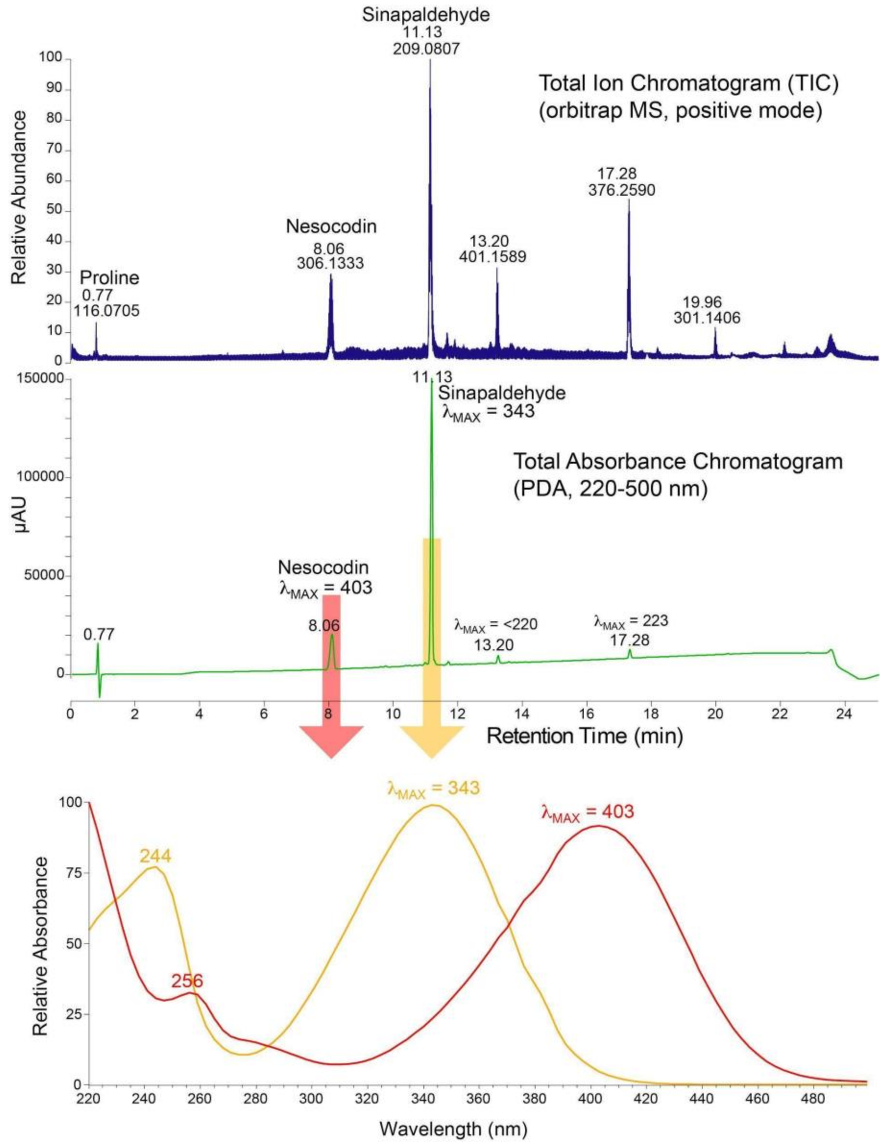


**Fig. S3. LC-MS/PDA analysis of red (+48 h) Nesocodon nectar.** Nectar pigment was first purified by solid phase extraction (SPE) using $C_{18}$-Ziptip (Millipore) and then subjected to $C_{18}$-reversed phase chromatography with in line UV/visible spectroscopy (photodiode array, PDA-detector) followed by mass spectrometric analysis (quadrupole-orbitrap hybrid MS). The top panel shows the total ion chromatogram (TIC) for the analysis with major peaks visible for sinapaldehyde @11.31 min and nesocodin at 8.06 min. The second panel shows the total UV/visible absorbance chromatogram, again with the sinapaldehyde and nesocodin as the major peaks. The bottom panel shows the UV/visible spectra for the nesocodin peak (red) with $\lambda_{MAX}$ = 403 nm and the sinapaldehyde peak (yellow) with $\lambda_{MAX}$ = 343 nm. Since the LC solvents contained 0.1% formic acid the pH was low enough that both compounds maintained fully protonated phenolic hydroxyl groups and the spectra are consistent with spectra of red nectar acidified to pH = 6.0 or lower.

**Fig. S4**

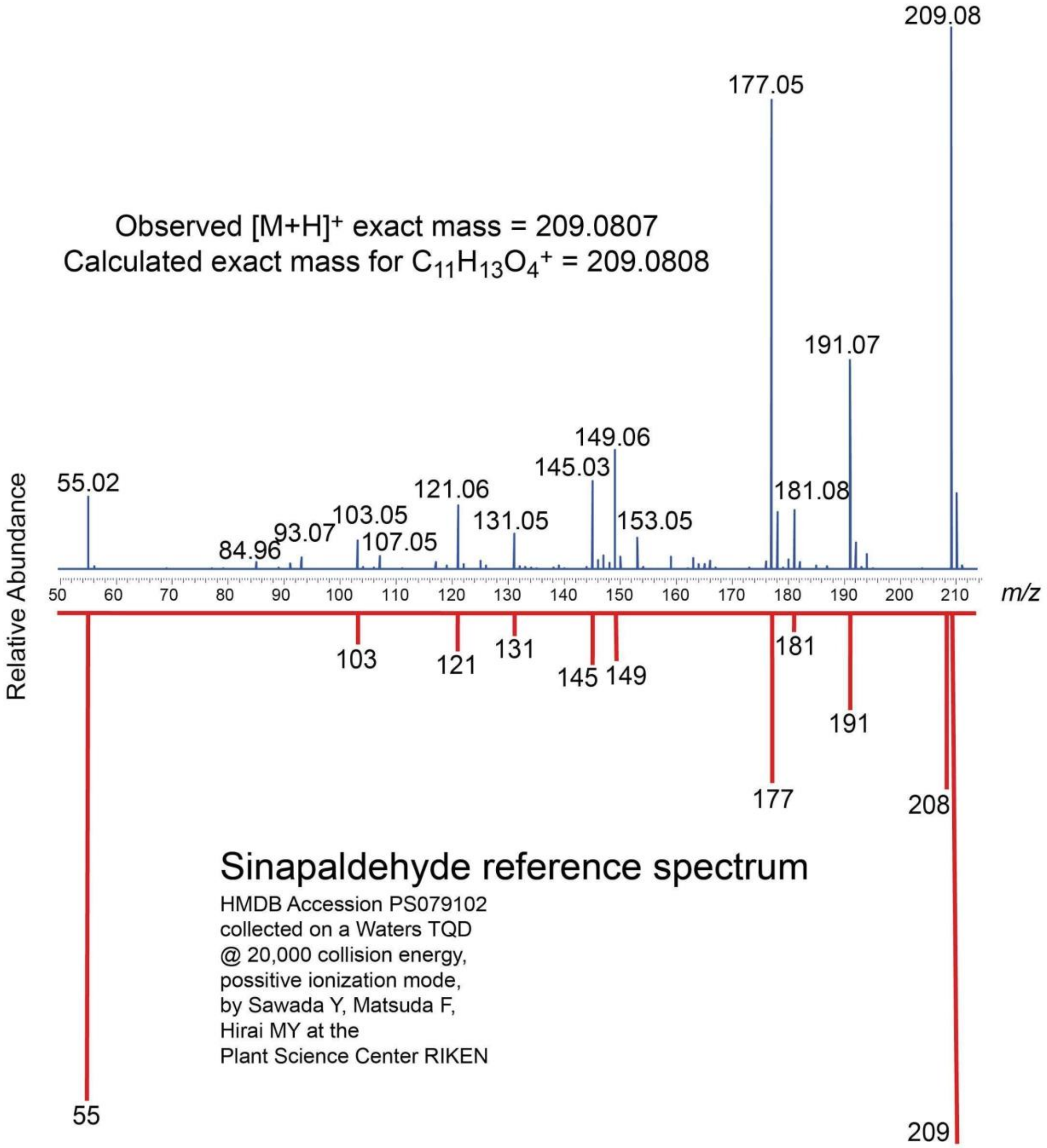


**Fig. S4. Identification of sinapaldehyde in *Nesocodon* nectar by HRMS & MS/MS.**

**Fig. S5**

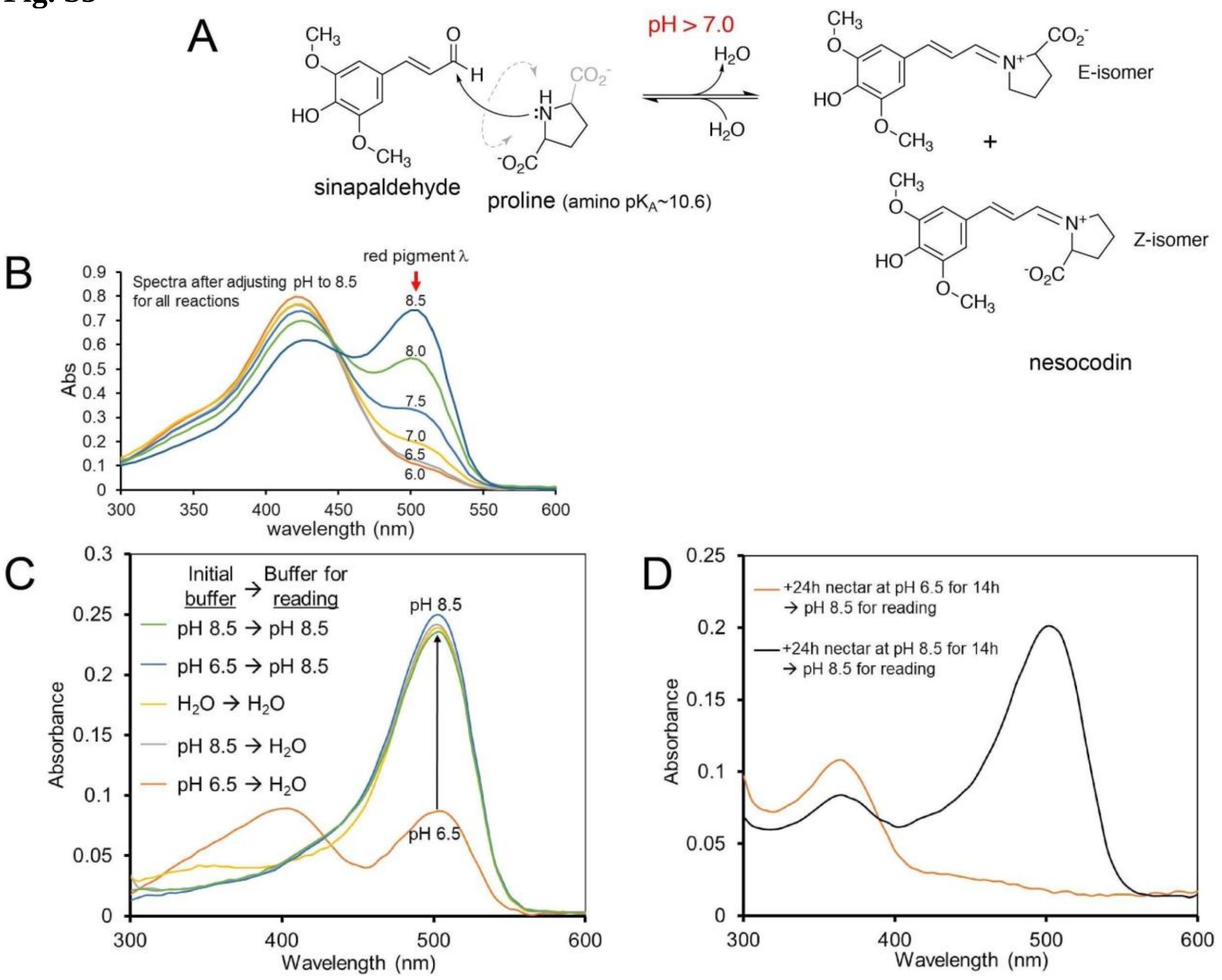


**Fig. S5. An alkaline pH is required for nectar pigment (nesocodin) synthesis and stability.** **(A)** Shows the reaction between sinapaldehyde and proline to generate E- and Z- isomers of nesocodin. The $pK_A$ for the amino group of proline is ~10.6 and so as the pH increases more of the deprotonated (nucleophilic) form of proline is available to react with the aldehyde. The reaction is observable in water due to the color change as the pH is raised above 7.0. **(B)** Absorbance spectra of the reactions from Fig. 1F after adjustment to pH 8.5 via 1:1 dilution with 100 mM Tricine, pH 8.5. **(C)** Alkaline red nectar (+24 h) was diluted 1:19 with 50 mM MES, pH 6.5 and immediately diluted again 1:4 in either di$H_2O$ or 100 mM Tricine, pH 8.5 (i.e., 20 µL of sample in 80 µL buffer) and subjected to UV-Vis spectrophotometric analysis. Control samples similarly diluted stepwise in either di$H_2O$ or 50 mM Tricine, pH 8.5 were included as controls. **(D)** Red nectar (+24 h) was diluted 1:1 with either 100 mM MES, pH 6.5 or 100 mM Tricine, pH 8.5 and incubated in the dark at 21$^{o}$C for 14h. The samples were then diluted 1:19 with 100 mM Tricine, pH 8.5 (i.e., 5 µL of sample into 95 µL buffer) and subjected to UV-Vis spectrophotometric analysis.

**Fig. S6**

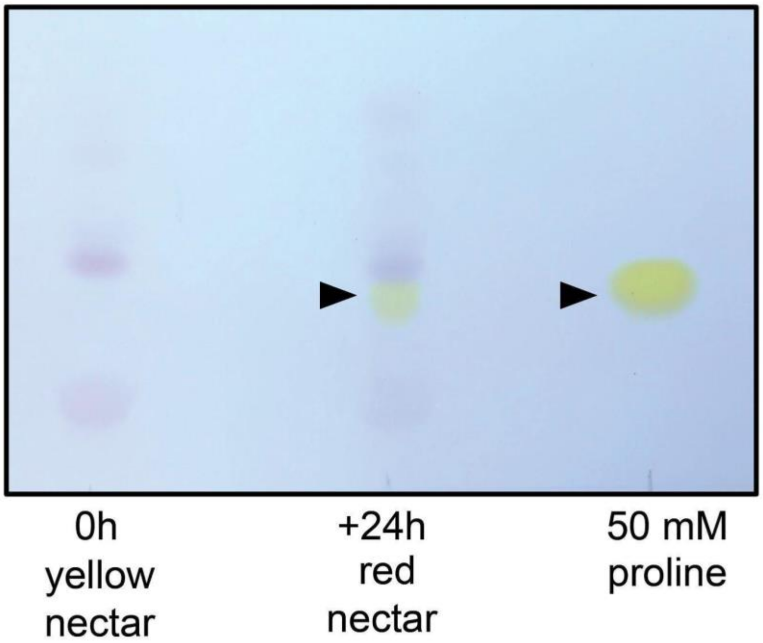


**Fig. S6. Proline in Nesocodin Nectar.** Thin layer chromatographic (TLC) analysis of yellow and red *Nesocodon* nectar was performed alongside a proline standard and visualized using ninhydrin staining. The TLC shows that proline is by far the most abundant amine in red nectar and is not detectable in yellow nectar. In red nectar proline appears to be present in the 10s of mM but this approach is quantitatively approximate. A second method for quantitative analysis of proline was performed using isotope dilution with [$^{13}C_5$]-proline as an internal standard and a GC-MS-based amino acid analysis method performed in triplicate indicated that the proline concentration in red nectar is 66 (±5) mM (data not shown).

**Fig. S7**

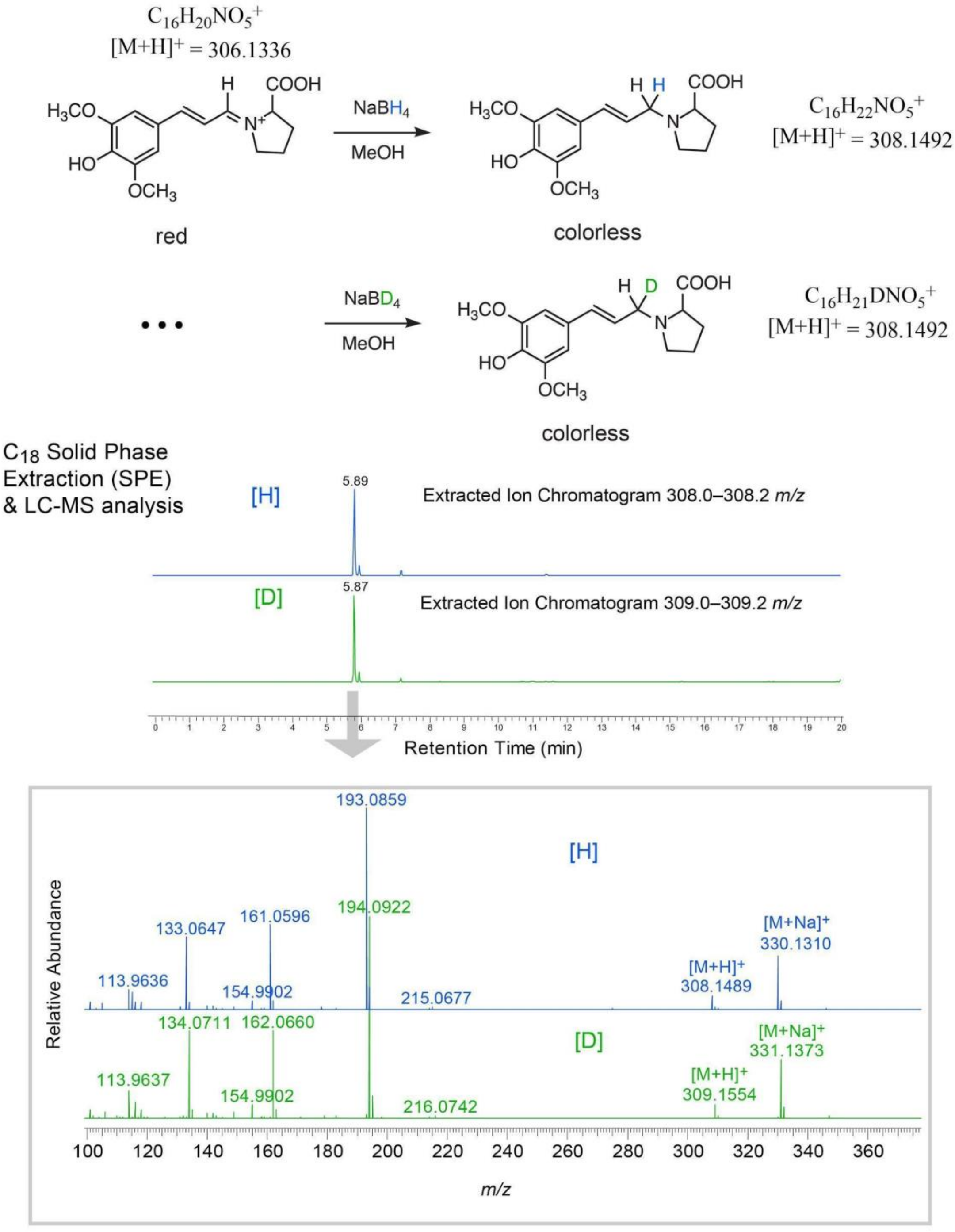


**Fig. S7. Derivatization of synthetic nesocodin by reduction with $NaBH_4$ and $NaBD_4$ followed by LC-MS analysis of products verifies the presence of an imine.**

**Fig. S8.**

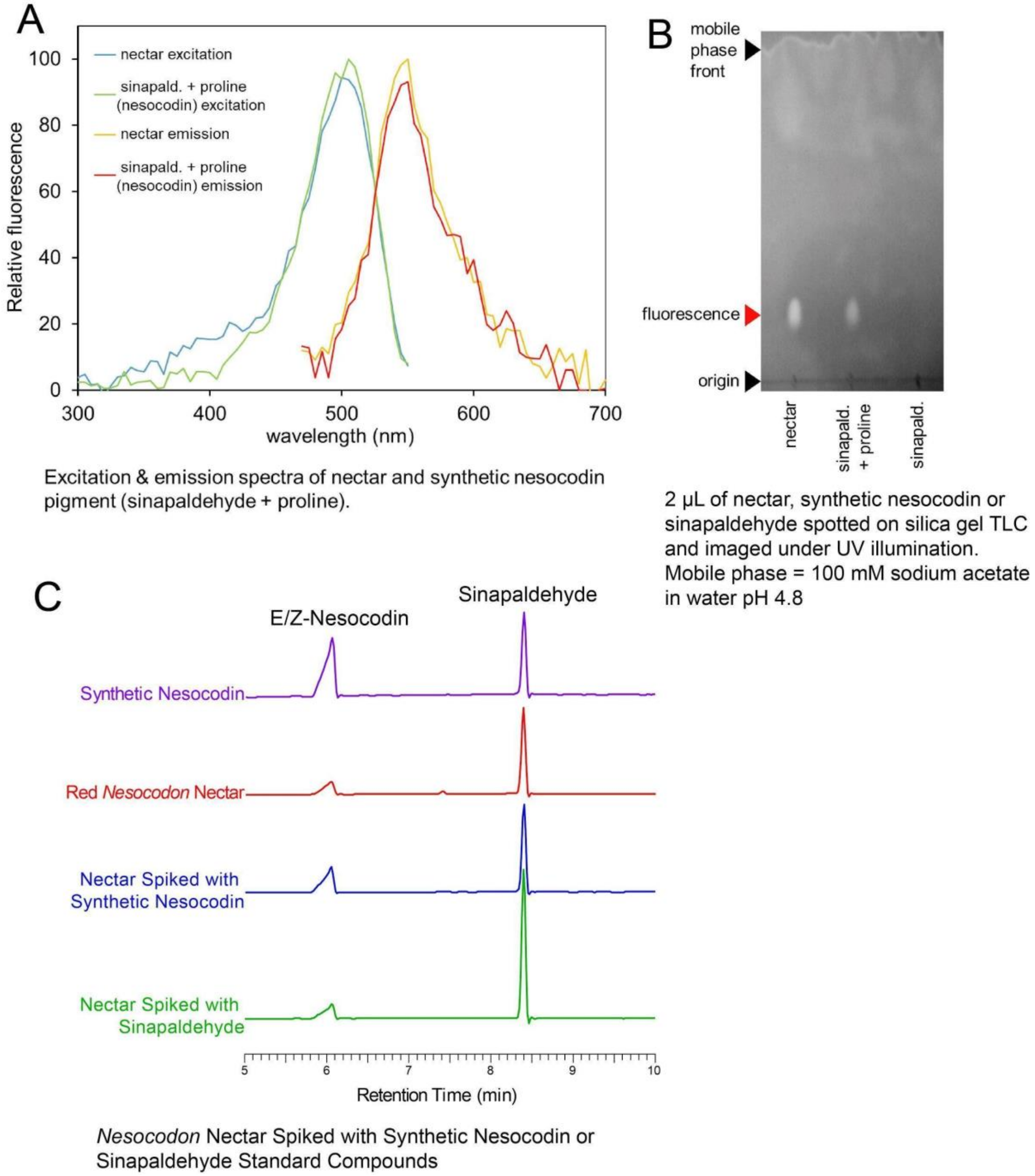


**Fig. S8. Validation of nesocodin pigment synthesis** via **(A)** excitation and emission spectra of synthetic nesocodin (1 mM sinapaldehyde + 10 mM proline in 50 mM Na(H)$CO_3$ pH 9.0) and raw nectar, both diluted 1:99 in 25 mM HEPES, pH 8.0; **(B)** thin-layer chromatography of synthetic nesocodin (sinapaldehyde + proline) and raw nectar (2 µl of each was spotted onto a silica gel TLC plate with a mobile phase of 100 mM sodium acetate, pH 4.8); and **(C)** LC-MS/PDA analyses (5 to 10 min) of 1) synthetic nesocodin, 2) red nesocodon nectar, 3) nectar spiked with synthetic nesocodin, and 4) nectar spiked with sinapaldehyde standard.

**Fig. S9**

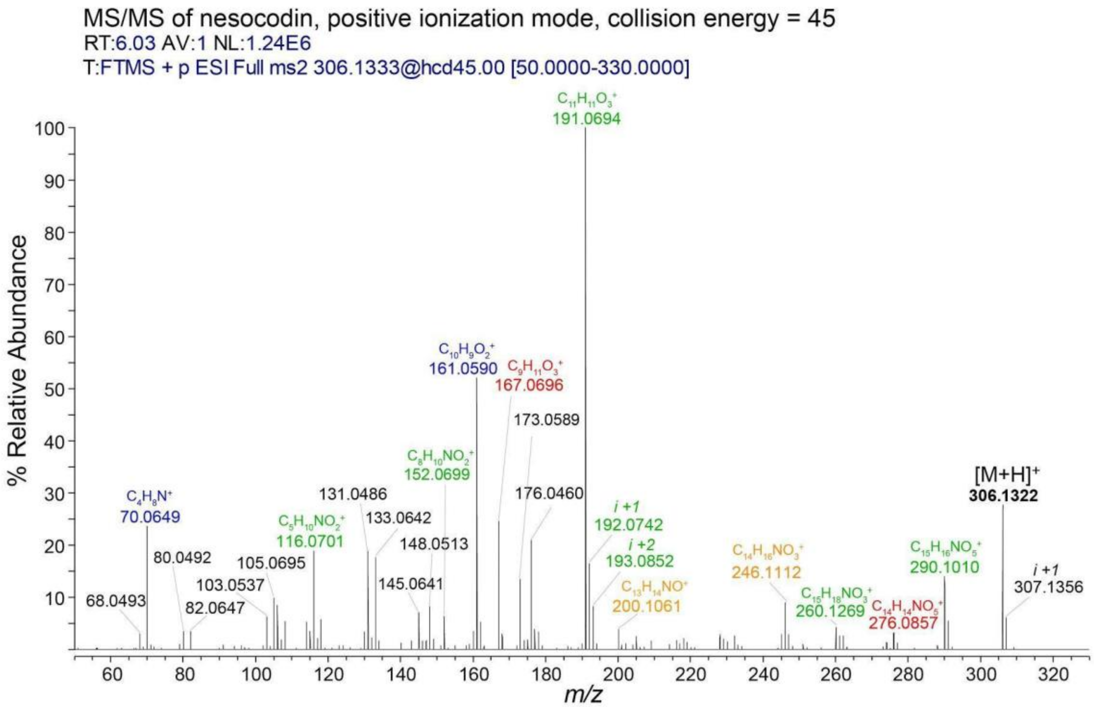


| fragment ion m/z | relative intensity (all peaks > 3) | elemental composition of ion | neutral DBE | difference from molecular ion |
|---|---|---|---|---|
| 68.0493 | 3.14 | $C_4H_6N^+$ | 3 | C12H14O5 |
| 70.0649 | 23.79 | $C_4H_8N^+$ | 2 | C12H12O5 |
| 80.0492 | 3.45 | $C_5H_6N^+$ | 4 | C11H14O5 |
| 82.0647 | 3.8 | $C_5H_9N^+$ | 3 | C11H12O5 |
| 103.0537 | 6.38 | $C_8H_7^+$ | 6 | C8H13NO5 |
| 105.0695 | 9.86 | $C_8H_9^+$ | 5 | C8H11NO5 |
| 106.0695 | 8.73 | $C_7H_8N^+$ | 5 | C9H12O5 |
| 108.0803 | 5.36 | $C_7H_{10}N^+$ | 4 | C9H10O5 |
| 114.0544 | 5.23 | $C_5H_8NO_2^+$ | 3 | C11H12O3 |
| 115.0536 | 3.54 | $C_9H_7^+$ | 7 | C7H13NO5 |
| 116.0701 | 19.67 | $C_5H_{10}NO_2^+$ | 2 | C11H10O3 |
| 118.0407 | 5.96 | $C_4H_8NO_3^+$ | 2 | C12H12O2 |
| 130.0408 | 3.47 | $C_5H_8NO_3^+$ | 3 | C11H12O2 |
| 131.0486 | 18.67 | $C_9H_7O^+$ | 7 | C7H13NO4 |
| 133.0642 | 17.47 | $C_9H_9O^+$ | 6 | C7H11NO4 |
| 145.0641 | 7.29 | $C_{10}H_9O^+$ | 7 | C6H11NO4 |
| 148.0513 | 8.4 | $C_8H_6NO_2^+$ | 7 | C8H14O3 |
| 152.0699 | 6.29 | $C_8H_{10}NO_2^+$ | 5 | C8H10O3 |
| 160.0512 | 3.66 | $C_9H_6NO_2^+$ | 8 | C7H14O3 |
| 161.0590 | 51.72 | $C_{10}H_9O_2^+$ | 7 | C6H11NO3 |
| 167.0696 | 24.56 | $C_9H_{11}O_3^+$ | 5 | C7H9NO2 |
| 173.0589 | 13.7 | $C_{11}H_9O_2^+$ | 8 | C5H11NO3 |
| 176.0460 | 21.44 | $C_9H_6NO_3^+$ | 8 | C7H14O2 |
| 177.0526 | 3.96 | $C_{10}H_9O_3^+$ | 7 | C6H11NO2 |
| 191.0694 | 100 | $C_{11}H_{11}O_3^+$ | 7 | C5H9NO2 |
| 200.1061 | 3.87 | $C_{13}H_{14}NO^+$ | 8 | C3H6O4 |
| 246.1112 | 9.35 | $C_{14}H_{16}NO_3^+$ | 8 | C2H4O2 |
| 260.1269 | 4.36 | $C_{15}H_{18}NO_3^+$ | 8 | CH2O2 |
| 276.0857 | 3.46 | $C_{14}H_{14}NO_5^+$ | 9 | C2H6 |
| 290.1010 | 13.96 | $C_{15}H_{16}NO_5^+$ | 9 | CH4 |
| **306.1322** | 28.99 | $C_{16}H_{20}NO_5^+$ | 8 | - |

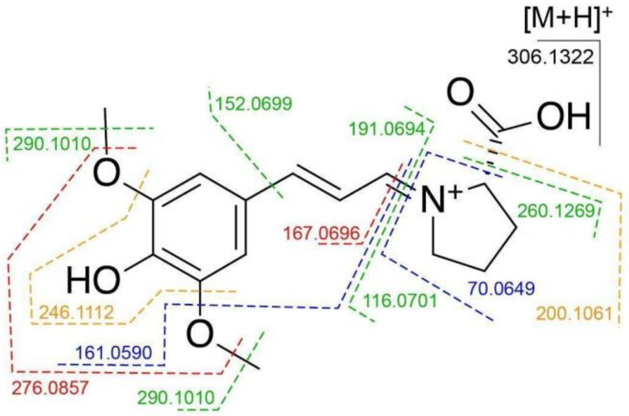


**Fig. S9. MS/MS spectrum of nesocodin with fragmentation assigned.**

**Fig. S10**

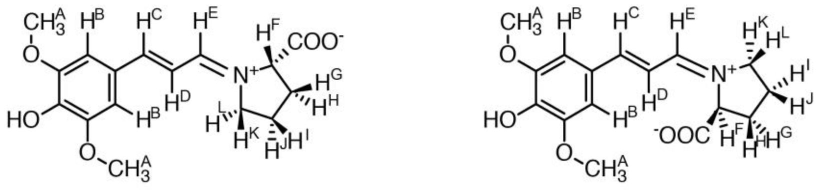

*E*-nesocodin (64%)    *Z*-nesocodin (36%)

Summary of 1D and 2D $^1$H-NMR resonances for both E and Z isomers of nesocodin (600 MHz) in methanol-*$d_4$*

| Proton(s) (*isomer*) | 1D $^1$H-NMR | DQF-COSY indicated couplings |
|---|---|---|
| $H^A$ (*E*) | d 3.82 (s, 3H, $CH_3$) | none |
| $H^A$ (*Z*) | d 3.80 (s, 3H, $CH_3$) | none |
| $H^B$ (*E*) | d 6.99 (broad s, 2H) | none |
| $H^B$ (*Z*) | d 6.92 (broad s, 2H) | none |
| $H^C$ (*E*) | d 7.47 (d, 1H, J=14 Hz) | d 6.47 ($H^D$) |
| $H^C$ (*Z*) | d 7.42 (d, 1H, J=14 Hz) | d 6.26 ($H^D$) |
| $H^D$ (*E*) | d 6.47 (dd, 1H, J=12, 14 Hz) | d 7.83, 7.47 ($H^C$, $H^D$) |
| $H^D$ (*Z*) | d 6.26 (dd, 1H, J=12, 14 Hz) | d 7.87, 7.42 ($H^C$, $H^D$) |
| $H^E$ (*E*) | d 7.83 (d, 1H, J=12 Hz) | d 6.47, 4.41, 3.80, 3.70 ($H^D$, $H^F$, $H^K$, $H^L$) |
| $H^E$ (*Z*) | d 7.87 (d, 1H, J=12 Hz) | d 6.26, 4.35, 3.94 ($H^D$, $H^H$, $H^L$) |
| $H^F$ (*E*) | d 4.41 (dd, 1H, J=5, 8 Hz) | d 7.83, 2.31, 2.25 ($H^E$, $H^G$, $H^H$) |
| $H^F$ (*Z*) | d 4.35 (dd, 1H, J=5, 8 Hz) | d 7.87, 2.44, 2.22 ($H^E$, $H^G$, $H^H$) |
| $H^G$ (*E*) | d 2.31 (=13, 8 Hz) | d 4.41, 3.80, 3.70, 2.25 ($H^F$, $H^L$, $H^K$, $H^H$) |
| $H^G$ (*Z*) | d 2.44 (=13, 8 Hz) | d 4.35, 3.94, 3.82, 2.22, 1.98 ($H^F$, $H^L$, $H^K$, $H^I$, $H^J$) |
| $H^H$ (*E*) | d37 Hz | d 4.41, 2.31, 2.05 ($H^F$, $H^G$, $H^{I,J}$) |
| $H^H$ (*Z*) | d 2.22 37 Hz | d 4.35, 2.44, 2.09, 1.98 ($H^F$, $H^G$, $H^I$, $H^J$) |
| $H^{I,J}$ (*E*) | d 2.05 (, 2H) | d 3.80, 3.70, 2.25 ($H^L$, $H^K$, $H^H$) |
| $H^I$ (*Z*) | d 2.09 (, 1H) | d 3.94, 3.82, 2.22, 1.98 ($H^L$, $H^K$, $H^H$, $H^J$) |
| $H^J$ (*Z*) | d 1.98 (tq, 1H, 12, 7) | d 3.94, 3.82, 2.44, 2.22, 2.09 ($H^L$, $H^K$, $H^G$, $H^H$, $H^I$) |
| $H^K$ (*E*) | d 3.70 (dt, 1H, J=13, 8 Hz) | d 3.80, 2.05 ($H^L$, $H^{I,J}$) |
| $H^K$ (*Z*) | d 3.82 (dt, 1H, J=12, 7 Hz) | d 3.94, 2.09, 1.98 ($H^L$, $H^I$, $H^J$) |
| $H^L$ (*E*) | d 3.80 (dt, 1H, J=13, 8 Hz) | d 3.70, 2.05 ($H^K$, $H^{I,J}$) |
| $H^L$ (*Z*) | d 3.94 (dt, 1H, J=12, 7 Hz) | d 3.82, 2.09, 1.98 ($H^K$, $H^I$, $H^J$) |

**Fig. S10. $^1$H-NMR assignments for synthetic nesocodin.** Structures of E- and Z- isomers of nesocodin are shown with protons labeled ‘A’ through ‘L’ for assignment of $^1$H-NMR data.

**Fig. S11**

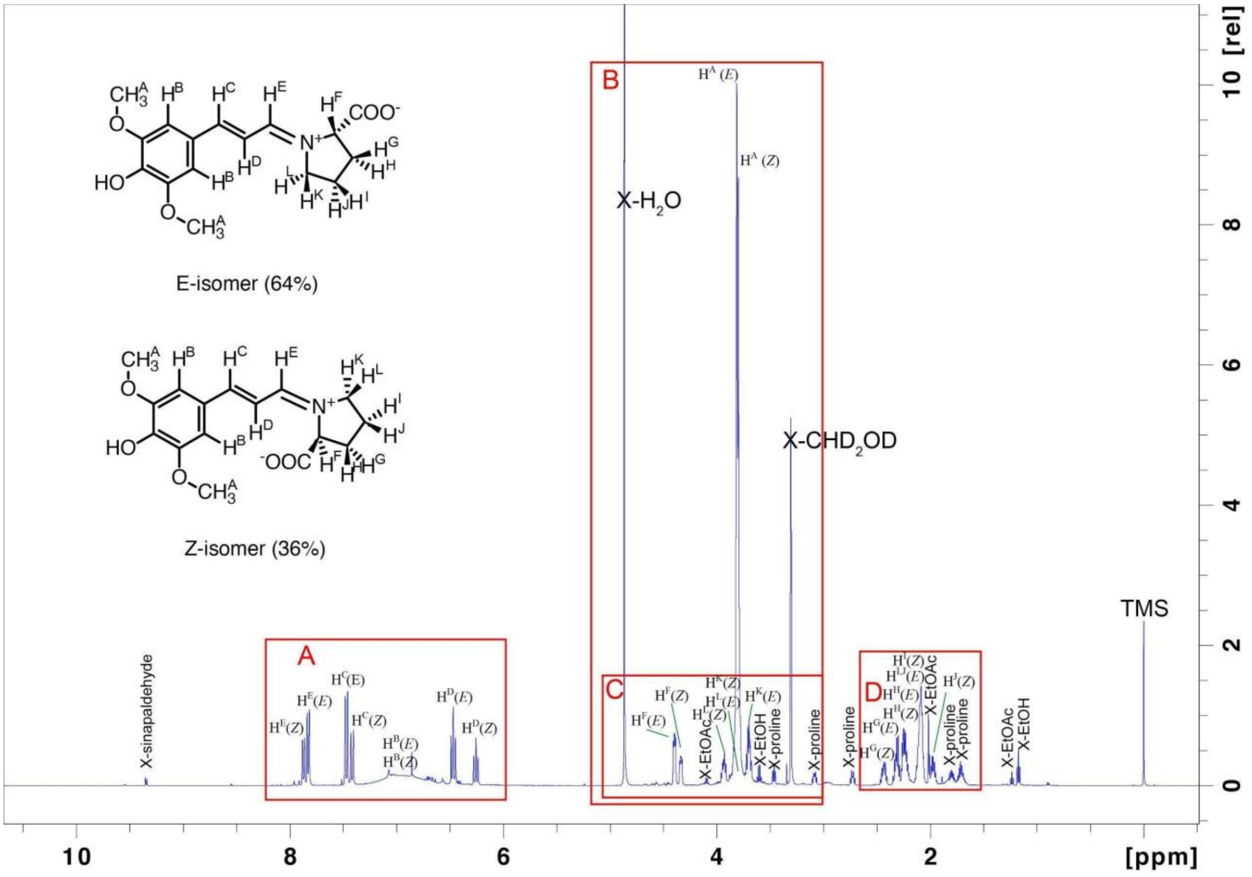

**Fig. S11. 1D-$^1$H-NMR spectrum (600 MHz) of E- and Z-nesocodin in methanol-$d_4$.** Subsequent **Figs. S11A-D** show the specified regions of the spectrum outlined in red boxes on the full spectrum in more detail.

**Fig. S11A**

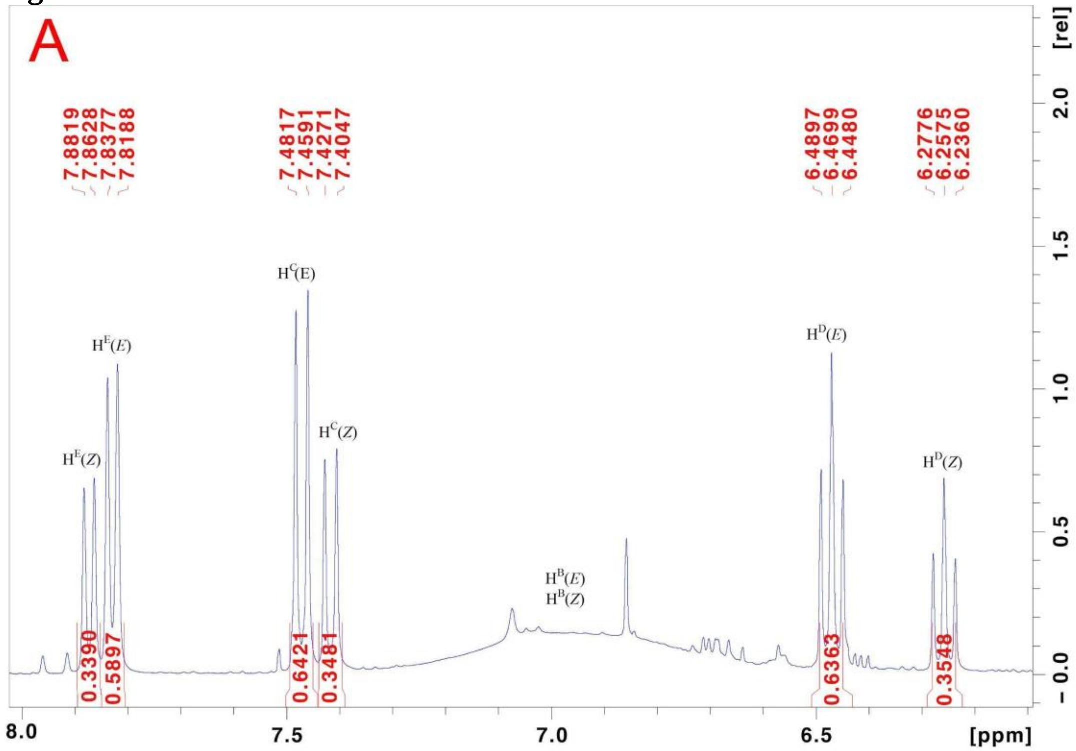


**Fig. S11B**

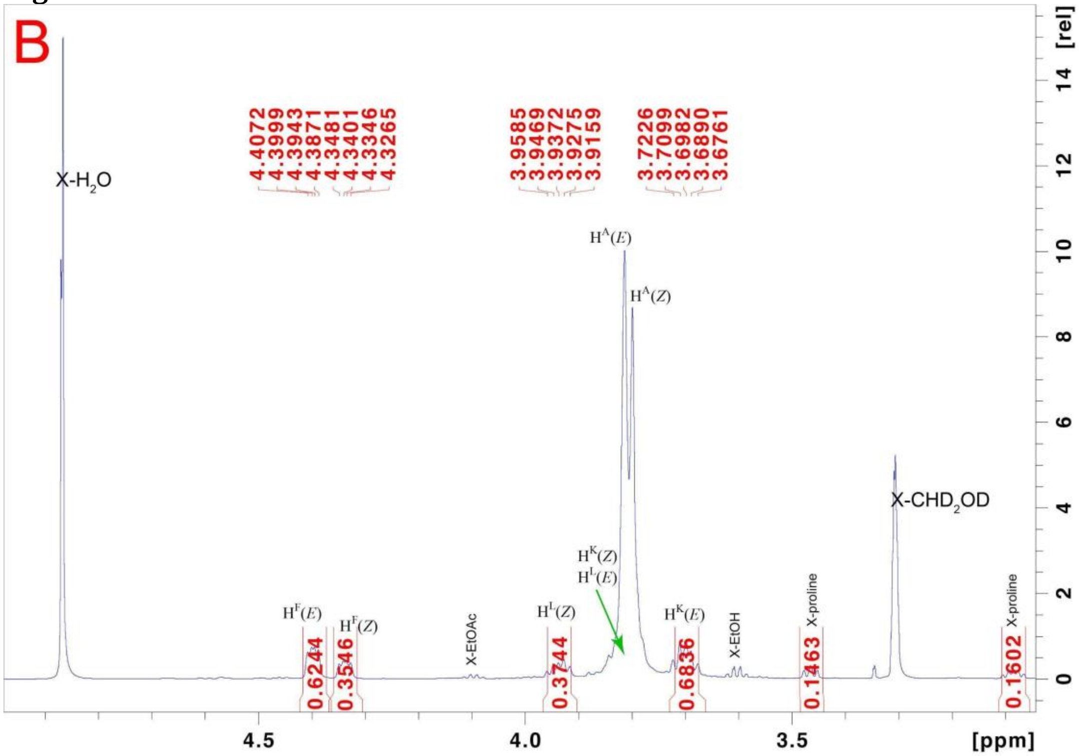

Fig. S11C

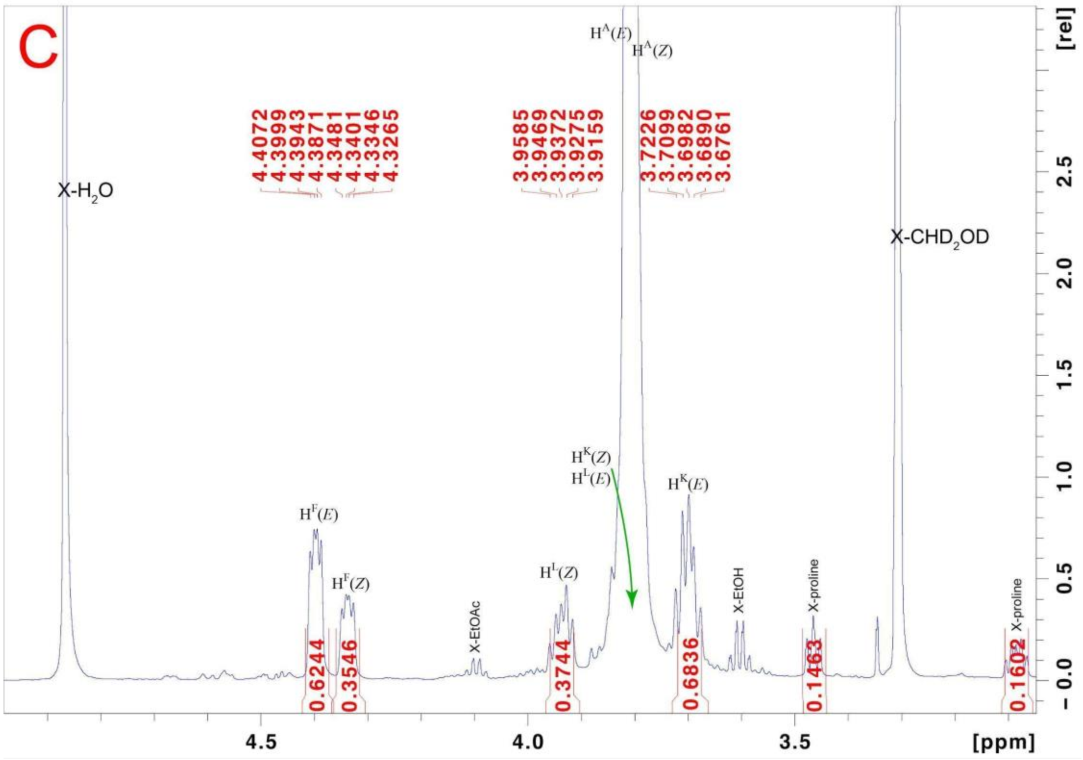


Fig. S11D

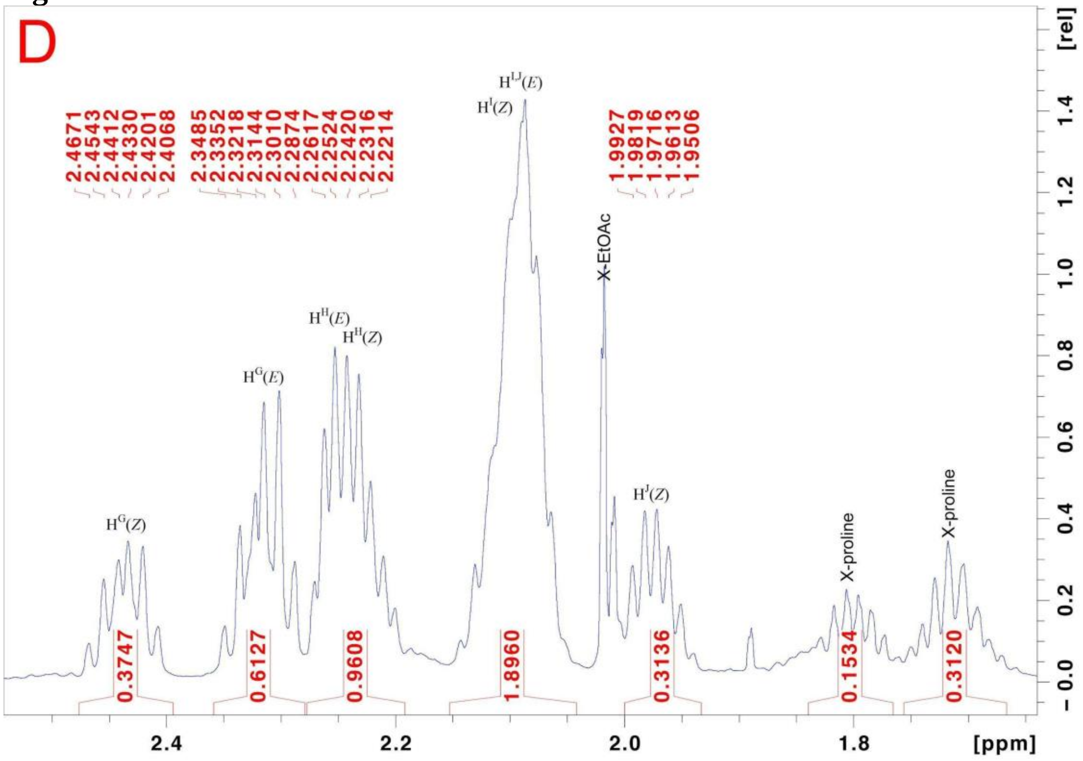

**Fig. S12**

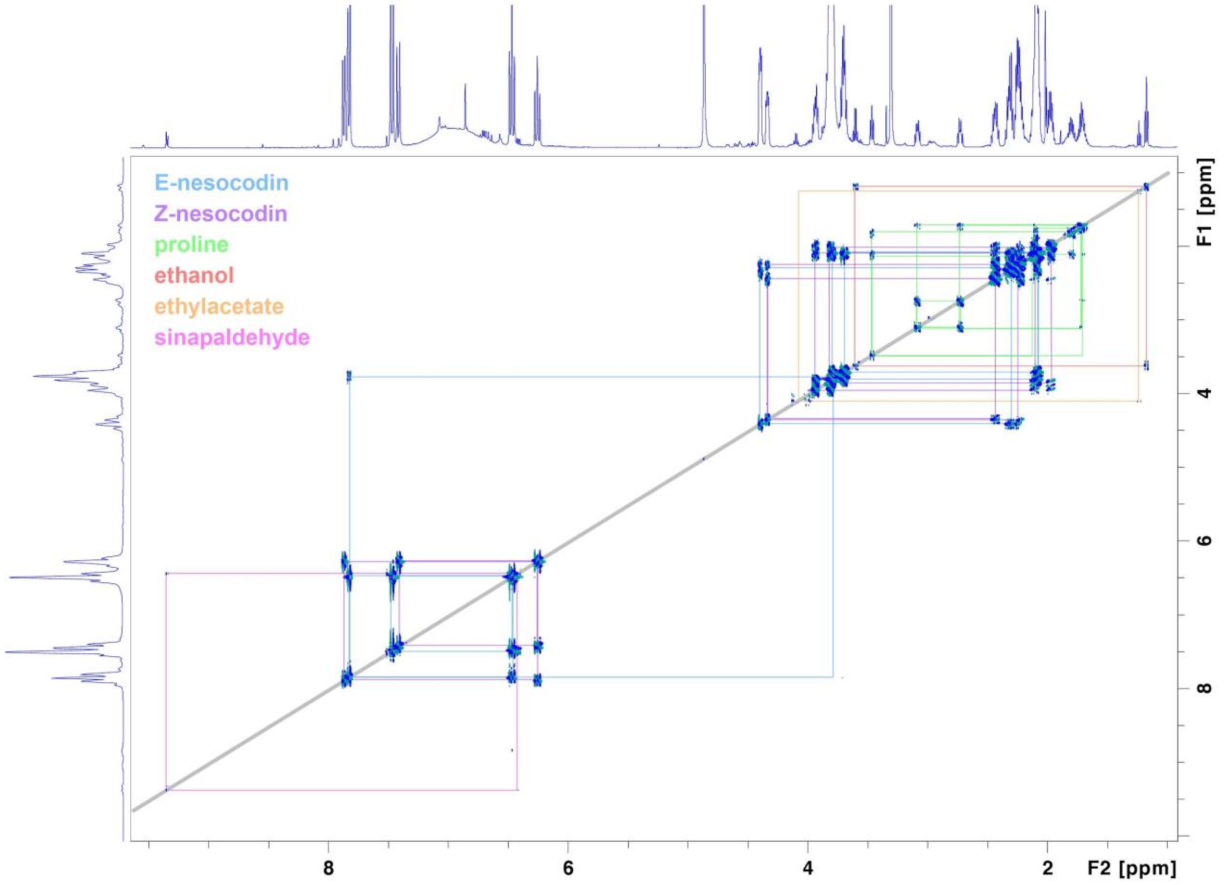


**Fig. S12. 2D DQF-COSY (Double Quantum Filtered-COrrelation SpectroscopY) $^{1}$H-NMR analysis (600 MHz) of *E*- and *Z*-nesocodin in methanol-$d_4$.** Subsequent sub-Figures show additional details: **Fig. S12Key** shows the regions A-L of the spectrum outlined in red boxes in the full spectrum that are given in more detail in the following **Figs. S12A-E**. Long range coupling (4J) between the imine proton HE and proton HF, HK and HL on the proline moiety are most easily seen in panel B. These were used to assign the E- and Z-isomers in the NMR spectrum as the E-isomer has the largest coupling between HE and HK and HL, while the Z-isomer has the largest coupling between HE and HF. The largest 4J coupling constants would be expected as the dihedral angle between the protons approaches 180˚, as is consistent with our structural assignments. The assignment is also consistent with the Z-isomer having lower abundance due to steric interactions between the prolyl carboxylate moiety and proton HD that would be alleviated in the E-isomer.

**Fig. S12 Key**

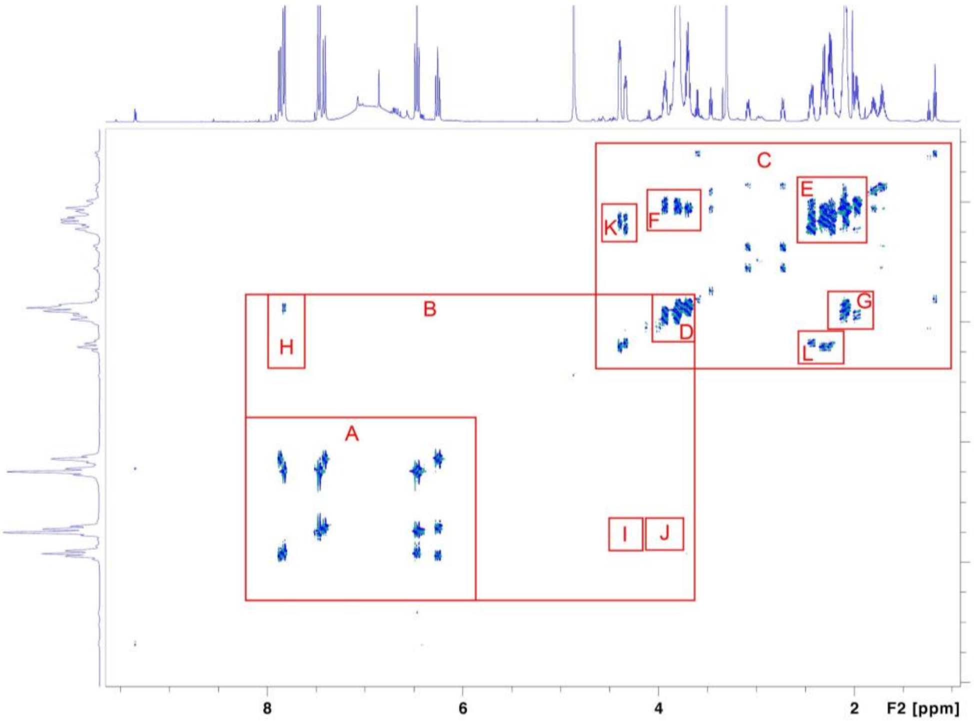


**Fig. S12A**

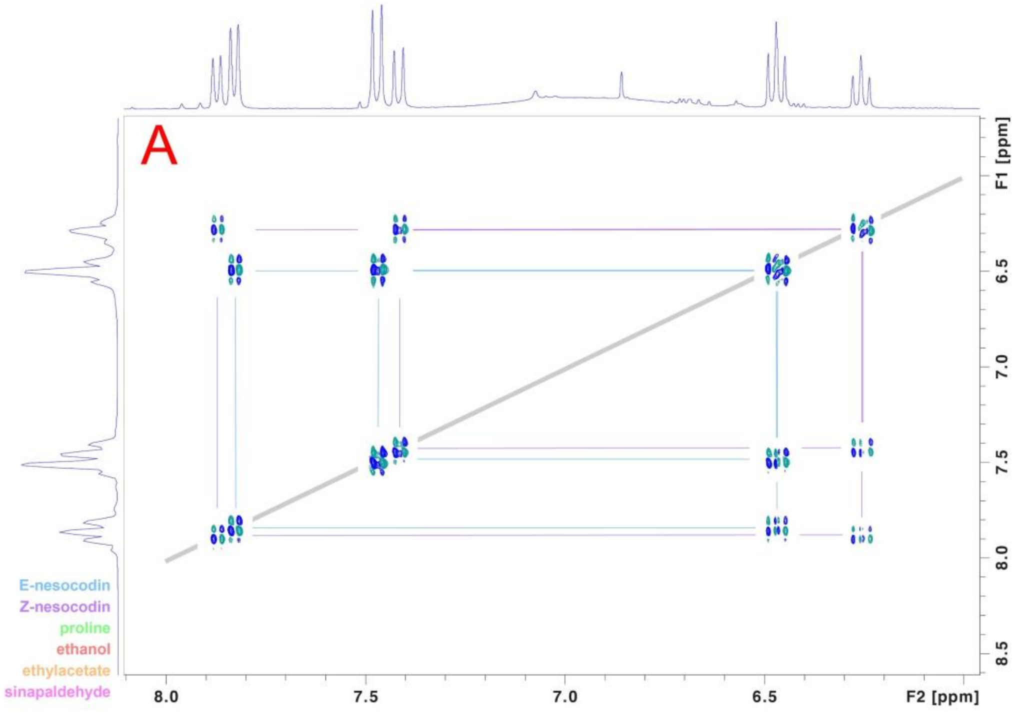

**Fig. S12B**

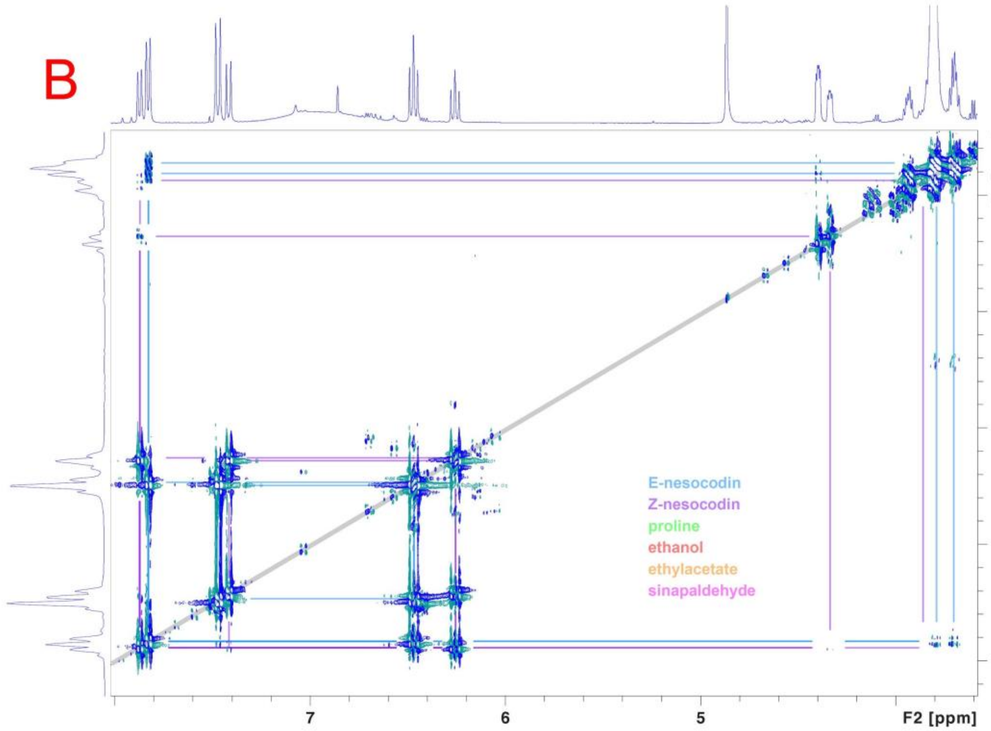


**Fig. S12C**

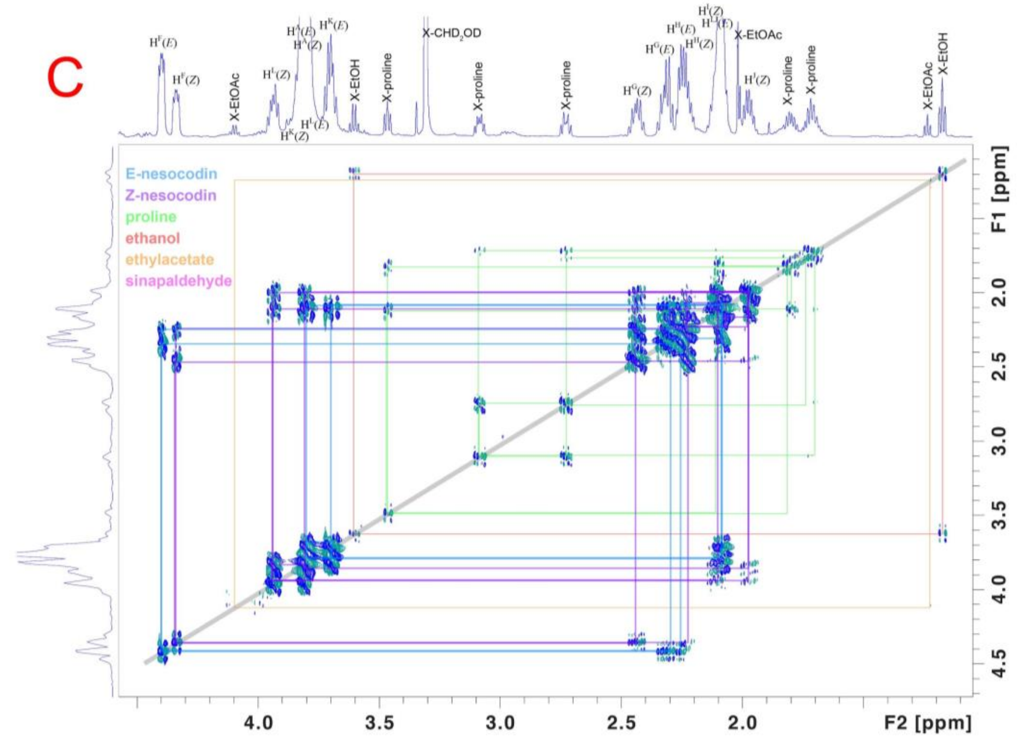

**Fig. S12D**

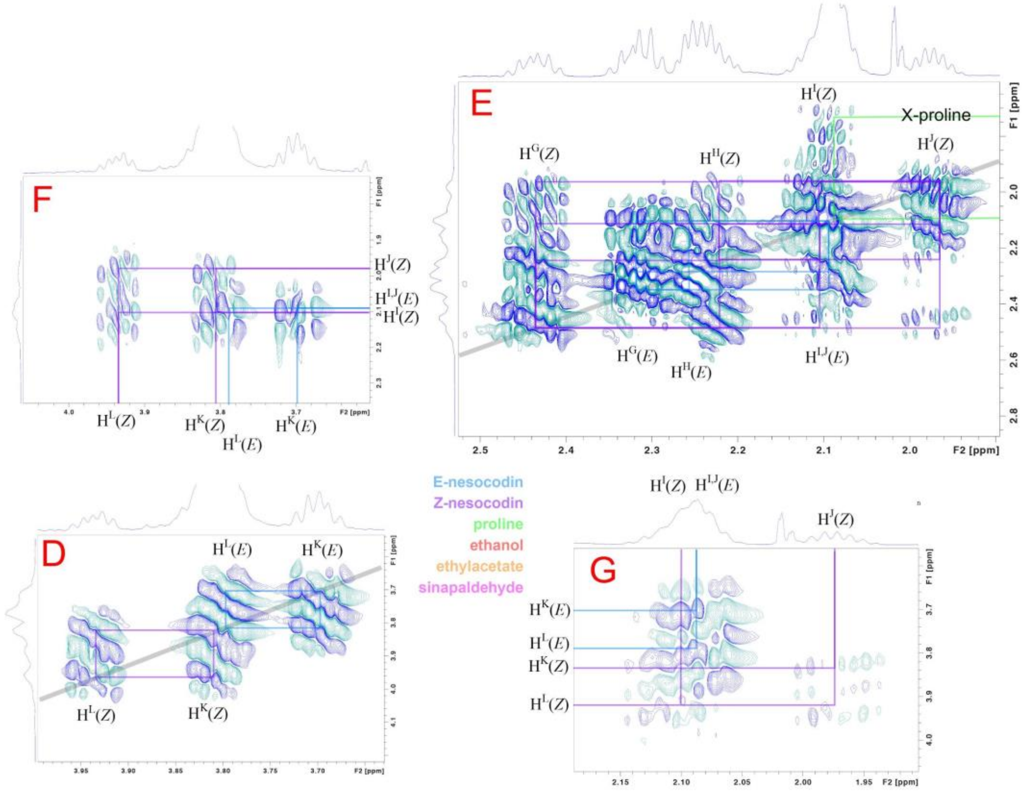


**Fig. S12E**

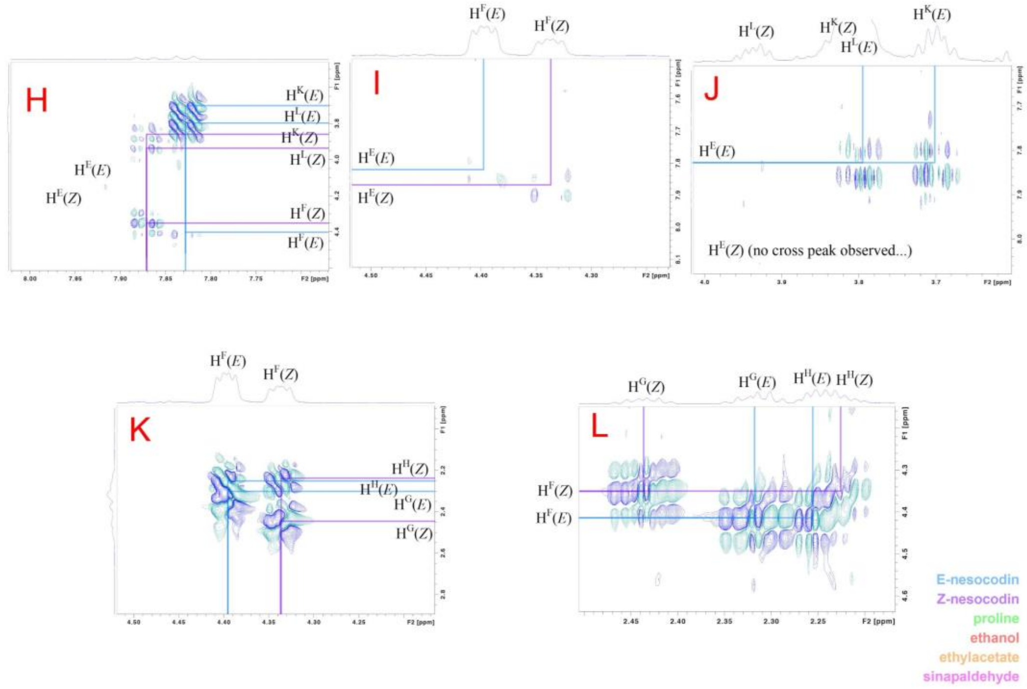

**Fig. S13**

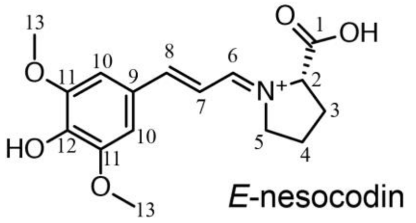


Summary of 1D $^{13}$C-NMR resonances for both E and Z isomers of nesocodin (600 MHz) in methanol-$d_4$

| Proton(s) (*isomer*) | 1D $^{13}$C-NMR proton decoupled | 1D $^{13}$C-NMR proton coupling |
|---|---|---|
| $C^1$ (*E*) | d 176.78 | d, 4Hz |
| $C^1$ (*Z*) | d 176.90 | d, 4Hz |
| $C^2$ (*E*) | d 70.19 | dt, 146, 4Hz |
| $C^2$ (*Z*) | d 66.16 | dt, 146, 4Hz |
| $C^3$ (*E*) | d 31.46 | q, 131, 4Hz |
| $C^3$ (*Z*) | d 32.34 | q, 131, 4Hz |
| $C^4$ (*E*) | d 24.76 | tm, 132, 4Hz |
| $C^4$ (*Z*) | d 24.94 | tm, 132, 4Hz |
| $C^5$ (*E*) | d 50.60 | tt, 143, 4Hz |
| $C^5$ (*Z*) | d 56.08 | tt, 140, 4Hz |
| $C^6$ (*E*) | d 160.57 | dd, 172, 7Hz |
| $C^6$ (*Z*) | d 160.52 | dd, 172, 7Hz |
| $C^7$ (*E*) | d 159.27 | dt, 151, 5Hz |
| $C^7$ (*Z*) | d 159.22 | dt, 151, 5Hz |
| $C^8$ (*E*) | d159.83 | dd, 151, 5Hz |
| $C^8$ (*Z*) | d 159.77 | dd, 151, 5Hz |
| $C^9$ (*E*) | d 166.63 | t, 6Hz |
| $C^9$ (*Z*) | d 167.11 | t, 6Hz |
| $C^{10}$ (*E*) | d 106.20 | dd, 158, 3 Hz |
| $C^{10}$ (*Z*) | d 106.01 | dd, 158, 3 Hz |
| $C^{11}$ (*E*) | d 118.76 | d, 4Hz |
| $C^{11}$ (*Z*) | d 118.66 | d, 4Hz |
| $C^{12}$ (*E*) | d 161.52 | s |
| $C^{12}$ (*Z*) | d 161.49 | s |
| $C^{13}$ (*E*) | d 56.22 | q, 143.7 Hz |
| $C^{13}$ (*Z*) | d 56.20 | q, 143.7 Hz |

**Fig. S13. Supplemental Figure 13. $^{13}$C-NMR Assignments for Synthetic Nesocodin.** Structure of nesocodin is shown with carbon atoms labeled 1 through 13 for assignment of $^{13}$C-NMR data.

Fig. S14

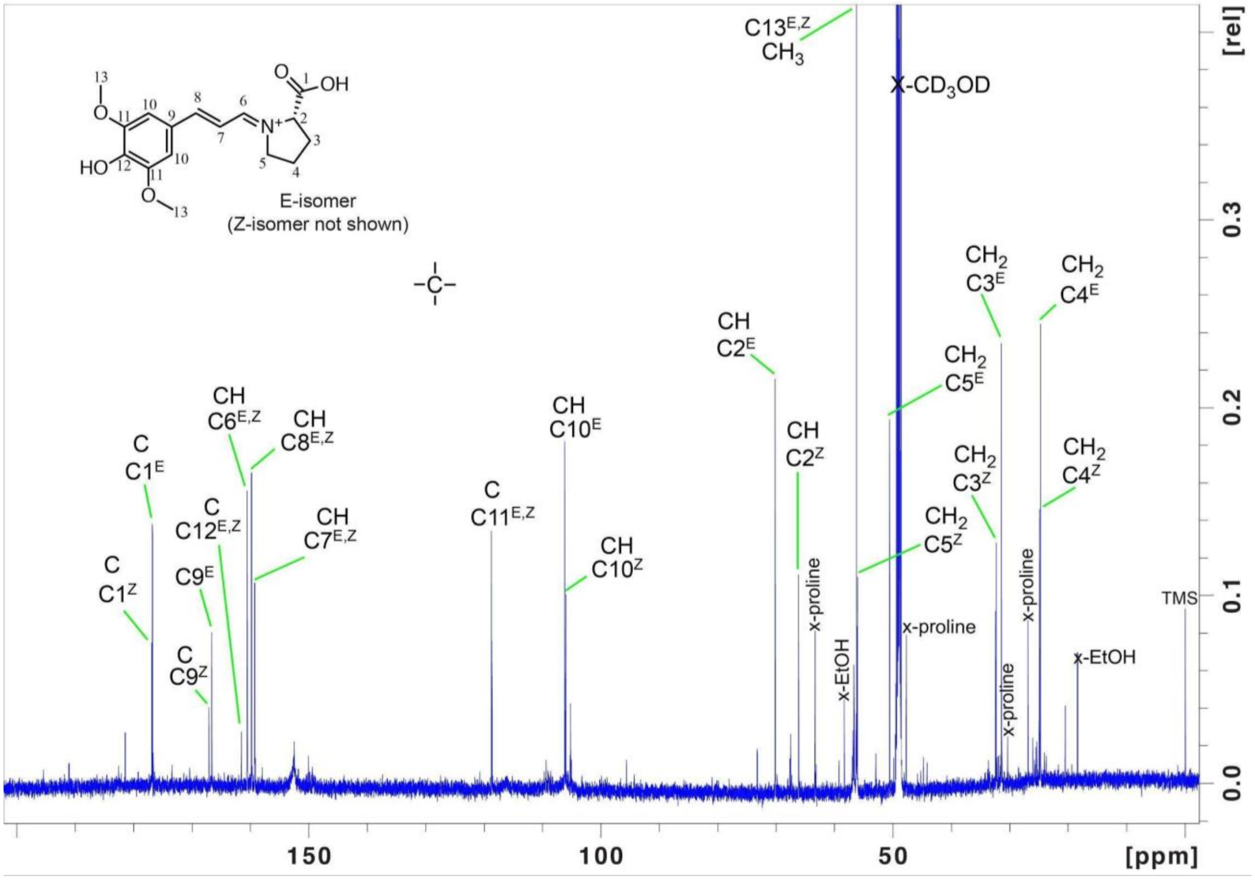


**Fig. S14.** **1D [13]C-NMR spectrum of E- and Z-nesocodin in methanol-*d*$_4$.**

**Fig. S15**

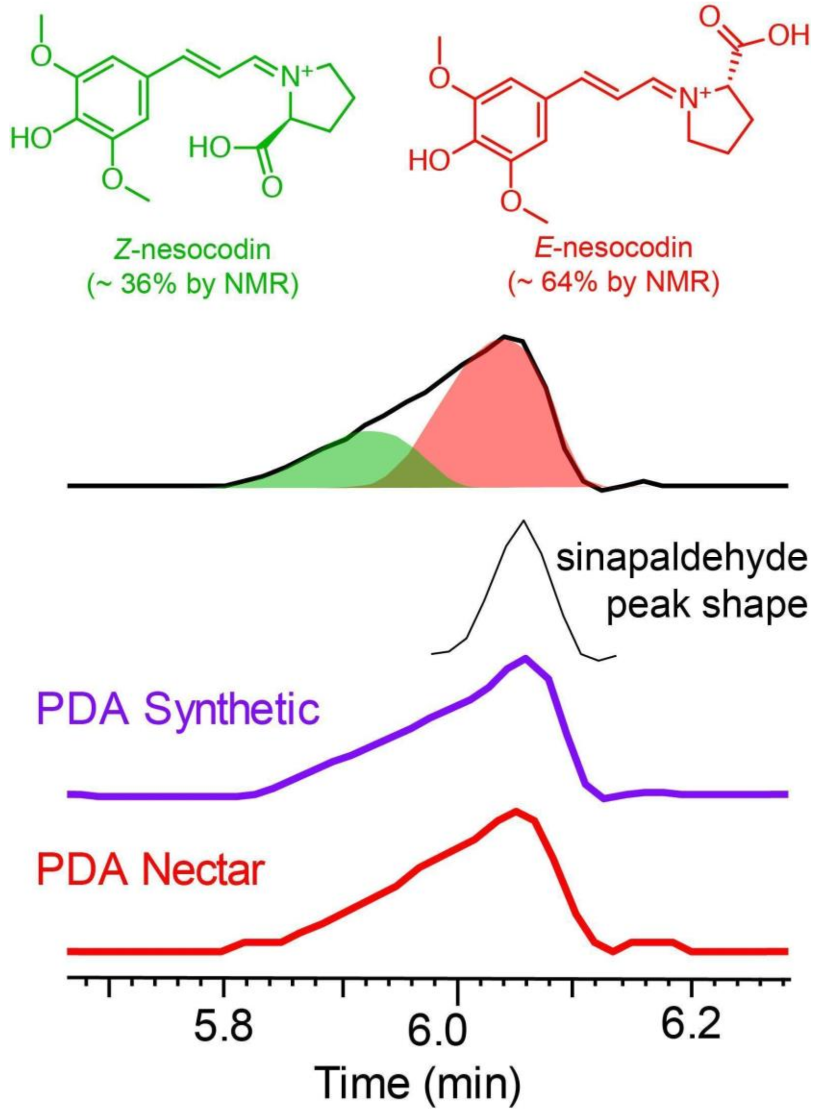


**Fig. S15. LC-PDA of nesocodin isomers comparing *Nesocodon* nectar and synthetic nesocodin.** LC-PDA data are consistent with NMR indicating that E- and Z-nesocodin isomers are present in a 64 to 36 (nearly 2 to 1) ratio (respectively) when produced synthetically. The sinapaldehyde peak shape, which was observable several minutes later during the same linear elution gradient, was used to model the E- (red) and Z- (green) nesocodin isomer peaks to match the observed isomer combination peak. The floral pigment also appears to exist as a 2:1 mixture of E- and Z- isomers based on the presence of two closely eluting peaks in the LC-MS analysis. This would be expected if the proline/sinapaldehyde reaction is non-enzymatic but occurs spontaneously as the pH increases during nectar maturation.

**Fig. S16**

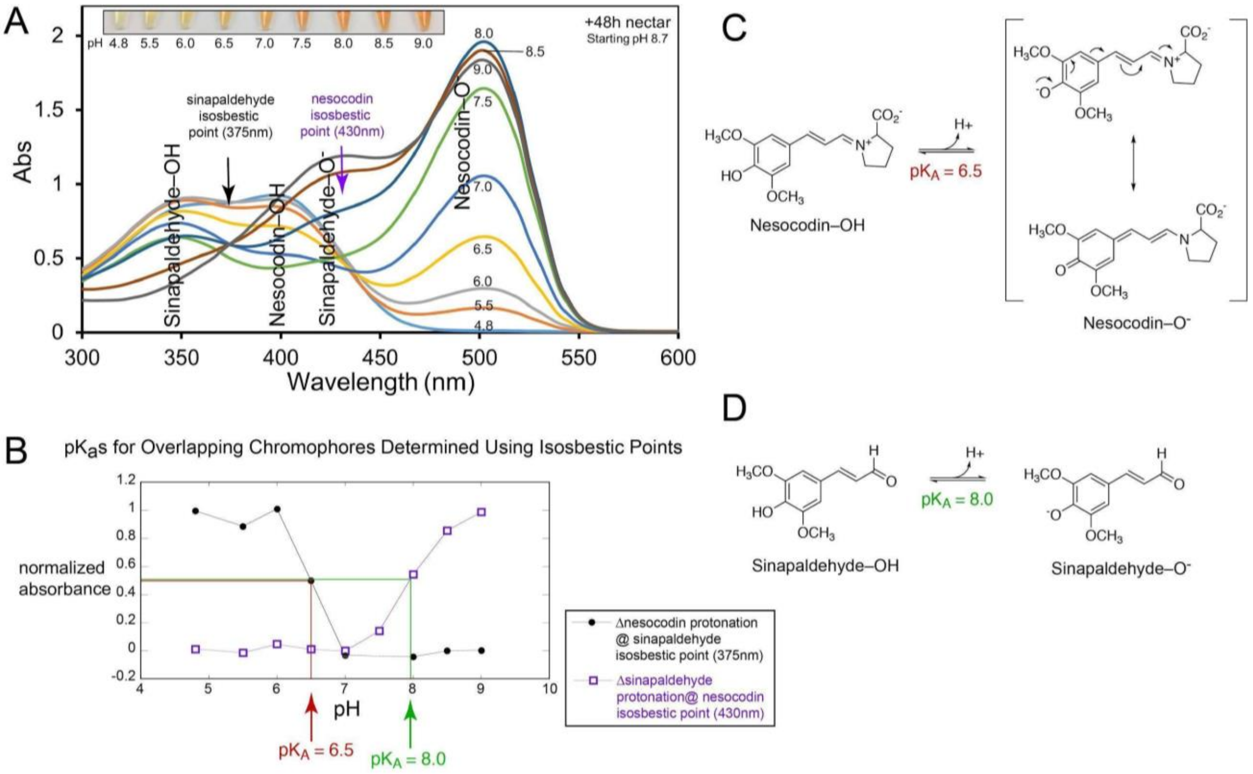


**Fig. S16. The pH dependence of *Nesocodon* nectar pigments nesocodin and sinapaldehyde.** Data from Fig. S2B showing UV/visible spectra of red *Nesocodon* nectar (+48 h nectar, starting at unbuffered pH = 8.7) as the pH is varied by addition to buffer ranging in pH from 4.8 to 9.0 were reanalyzed for $pK_A$ determination. **(A)** shows a pile up of the UV/visible spectra from 300 nm to 600 nm at pHs 4.8, 5.5, 6.0, 6.5, 7.0, 7.5, 8.0, 8.5 and 9.0. Isosbestic points are apparent at 375 nm for the protonation/deprotonation of sinapaldehyde and 430 for the protonation/deprotonation of nesocodin. **(B)** shows a plot of the absorbance at these isosbestic points normalized to range between 1 and 0. By observing the spectral changes at the isosbestic points it is possible to remove the pH dependent contributions from the overlapping chromophores. The nesocodin $pK_A$ can be measured at the sinapaldehyde isosbestic point by following the pH dependent decrease in the protonated nesocodin as the pH increases, and the sinapaldehyde pKa can be measured at the nesocodin isosbestic point by following the pH dependent increase in the deprotonated sinapaldehyde as the pH increases. Using this approach the nesocodin appears to have a $pK_A$ of 6.5 and the sinapaldehyde has a $pK_A$ of 8.0. The lower $pK_A$ for nesocodin is consistent with the added resonance delocalization of the phenolate negative charge through the conjugated system to the positively charged imino functionality as shown in panel **C**. **(D)** shows the conjugate acid/base equilibrium for sinapaldehyde.

**Fig. S17**

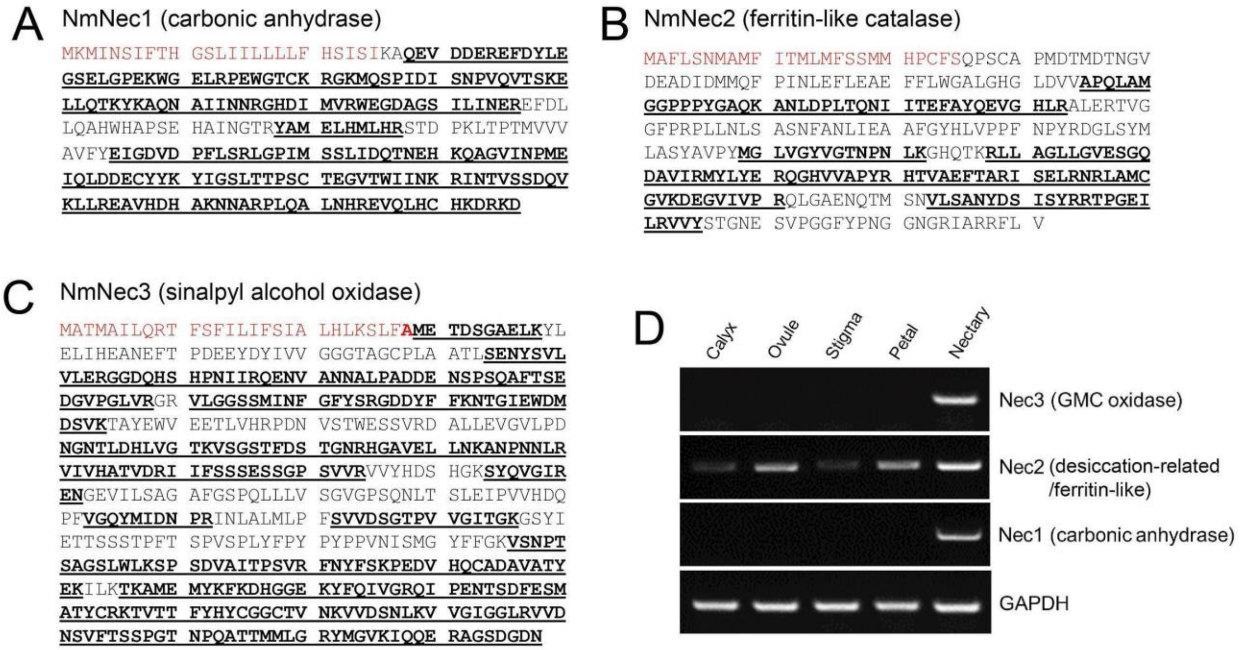


E

| | |
|---|---|
| NmGAPDH RT-F | GGG CTC AAA GCA ATC CCG GCT |
| NmGAPDH RT-R | GGA GCT GCC AAG GCT GTC GG |
| NmNec1 RT-F | CCC ACT GGC ATG CCC CTT CTG |
| NmNec1 RT-R | AGC GCT TGA AGT GGC CTT GCA T |
| NmNec2 RT-F | ACC TCC TCC CTA CGG TGC CC |
| NmNec2 RT-R | CGG ATC ACG GCA TCC TGC CC |
| NmNec3 RT-F | CAG CCG GTT CAC TCT GGC TCA |
| NmNec3 RT-R | TGG TTC CCG GCG AGG ACG TA |

**Fig. S17. Identification of NmNec1, NmNec2 and NmNec3 in *Nesocodon* nectar. (A-C)** The full predicted sequences of **(A)** NmNec1 (accession # pending), **(B)** NmNec2 (accession # pending), and **(C)** NmNec3 (accession # pending) are shown, with bolded and underlined regions corresponding to peptides identified by LC MS/MS analysis. The regions highlighted in yellow corresponds to a predicted signal peptide required for secretion from the cell. **(D)** Expression patterns of Nesocodon nectar proteins by semi-quantitative RT-PCR. Total RNA was isolated from each of the indicated tissues, reverse transcribed, and subjected to 30 cycles of PCR. GAPDH was used as a positive control. **(E)** Primer sequences used for semi-quantitative RT-PCR.

**Fig. S18**

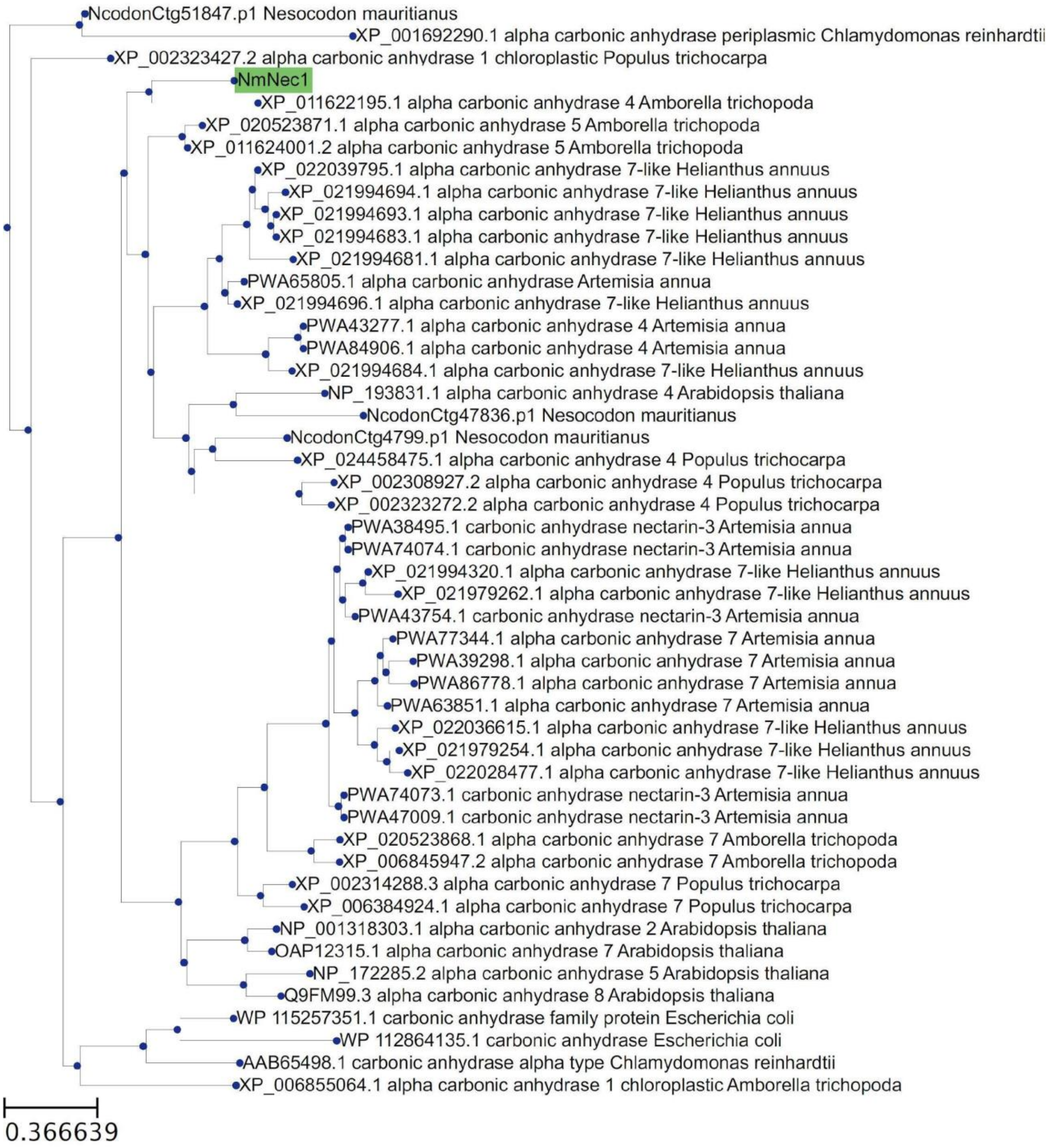


**Fig. S18. Phylogenetic tree of NmNec1 in relation to alpha-type carbonic anhydrases.**

**Fig. S19**

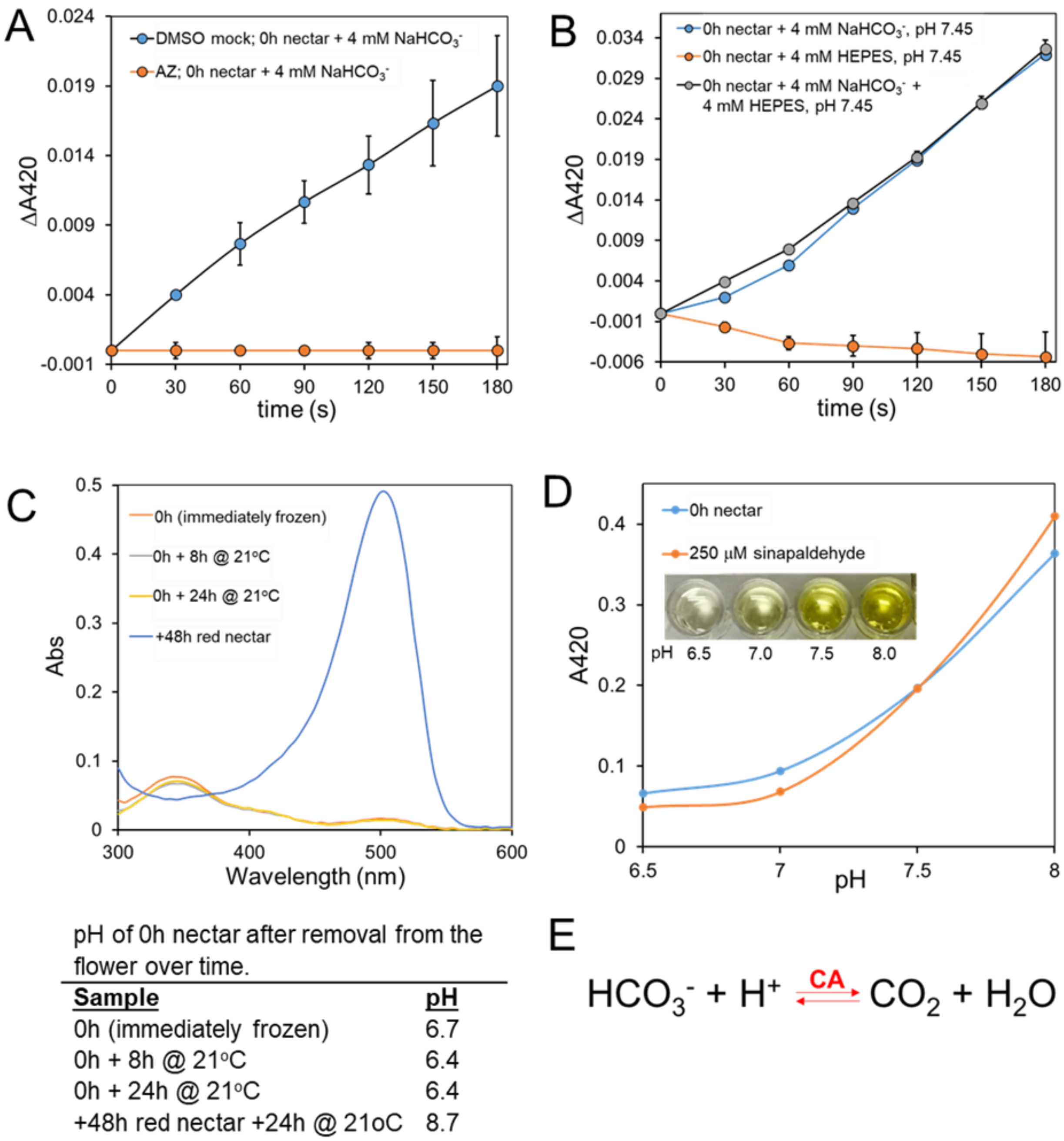


**Fig. S19. Nectar pH can be alkalinized with exogenous application of bicarbonate.**

**(A)** Spectrophotometric assay of carbonic anhydrase activity in 0 h (yellow) nectar. Raw 0 h nectar was first mixed with either acetazolamide (1 mM final; CA inhibitor) or an equivalent amount of 50% DMSO. The reaction was initiated by adding $NaHCO_3$, pH 7.45 at a final concentration of 4 mM, with the absorbance at 420 nm (positively correlating to an increase in pH) being monitored for three minutes. Error bars represent standard deviation, n = 3.

**(B)** In a parallel experiment to that shown in panel A, raw 0 h nectar was mixed at a 1:3 ratio with either $NaHCO_3$, pH 7.45 (blue line) or HEPES, pH 7.45 (orange line; final concentration of 4 mM for both), with the absorbance at 420 nm being monitored for three minutes. Equal mixtures of $NaHCO_3$ and HEPES buffers were used as a control (gray line). Error bars represent standard deviation, n =3.

**(C)** Evaluation of pH of 0 h nectar after removal from the flower over time. Nectar was removed from a newly opened flower (0 h) and 4 µL was immediately frozen at –80˚C, with the remaining sample being stored at 21˚C with the tube lids open in humidified chamber. Additional 4 µL aliquots were removed after 8 h and 24 h incubations and immediately frozen. Absorbance spectra were measured after mixing nectar 1:49 with $H_2O$ and the final pH of eaSFich sample is indicated in the table at right.

**(D)** Validation of endogenous sinapaldehyde as a pH sensor. Shown is an example of the absorbance of 0 h nectar (starting pH of 6.5) at 420 nm as a function of pH. Commercially purified sinapaldehyde (250 µM) has an absorbance peak at 420 nm under alkaline conditions (e.g. Fig. 3A), which also increases with increasing pH. The 0 h nectar (blue line) or purified 500 µM sinapaldehyde (orange line) were mixed 1:1 with 100 mM buffers at a pH of 6.5 (MES), 7.0 (HEPES), 7.5 (HEPES), or 8.0 (HEPES).

**(E)** Overall reaction catalyzed by carbonic anhydrase.

**Fig. S20**

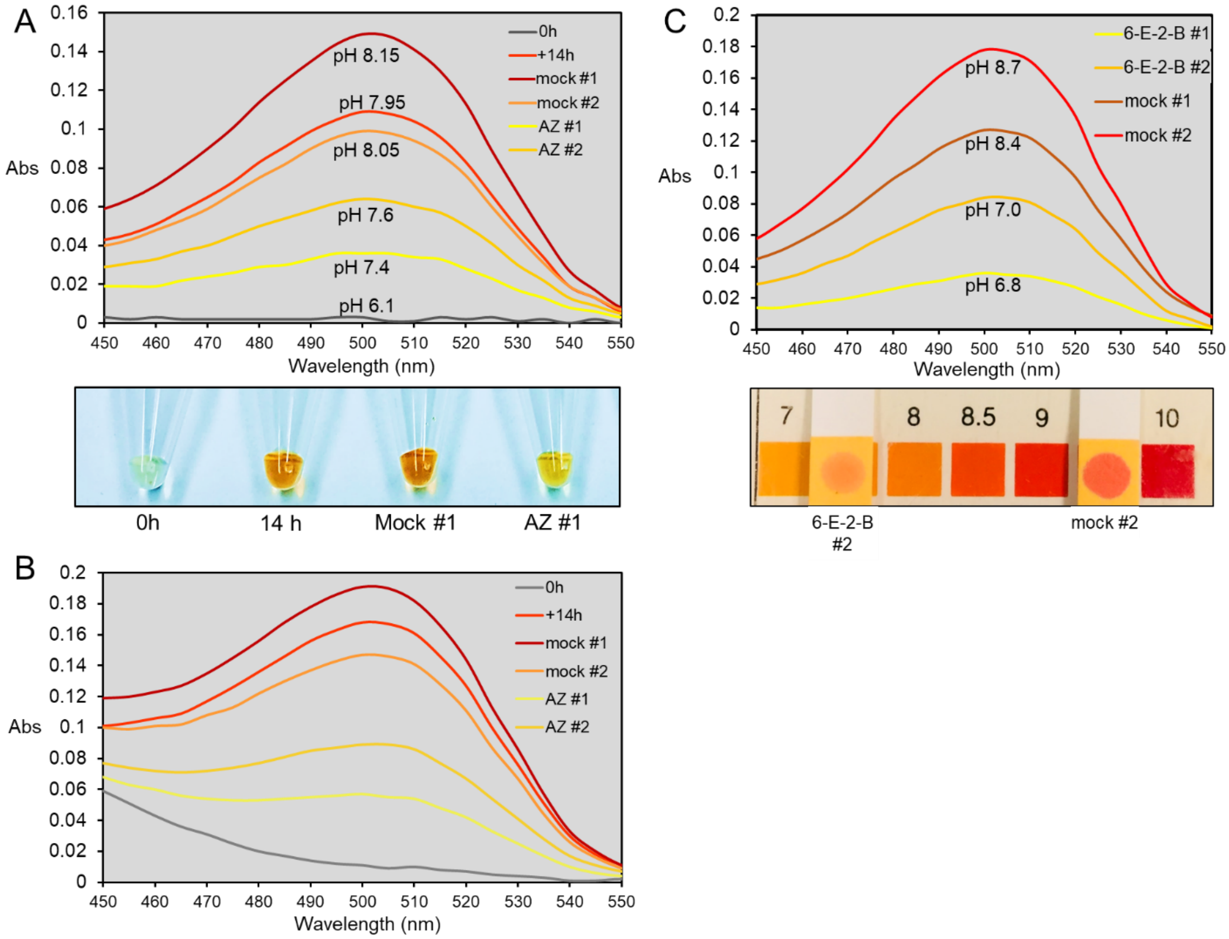


**Fig. S20. Representative additional results demonstrating inhibition of nectar alkalinization with carbonic anhydrase inhibitors.**

**(A, top)** Either acetazolamide (0.5 mM final; AZ #1 and AZ #2) or an equivalent amount of 10% of DMSO (mock #1 and mock #2) were added to 0 h nectar droplets on the same flower and later collected at +14h and directly measured for pH and diluted 3:97 in $H_2O$ prior to measuring absorbance; untreated 0 h and 14h nectar droplets were included for comparison. Images of non-diluted nectars are shown in the bottom panel.

**(B)** Absorbance spectra of nectar samples from **A** after adjustment to pH 8.5 (addition of 25 µl 200 mM Tricine, pH 8.5 directly to 3:97 dilutions from **A**).

**(C)** Individual yellow nectar droplets in a newly opened flower (0 h) were treated either with a final concentration of 0.5 mM 6-ethoxy-2-benzothiazolesulfonamide (6-E-2B; carbonic anhydrase inhibitor) or an equivalent volume of 50% DMSO. Nectar was then collected at +20 h and directly measured for pH and diluted 1:20 in $H_2O$ prior to measuring absorbance. A representative validation of pH is shown in the bottom panel of C.

**Fig. S21**

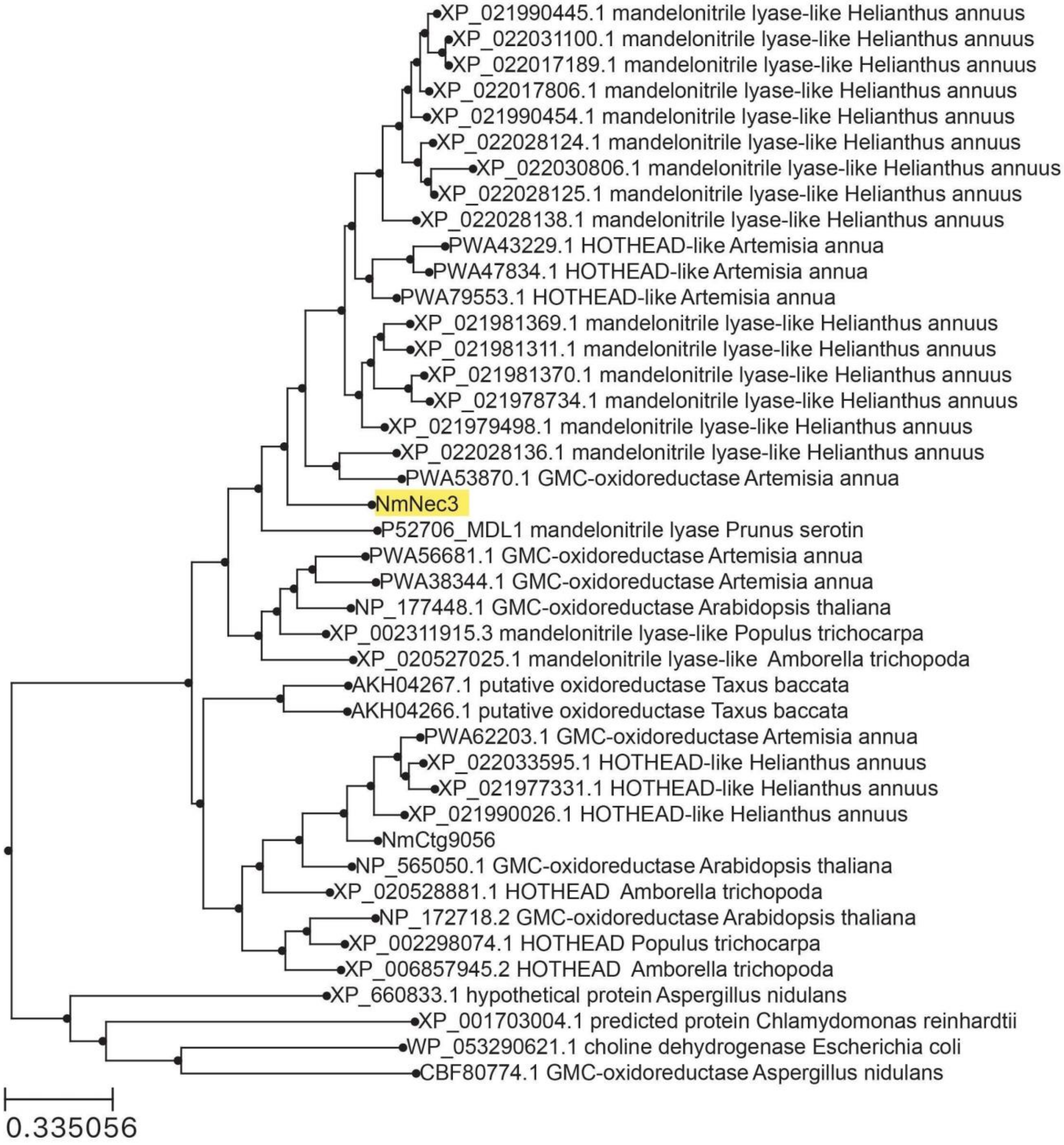


**Fig. S21. Phylogenetic tree of NmNec3 in relation to GMC oxidase superfamily proteins.**

**Fig. S22**

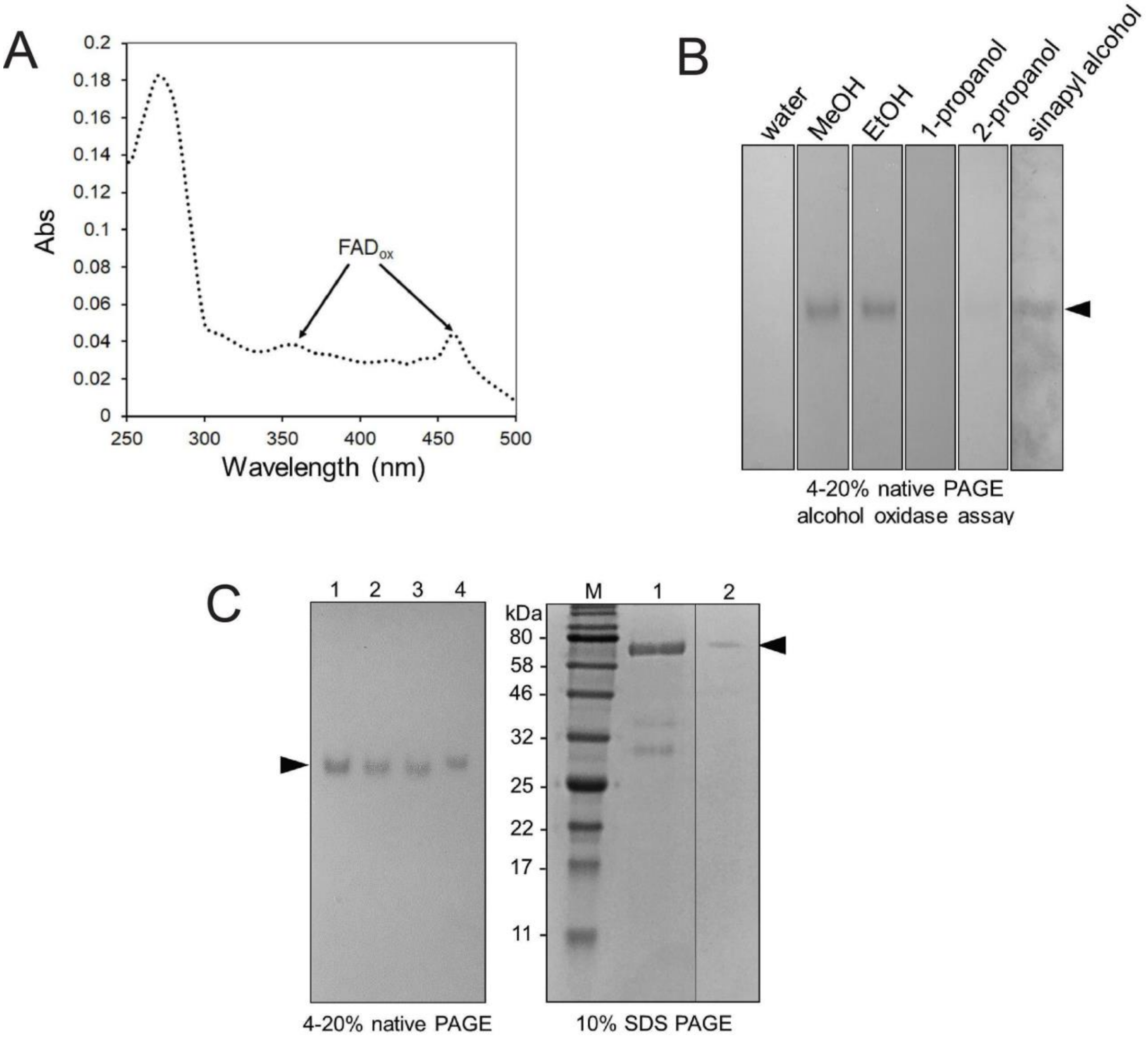


**Fig. S22. Further characterization of NmNec3. (A)** Absorbance spectrum of purified NmNec3, with absorbance peaks corresponding to oxidized FAD indicated. **(B)** In-gel alcohol oxidase activity of NmNec3 with different substrates [10% each of methanol (MeOH), ethanol (EtOH), 1- propanol, 2-propanol, or 5 mM sinapyl alcohol. **(C)** Raw red nectar from four different samples (lanes 1-4; 12.5 μL each on 4-20% gel) were subjected to native PAGE (left panel) of followed by activity staining (as in panel B) with 10% ethanol as the substrate. The activity bands from the gel treated with ethanol (left panel, arrowhead) were excised, incubated with 1 x SDS PAGE loading buffer and then loaded into lane 2 of the right hand 10% denaturing polyacrylamide gel; M = marker (NEB prestained broad), lane 1 contained 45 μL of raw red nectar. Electrophoresis was performed at 150 V for six hours on a 10% denaturing gel and stained with PAGE BLUE (Thermo). The arrowhead on the right panel indicates the location of protein present in the activity band from the activity gel (right panel), which corresponds to NmNec3.

**Fig. S23**

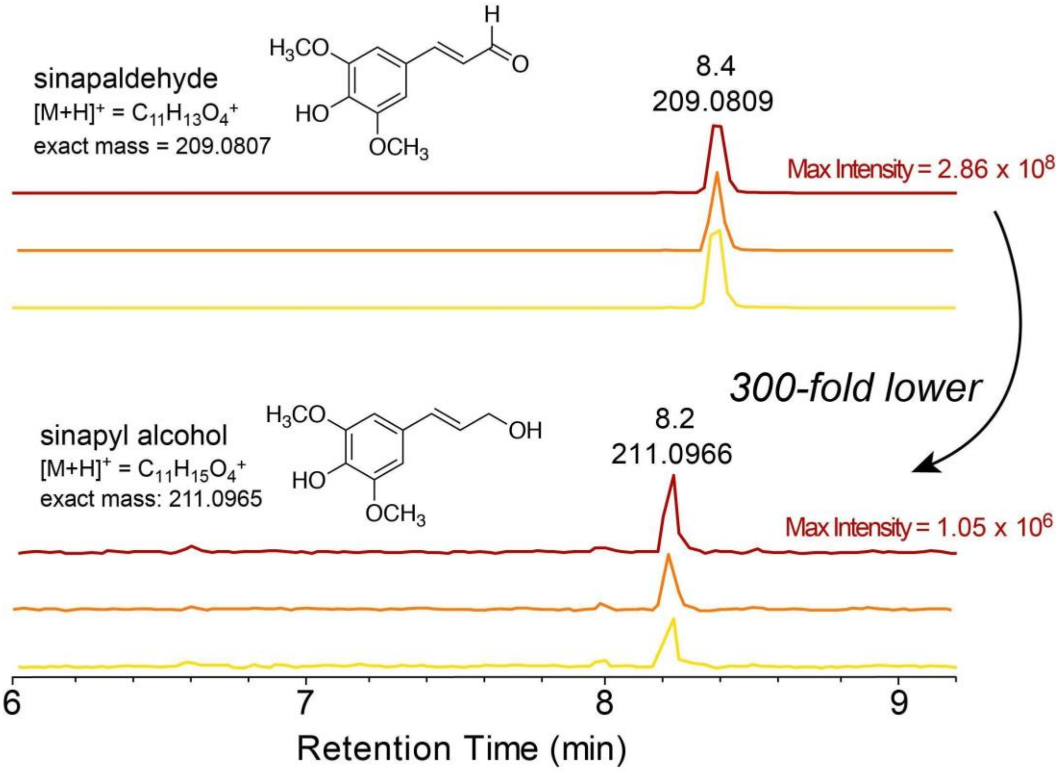


**Fig. S23. Sinapyl alcohol is detectable in yellow, orange, and red *Nesocodon* nectar by LC-MS.**

**Fig. S24**

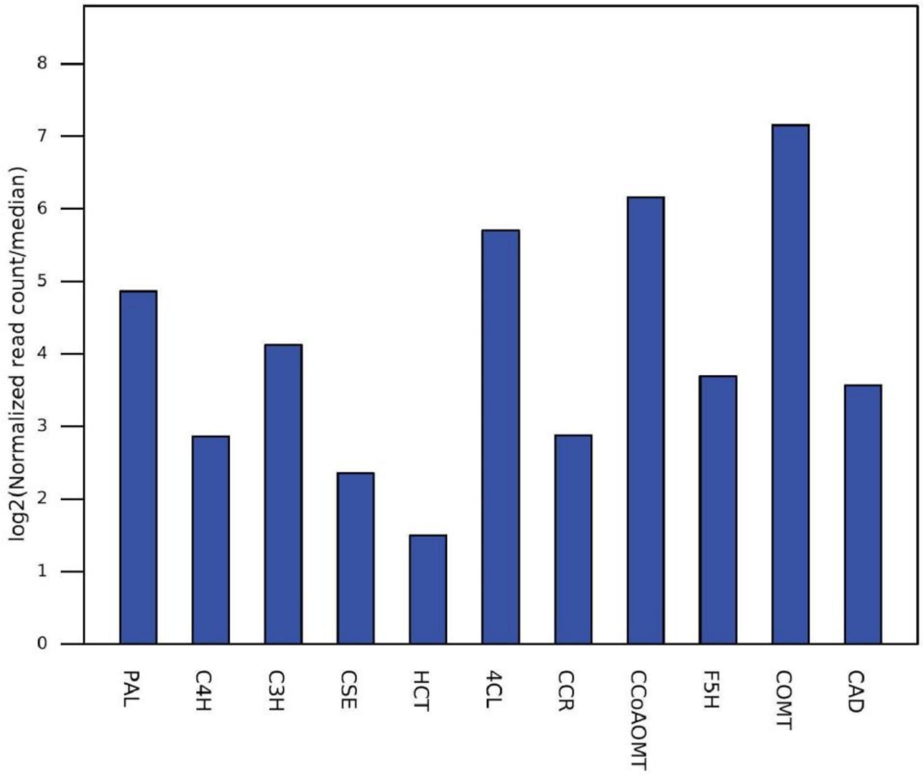


| Enzyme abbreviation | Enzyme description | *H. annuus* top hit | Log base 2 ratio of counts to median |
|---|---|---|---|
| PAL | phenylalanine ammonia-lyase | XM_022168714.1 | 4.86 |
| C4H | trans-cinnamate 4-monooxygenase | XM_022115130.1 | 2.87 |
| C3H | p-coumarate/p-coumaroyl-shikimate/ quinate-3-hydroxylase | XM_022143123.1 | 4.12 |
| CSE | caffeoyl-shikimate esterase | XM_022178021.1 | 2.36 |
| HCT | shikimate O-hydroxycinnamoyltransferase | XM_022162624.1 | 1.5 |
| 4CL | 4-coumarate CoA ligase | XM_022128564.1 | 5.7 |
| CCR | hydroxy cinnamoyl-CoA reductase | XM_022134071.1 | 2.88 |
| CCoAOMT | caffeoyl-CoA O-methyltransferase | XM_022133318.1 | 6.16 |
| F5H | ferulate/coniferaldehyde-5-hydroxylase | XM_022160117.1 | 3.7 |
| COMT | caffeic acid/5-hydroxyconiferaldehyde O-methyltransferase | XM_022179738.1 | 7.16 |
| CAD/SAD | cinnamyl/sinapyl alcohol dehydrogenase | XM_022170114.1 | 3.57 |

**Fig. S24. Transcriptomic analysis of phenylpropanoid pathway genes that lead to sinapyl alcohol.** The log base 2 ratio of the counts for each transcript relative to the median counts for all genes are shown.

**Fig. S25**

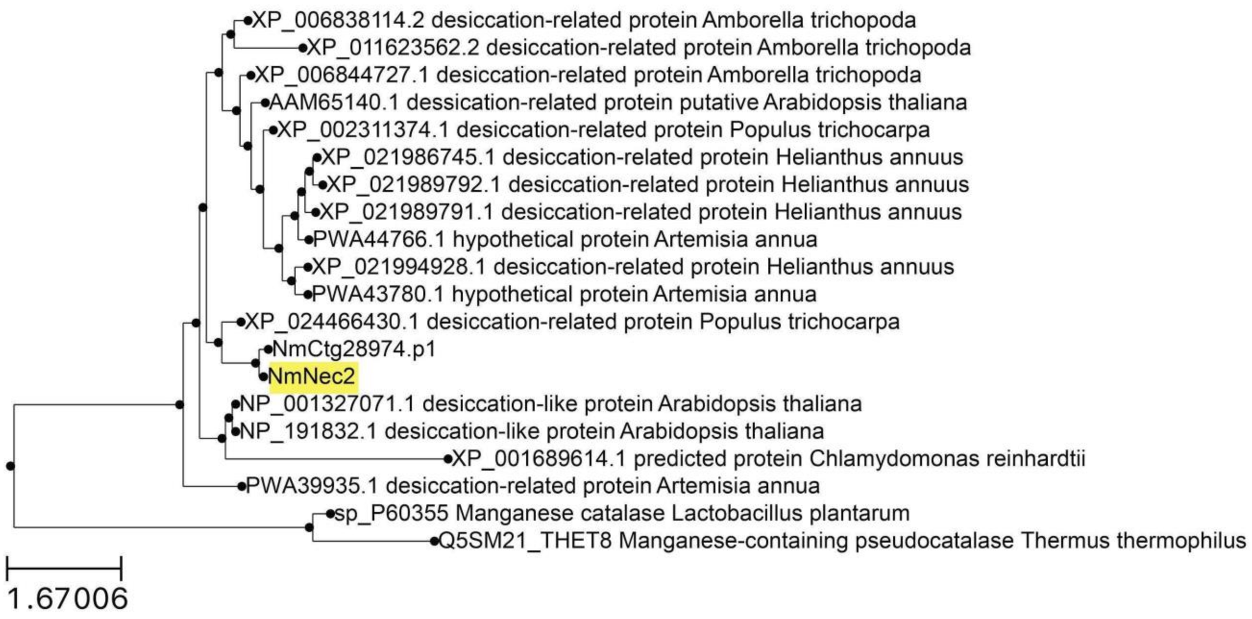


**Fig. S25. Phylogenetic tree of NmNec2 in relation to desiccation-related proteins and bacterial manganese-containing catalases.**

Fig. S26

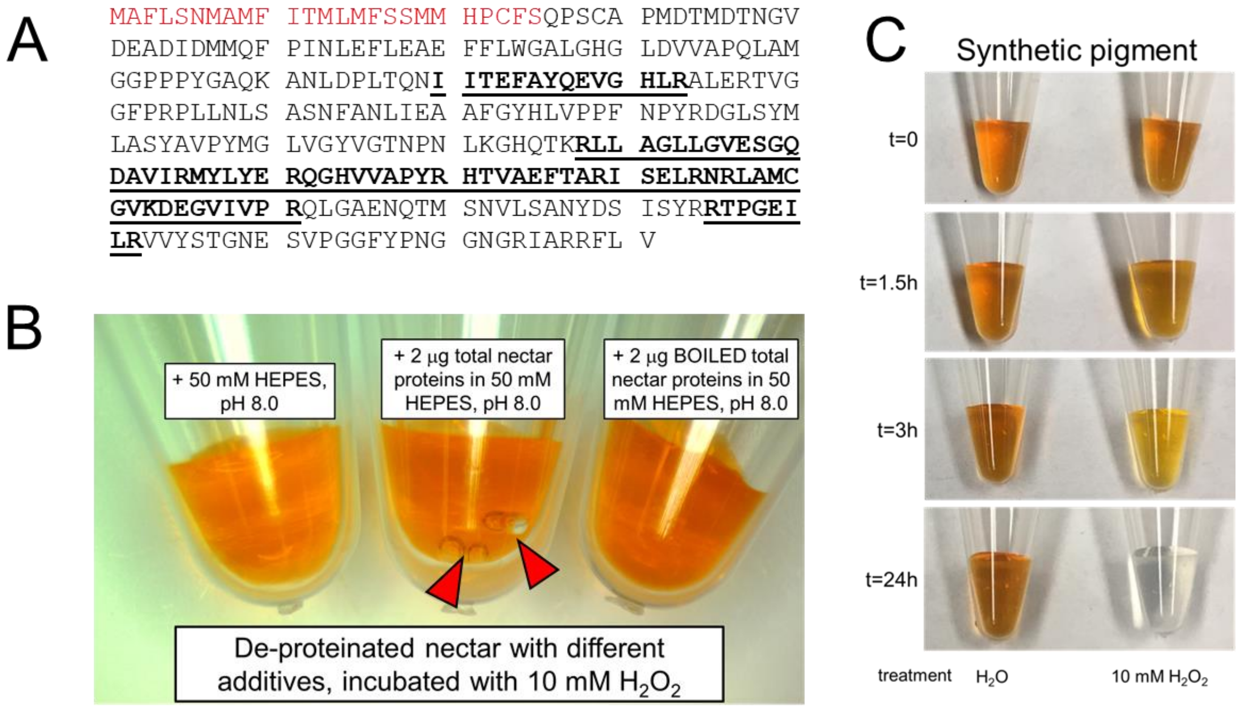


**Fig. S26. Identification of NmNec2 in catalase activity band and further characterization of the impacts of $H_2O_2$ on nectar quality.**

**(A)** Peptides from NmNec2 identified in the catalase activity band from Fig. 4A are bolded and underlined.

**(B)** Oxygen ($O_2$) bubbles (arrowheads) produced by catalase activity from nectar proteins added to de-proteinated nectar.

**(C)** Sensitivity of synthetic nesocodin to exogenous $H_2O_2$.

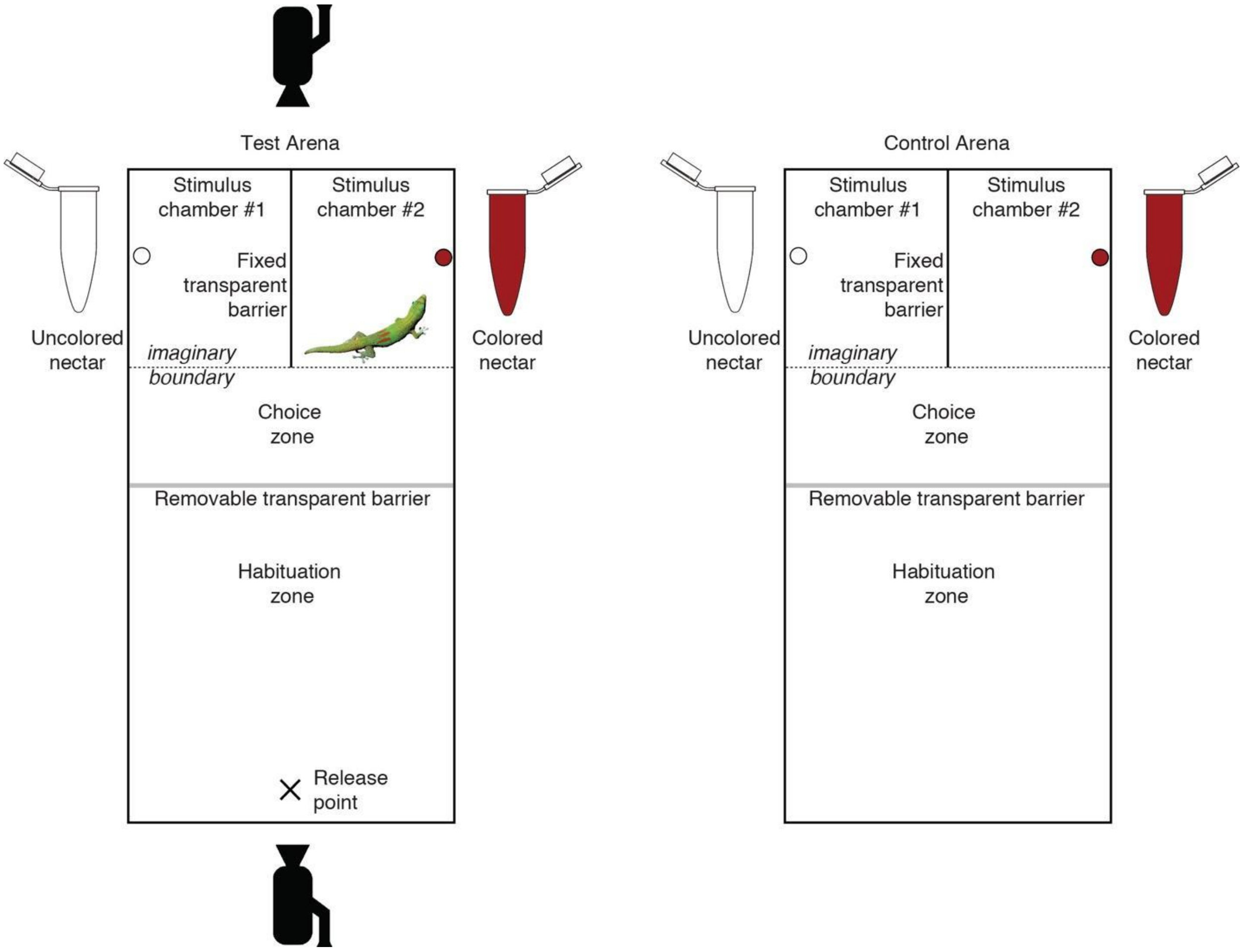


**Fig. S27. Experimental setup for conducting two-alternative choice tests of nectar preferences in gold dust day geckos (*Phelsuma laticauda*).** Two identical arenas (test and control) differed only in the presence/absence of the gecko subject and the associated video cameras for recording behavior. At the beginning of a choice test, the subject was placed at the release point and allowed to explore the habituation zone for 45 min before removal of the removable transparent barrier, which began a 3-hour choice phase during which the subject could explore the entire arena to investigate and sample both sources of nectar stimuli. Gecko image courtesy Thierry Caro (CC BY-SA 3.0; background removed).

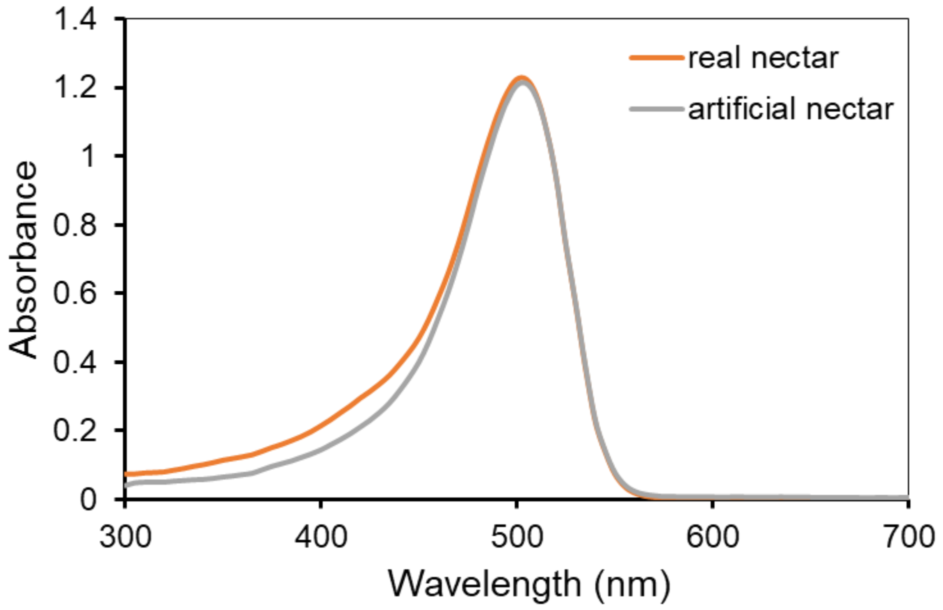


**Fig. S28. Absorbance spectra of synthetic nectar made to match a freshly collected 'real' nectar sample for gecko behavioral assays.** We tested for gecko choice between synthetic red colored versus non-colored nectars. Synthetic nectars, with and without nesocodin (colored and non-colored), contained: 20% sucrose (w/v), 10 mM proline, 1 mM Tricine pH. 8.5, +/- 3 mM nesocodin (pigment), with the final pH being adjusted to 9.0 in order to match the absorbance spectrum of a freshly collected nectar sample.

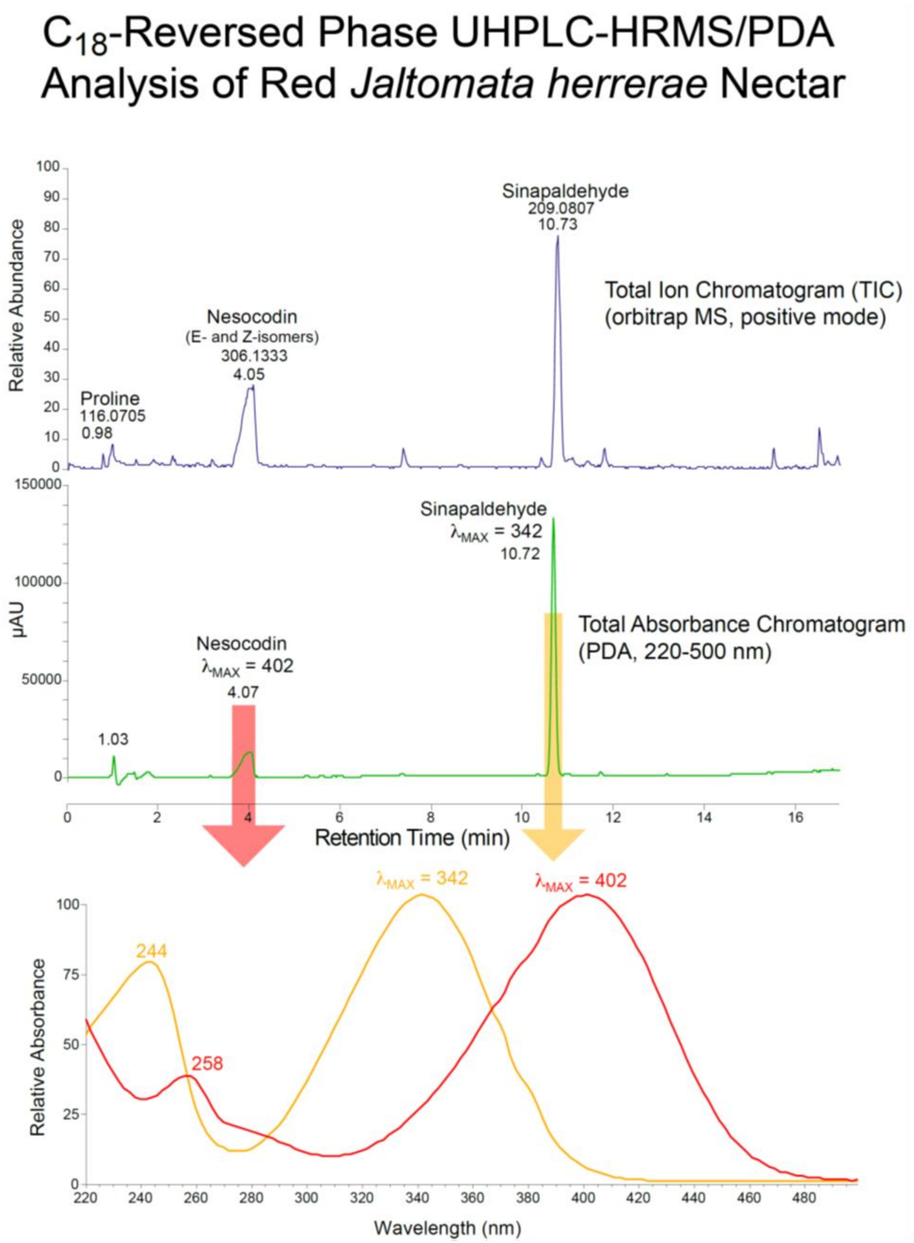


**Fig. S29. LC-MS/PDA analysis of *Jaltomata herrerae* nectar.** Nectar pigment was first purified by solid phase extraction (SPE) using $C_{18}$-Ziptip (Millipore) and then subjected to $C_{18}$-reversed phase chromatography with in line UV/visible spectroscopy (photodiode array, PDA-detector) followed by mass spectrometric analysis (quadrupole-orbitrap hybrid MS). The top panel shows the total ion chromatogram (TIC) for the analysis with major peaks visible for sinapaldehyde @10.72 min and nesocodin at 4.05 min. The second panel shows the total UV/visible absorbance chromatogram, again with the sinapaldehyde and nesocodin as the major peaks. The bottom panel shows the UV/visible spectra for the nesocodin peak (red) with $\lambda_{MAX}$ = 403 nm and the sinapaldehyde peak (yellow) with $\lambda_{MAX}$ = 343 nm. Since the LC solvents contained 0.1% formic acid the pH was low enough that both compounds maintained fully protonated phenolic hydroxyl groups and the spectra are consistent with spectra of red nectar acidified to pH = 6.0 or lower.

```
CLUSTAL O(1.2.4) multiple sequence alignment


JhNec5      --MMNLAAITKILFISLLFLSFAFLARSGEVDDESEFSYDENAKNGPANWGNIHLEWRAC    58
NmNec1      MKMINSI-FTHGSLIILLLLFHSISIKAQEVDDEREFDYLEGSELGPEKWGELRPEWGTC    59
              *:*   :*:  :* **:* .::  :: ***** **.* *.:: ** :**::: ** :*

JhNec5      KNGTMQSPIDLLNERVEVVSNLGILQKYYKPSKATILNRGHDFMLRLD-DGGYLNINGTQ    117
NmNec1      KRGKMQSPIDISNPVQ--VTSKELLQTKYKAQNAIINNRGHDIMVRWEGDAGSILINERE    117
            *.*.******: *     *:.  :**. ** .:* * *****:*:* : *.* : **  :

JhNec5      YQLKQVHWHSPSEHTIDGKRFDMEGHLVHETYDGKK---IVVIAFVFEIGLFPDFFLSII    174
NmNec1      FDLLQAHWHAPSEHAINGTRYAMELHMLHRSTDPKLTPTMVVVAVFYEIGDVDPFL-SRL    176
            ::* *.***:****:*:*.*: ** *::*.: * *    :**:*..:*** .  *: * :

JhNec5      EEDIKAVADKNGAQRAIRIIDPNLIKLDSKKYYRYIGSLTTPPCTEDVVWIIDGKVNTVT    234
NmNec1      GPIMSSLIDQTNEHKQAGVINPMEIQLDDECYYKYIGSLTTPSCTEGVTWIINKRINTVS    236
               :.:: *:.. ::   :*:*  *:**.: **:******** ***.*.***: ::***:

JhNec5      GRQMQLLNDEF----ETNARPVQLLNGRPIKFNKPWPFI-       269
NmNec1      SDQVKLLREAVHDHAKNNARPLQALNHREVQLHCHKDRKD       276
            . *::**.:  .   :.****:* ** * ::::
```

**Figure S30. Multiple sequence alignment between the carbonic anhydrases found in the nectars of *Nesocodon mauritianus* (NmNec1) and *Jaltomata herrerae* (JhNec5)**. These proteins share 42% identity.

```
CLUSTAL O(1.2.4) multiple sequence alignment


NmNec3      ------MAILQRTFSFILIFSIALHLKSLFAMETDSGAELKYLELIHEA----NE-----     45
JhNec7      NLQKKIMTTF-KKLTISLFLTICLSWANY----TSASTDDDFLECLSNQIMNSNSISQVI     55
                  *: : :.::: *:::*.*   .     *.:.:: .:** : :     *.

NmNec3      -FTPDEEYDYIVVGGGTAGCPLAATL--------------------SENYSV-LVLERGG     83
JhNec7      HTRKNSSYSTILNS-FTINQRIRSNLEPSIIITPFNESHIQAAIYCSKIHDVQIRIRSGG     114
                :..*. *: .  * .  : :.*                    *: :.* : :. **

NmNec3      DQHSHPN--------IIRQENVANNALPA----------------------DDENSPSQ     112
JhNec7      HDYEGLSYISETPFVIIDLRNLRSISIDTENKTAWIQSGAILGEVYYRMAEKSKKLAVVA     174
            .::.  .        **  .*: . :: :                       ..: :

NmNec3      AFTSEDGVPGLVRGRVLGGSSMINFGF----------------------YSRGDDYFFKN     150
JhNec7      GFCPTVGVGGLFSGG---GFSLLSRKFGIAADNIIDAKLIDANGQIQDRESMGEDLFWAI     231
            .*    ** **. *    * *::.  *                      * *:* *:

NmNec3      TGIEWDMDSVKTAYEWVEETLVHRPDNVSTWESSVRDALLEVGVLPDNGNTLDHLVGTKV     210
JhNec7      RGGGG--TSFGIIISWKA-KLVDIPEKVTVFNLTR---------------TLEQN----V     269
             *      *.    .*   .**. *::*.:: :                **::     *

NmNec3      SGSTFDSTGNRHGAVELLNKANPNNLRVIVHATVDRIIFSSSESSGPSVVRVVYHDSHGK     270
JhNec7      TQLVYKW---QH----IASKLDE--------NLLLRLILTNSESPFHRGKR-TVHATFSA     313
            :  .:.    :*    : .* :           : *:*::.***      * . * :..

NmNec3      SYQVGIRENGEVILSAGAFGSPQLLLVSGVGPSQNLTSLEIPVVHDQPFVGQYMIDNPRI     330
JhNec7      MFLGGINEL----LREMQKSFPELGLVRE-------DCIEMSLVEANLYI----YGYPRG     358
             :  **.*     *     . *:* **          .:*: :*. : ::     . **

NmNec3      N-LALML--------------PFSVVDSGTPVVGITGKGSYIETTS---S---STPFTS-     368
JhNec7      TALDELLNRKISDIQEGYFKLKSDYVQHPISIDGLEGIWKLMNQVGENSANLMFMPYGGK     418
            . *  :*                  . *:    : *: *  . :: ..   :     *: .

NmNec3      --PVSPLYFPYPYPPVNISMGYF---FGKVSNPTSAGSLWLKSPSDVA---ITPSVRFNY     420
JhNec7      LNDFSESDTPFPHRPGNIFLIHYSVNWGEKE-SSEKHVSWIRELYGYMATYVSKSPRAAY     477
               .*    *:*: * ** : ::   :*: .  :.    *::.  .     :: * *  *

NmNec3      FSKPEDVHQCADAVATYEKILKTKAMEMYKFKDHGGEKYFQIVGRQIPENTSDFESMATY     480
JhNec7      FNYRDL-----------DLGVNNKGNTSYAQARIWGERYFKNNFDRLVQVKTKFDPTNFF     526
            *.  :             :  ::.*.   *      **:**:    :: : .:.*:    :

NmNec3      CRKTVTTFYHYCGGCTVNKVVDSNLKVVGIGGLRVVDNSVFTSSPGTNPQATTMMLGRYM     540
JhNec7      RNE-----------QSIPPLIS--------------------------------------     537
             .:            ::  ::.

NmNec3      GVKIQQERAGSDGDN       555
JhNec7      ---------------       537
```

**Figure 31. Multiple sequence alignment between the alcohol oxidases found in the nectars of *Nesocodon mauritianus* (NmNec3) and *Jaltomata herrerae* (JhNec7)**. These proteins share 21% identity.

**Table S1. Data for gecko visitation.** Gecko ID indicate the unique code of each tested gecko. Sex refers to the sex of each tested animal. Chamber (compartments) visited first correspond to the chamber where the gecko entered first after the removal separator was lifted. Red or Clear refers to the color of the nectar in the tube placed in the chambers. Tube investigation (red or clear) only includes data on gecko licking the tube or gecko touching the tube with the snout (see behavioral data collection above for additional information). Red WG, Clear WG, Red NG, Clear NG refer to the volume recorded after the experiment in the red or clear tube with (WG) or without (NG) the gecko respectively. Diff tube WG and Diff tube NG correspond to the different between the clear and the red nectar tube in the experiment with (WG) or without (NG) the gecko, respectively.

**Video S1.** Example of *Phelsuma laticauda* visiting and consuming synthetic nectar containing nesocodin.

SUPPLEMENTAL REFERENCES CITED


1. Thu YM, *et al.* (2016) Slx5/Slx8 promotes replication stress tlerance by facilitating mitotic progression. *Cell reports* 15(6):1254-1265.
2. Lin-Moshier Y, *et al.* (2013) Re-evaluation of the role of calcium homeostasis endoplasmic reticulum protein (CHERP) in cellular calcium signaling. *J Biol Chem* 288(1):355-367.
3. Ma B, *et al.* (2003) PEAKS: powerful software for peptide de novo sequencing by tandem mass spectrometry. *Rapid communications in mass spectrometry : RCM* 17(20):2337-2342.
4. De Luca V, Del Prete S, Supuran CT, & Capasso C (2015) Protonography, a new technique for the analysis of carbonic anhydrase activity. *Journal of enzyme inhibition and medicinal chemistry* 30(2):277-282.
5. Weydert CJ & Cullen JJ (2010) Measurement of superoxide dismutase, catalase and glutathione peroxidase in cultured cells and tissue. *Nature protocols* 5(1):51-66.
6. Beers RF, Jr. & Sizer IW (1952) A spectrophotometric method for measuring the breakdown of hydrogen peroxide by catalase. *J Biol Chem* 195(1):133-140.
7. Menzel WI, Chen WP, Hegeman AD, & Cohen JD (2012) Qualitative and quantitative screening of amino acids in plant tissues. *Methods in molecular biology* 918:165-178.
8. Grabherr MG, *et al.* (2011) Full-length transcriptome assembly from RNA-Seq data without a reference genome. *Nat Biotechnol* 29(7):644-652.
9. Edgar RC (2004) MUSCLE: multiple sequence alignment with high accuracy and high throughput. *Nucleic acids research* 32(5):1792-1797.
10. Talavera G & Castresana J (2007) Improvement of phylogenies after removing divergent and ambiguously aligned blocks from protein sequence alignments. *Systematic biology* 56(4):564-577.
11. Guindon S, *et al.* (2010) New algorithms and methods to estimate maximum-likelihood phylogenies: assessing the performance of PhyML 3.0. *Systematic biology* 59(3):307-321.
12. Huerta-Cepas J, Serra F, & Bork P (2016) ETE 3: Reconstruction, analysis, and visualization of phylogenomic data. *Molecular biology and evolution* 33(6):1635-1638.
13. Calvino-Cancela M (2005) Phelsuma laticauda laticauda (Golden dust day gecko) Nectarivory. *Herpetological Review* 36(2):182-183.
14. Maia R, Gruson H, Endler JA, & White TE (2018) pavo 2.0: new tools for the spectral and spatial analysis of colour in R. *Methods in Ecology and Evolution* 10:1097-1107.
15. Arden GB & Tansley K (1962) The electroretinogram of a diurnal gecko. *The Journal of general physiology* 45:1145-1161.
16. Taniguchi Y, Hisatomi O, Yoshida M, & Tokunaga F (2001) Pinopsin expressed in the retinal photoreceptors of a diurnal gecko. *FEBS letters* 496(2-3):69-74.
17. Herrera G, *et al.* (2008) Spectral sensitivities of photoreceptors and their role in colour discrimination in the green-backed firecrown hummingbird (Sephanoides sephaniodes). *Journal of comparative physiology. A, Neuroethology, sensory, neural, and behavioral physiology* 194(9):785-794.
18. Stoddard MC, *et al.* (2020) Wild hummingbirds discriminate nonspectral colors. *Proceedings of the National Academy of Sciences of the United States of America* 117(26):15112-15122.

19. Osorio D & Vorobyev M (2008) A review of the evolution of animal colour vision and visual communication signals. *Vision research* 48(20):2042-2051.
20. Endler JA & Mielke PWJ (2005) Comparing entire colour patterns as birds see them. *Biol J Linn Soc* 86:405–431.
21. Amdekar MS & Thaker M (2019) Risk of social colours in an agamid lizard: implications for the evolution of dynamic signals. *Biology letters* 15(5):20190207.
